# Caerulomycin and Collismycin Antibiotics Share a *trans* Flavin-Dependent Assembly Line for 2,2’-Bipyridine Formation and Sulfur Fate Differentiation

**DOI:** 10.1101/2020.12.07.415471

**Authors:** Bo Pang, Rijing Liao, Zhijun Tang, Shengjie Guo, Zhuhua Wu, Wen Liu

## Abstract

Linear nonribosomal peptide synthetases (NRPSs) and polyketide synthases (PKSs) template the modular biosynthesis of numerous nonribosomal peptides, polyketides and their hybrids though assembly line chemistry. This chemistry can be complex and highly varied, and thus challenges the understanding in the diverse polymerization processes of amino acid and carboxylate monomers programmed by various NRPSs and PKSs in nature. Here, we report that caerulomycin and collismycin peptide-polyketide hybrid antibiotics share an unusual assembly line that involves NRPS activity to recruit a flavoprotein acting *in trans* and catalyze C-C bond formation and heterocyclization during 2,2’-bipyridine formation. Simultaneously, this assembly line provides dethiolated and thiolated 2,2’-bipyridine intermediates through differential treatment of the sulfhydryl group arising from L-cysteine incorporation. Subsequent L-leucine extension, which does not contribute any atoms to either caerulomycins or collismycins, plays a key role in sulfur fate determination by selectively advancing one of the two 2,2’-bipyridine intermediates down a path to the final products with or without sulfur decoration. These findings further the appreciation of assembly line chemistry and will facilitate the development of related molecules using synthetic biology approaches.

## INTRODUCTION

Linear nonribosomal peptide synthetases (NRPSs) and polyketide synthases (PKSs) are often large molecular machines that are composed of multidomain modules^1–6^. They have evolved through functional unit (e.g., protein, module or domain) combination and variation to afford various assembly lines that program diverse polymerization and modification processes using amino acids, short carboxylic acids or both as monomers. Assembly line chemistry can be complex and highly varied, and thereby challenges our understanding of how nature creates numerous nonribosomal peptides, polyketides and hybrids thereof that historically play critical roles in medicinal chemistry and chemical biology. The biosynthesis of 2,2’-bipyridine, the core structure of a large class of synthetic bidentate metal-chelating ligands with a variety of applications in many areas of chemistry^7,8^, occurs via such a process.

2,2’-Bipyridine natural products include caerulomycins (e.g., CAE-A, **Fig. 1**), a group of structurally related antibiotics that possess potent immunosuppressive activity^9–13^. While the process through which 2,2’-bipyridine is formed remains poorly understood, we and others unraveled the biogenesis of CAEs and identified a NRPS-PKS assembly line that contains three modular proteins, i.e., CaeA1, CaeA2 and CaeA3, to template a peptide-polyketide hybrid skeleton for 2,2’-bipyridine formation (**Fig. 1**)^14–16^. CaeA1, a bifunctional enzyme composed of a peptidyl carrier protein (PCP) and an adenylation (A) domain, was supposed to prime picolinyl, which provides the unmodified pyridine unit (Ring **B**) of CAEs, as the starter unit for molecular assembly. CaeA2 is a hybrid protein, with a typical PKS module containing a ketosynthase (KS), acyltransferase (AT), and acyl carrier protein (ACP) and an atypical NRPS module containing a condensation/cyclization (Cy) domain, A domain, PCP and terminal C (Ct) domain. Logically, this protein can incorporate malonyl-*S*-CoA and L-cysteine in tandem and associate CaeA1 to provide the essential building atoms for constructing the di- or trisubstituted pyridine unit (Ring **A**) of CAEs^14,17,18^. As proposed, the PKS module of CaeA2 catalyzes two-carbon elongation and installs the atoms C3 and C4 following C2, the carboxyl carbon of the starter picolinyl unit. The atypical NRPS then catalyzes L-cysteine extension, which is unique and occurs at Cβ of the L-cysteine residue to form a C-C bond, instead of it’s α-amino group for amide bond formation as usual, and thus provides C5, C6 and N1 as well as exocyclic C7 after heterocyclization. To our knowledge, peptidyl elongation through a C-C linkage lacks precedent in NRPS catalysis. The domains of CaeA3 are organized as C-A-PCP for L-leucine extension^14,15^. Despite not contributing any atoms in the mature products, CaeA3 activity is necessary for the formation of 2,2’-bipyridinyl-L-leucine (**1**), an off-line intermediate that can be specialized to individual CAE members after L-leucinyl removal^19–23^. If **1** is the direct product of CaeA1, CaeA2 and CaeA3, 2,2’-bipyridine formation could occur in this assembly line through an unprecedented dethiolation-coupled heterocyclization process that involves NRPS-mediated C-C bond formation.

**Figure 1.**
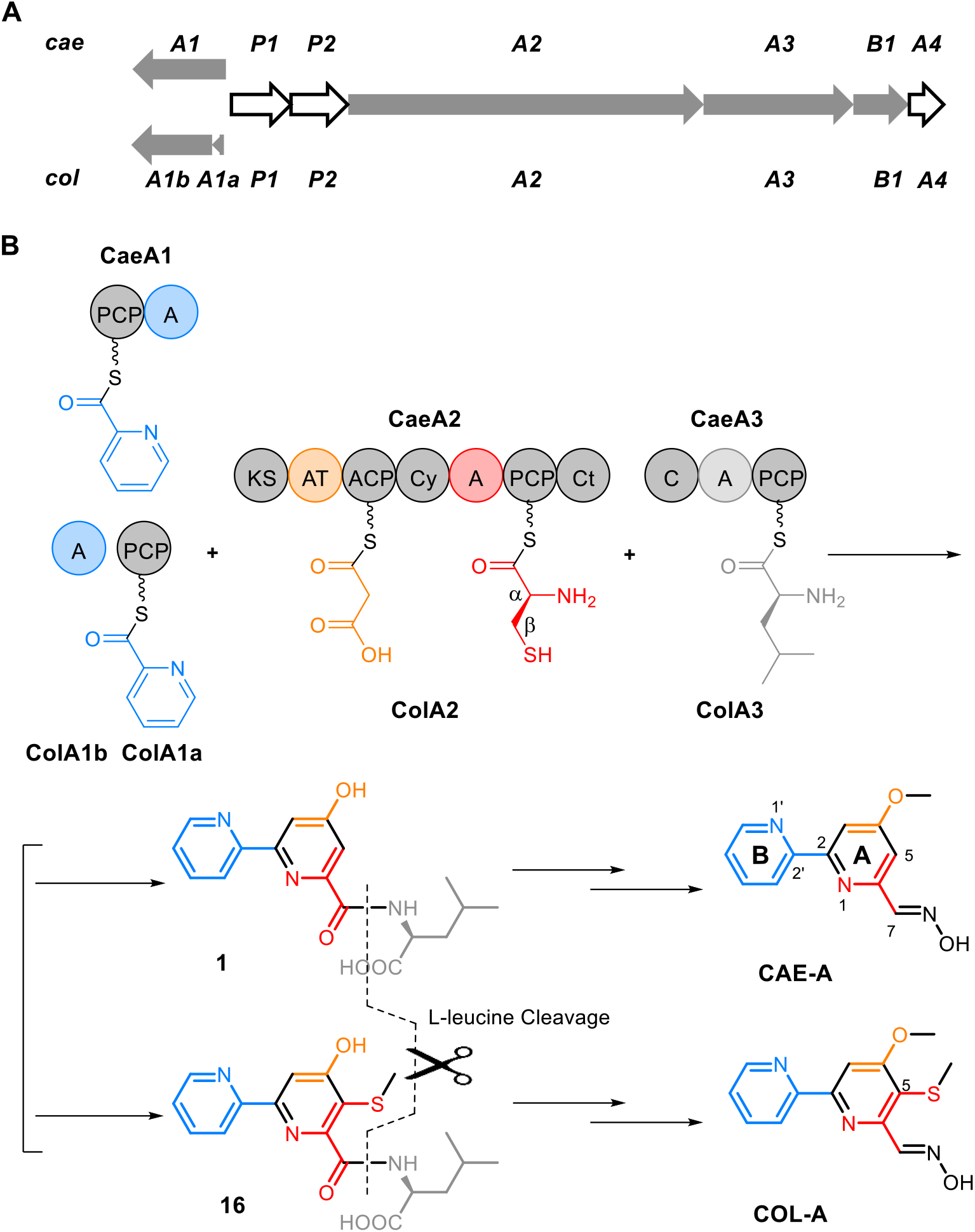
Biogenesis of the polyketide-peptide hybrid, 2,2’-bipyridine antibiotics CAE-A and COL-A. (**a**) Organization of the genes conserved in the CAE and COL biosynthetic gene clusters that codes for the hybrid NRPS/PKS assembly line and its associated *trans* FAD-dependent partner (gray) and other proteins (white). The didomain encoding gene *caeA1* in the *cae* cluster corresponds to the two discrete genes, *colA1a* and *colA1b*, in the *col* cluster. (**b**) Hybrid NRPS/PKS assembly line for 2,2’-bipyridine formation. The functional domains (AT and A domains) for substrate specificity are indicated in different colors with their associated building blocks: blue for picolinyl, yellow for malonyl, red for L-cysteinyl, and light gray for L-leucinyl. The sulfur atoms in the molecules are highlighted in red.

Interestingly, homologs of the above modular synth(et)ases CaeA1, CaeA2 and CaeA3 (i.e., ColA1a and ColA1b, ColA2 and ColA3) were observed in the biosynthetic pathway of collismycins (e.g., COL-A, **Fig. 1**)^14,24,25^, the 2,2’-bipyridine antibiotics differing from CAEs with a sulfur-containing group at C5 of Ring **A**. Most likely, COLs share a similar assembly line for 2,2’-bipyridine formation in which, however, the dethiolation step can be skipped to retain the sulfur-containing group during assembly of the completely identical substrates picolinic acid, malonyl-*S*-CoA, L-cysteine and L-leucine^16^. With great interest in the catalytic logic and the mechanism for sulfur fate determination during 2,2’-bipyridine formation, in this study, we reconstituted the assembly line for the dethiolated antibiotics CAEs, validated the generality of 2,2’-bipyridine formation in the biosynthesis of the thiolated antibiotics COLs and experimentally dissected the on-line process for 2,2’-bipyridine formation and differentiation. The findings reported here exemplify the complexity and variety of assembly line chemistry, which has long been an area of intense interest in the creation of structurally diverse nonribosomal peptides, polyketides and their hybrid molecules.

## RESULTS & DISCUSSION

### 2,2’-Bipyridine assembly line functions with a *trans* flavoprotein

We began by reconstituting the activities of the didomain NRPS CaeA1 (650-aa), the PKS-NRPS hybrid CaeA2 (2484-aa) and the single-module NRPS CaeA3 (1047-aa) *in vitro* (**Fig. 2**). These multifunctional enzymes were individually produced in an engineered *Escherichia coli* strain expressing Sfp^26,27^, a phosphopantetheinyl (Ppant) transferase (PPTase) from *Bacillus subtilis*, leading to the *in vivo* conversion of their PCP and ACP domains from inactive apo-form into active holo-form by the thiolation of each active-site L-serine residue with Ppant. The CaeA1, CaeA2 and CaeA3 proteins produced here were soluble (**Supplementary Fig. 1**), and believed to be active and able to sequentially incorporate the starter and extender units via thiolated PCP/ACP *S*-(amino)acylation for chain elongation (**Supplementary Fig. 2a**). However, no off-line products were observed during the incubation with the substrates picolinic acid, malonyl-*S*-CoA, L-cysteine and L-leucine in the presence of adenosine triphosphate (ATP, necessary for A domain activity) (**Fig. 2a, i**). The CAE-related assembly line likely functions with certain partner(s) that act(s) *in trans*.

**Figure 2.**
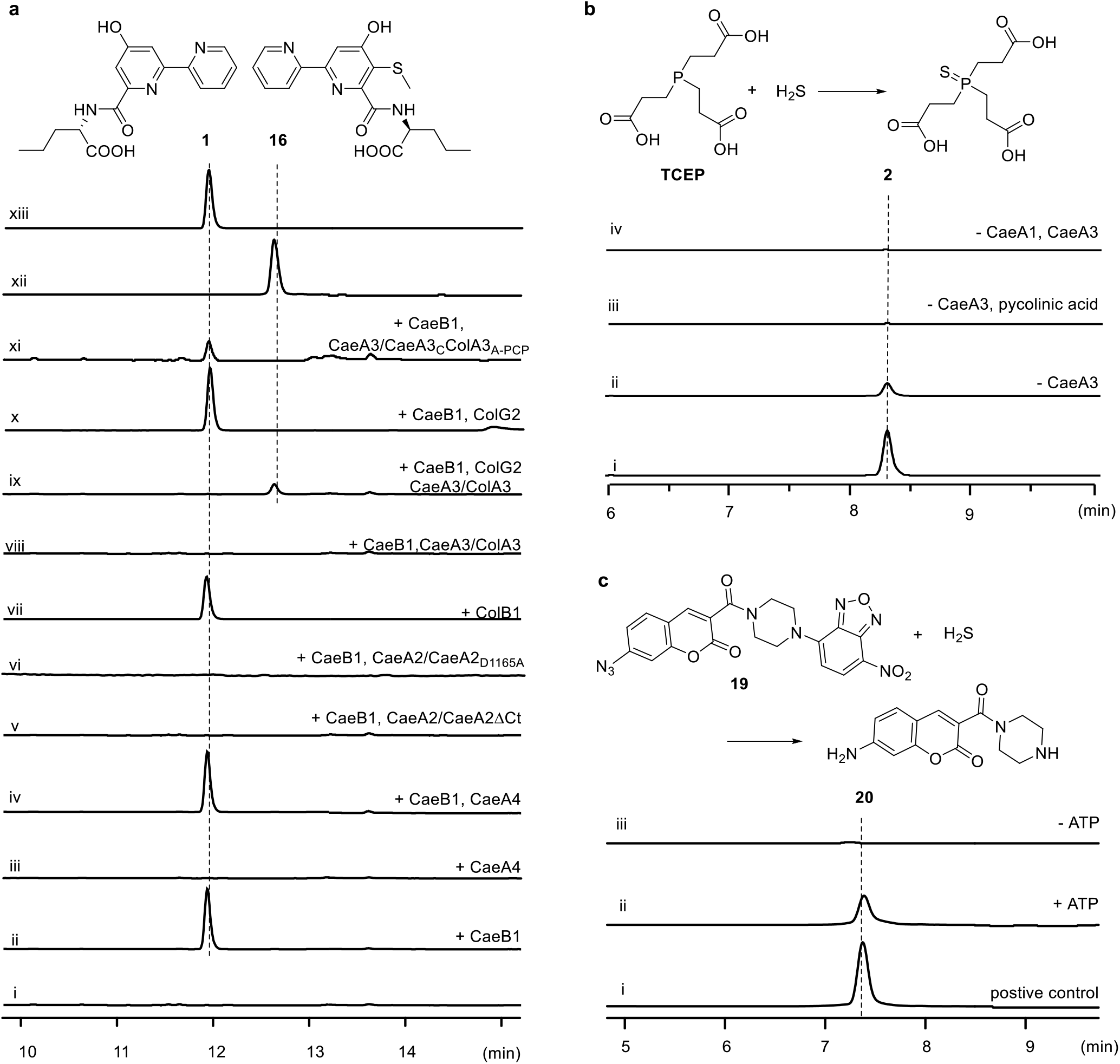
*In vitro* reconstitution of the 2,2’-bipyridine assembly line. (**a**) Analyses of 2,2’-bipyridine production by HPLC (λ 315 nm). The control reaction was conducted with the combination of the proteins CaeA1, CaeA2 and CaeA3 with the substrates picolinic acid, malonyl-*S*-CoA, L-cysteine, L-leucine, and ATP under aerobic conditions (i). To determine the *trans* partner, the reactions were conducted with CaeB1 (ii), CaeA4 (iii), or CaeB1 and CaeA4 (iv). To validate the necessity of Ct, CaeA2 was replaced with CaeA2ΔCt (v). To determine the catalytic role of the Cy domain of the atypical NRPS module, CaeA2 was replaced with CaeA2_D1165A_. (vi).To evaluate the component generality, the reactions were conducted with ColB1 (vii), CaeB1 and ColA3 (replacing CaeA3) (viii), CaeB1, ColA3 (replacing CaeA3) and ColG2 (ix), CaeB1 and ColG2 (x), or CaeB1 and CaeA3_C_ColA3_A-PCP_ (replacing CaeA3) (x). Synthetic **1** (xii) and **16** (xiii) were used as standards. (**b**) Examination of H_2_S production in TCEP-containing reaction mixtures by HPLC-MS. Electrospray ionization (ESI) *m/z* [M + H]^+^ at 283.0405 (calcd.) for **2**, which is generated by a reaction of H_2_S with TCEP. The initial reaction was conducted by combining CaeA1, CaeA2, CaeA3 and CaeB1 with picolinic acid, malonyl-*S*-CoA, L-cysteine, L-leucine, and ATP (i), and reactions in the absence of CaeA3 (ii), picolinic acid and CaeA3 (iii) or CaeA1 and CaeA3 (iv) were also conducted. (**C**) Examination of H_2_S production in TCEP-free reaction mixtures by HPLC with excitation at 370 nm and monitoring emission at 450 nm. The reaction of the fluorescent probe **19** with Na_2_S to yield **20** was used as the positive control (i). H_2_S production was examined in the reactions where CaeA1, CaeA2, CaeA3 and CaeB1 were combined with picolinic acid, malonyl-*S*-CoA, L-cysteine and L-leucine in the presence (ii) and absence (iii) of ATP.

Analyzing the *cae* cluster revealed two genes, *caeB1 and caeA4* (**Fig. 1a**), which encode a flavin-dependent protein and a Type II thioesterase, respectively. Both genes are immediately downstream of *caeA2* and *caeA3* and are likely cotranscripted with these upstream assembly line-encoding genes. To determine whether CaeB1 and CaeA4 are functionally related, we expressed them in *E. coli* (**Supplementary Fig. 1**). Purified CaeB1, which appeared light yellow, was determined to bind oxidized flavin adenine dinucleotide (FAD) in a noncovalent manner based on absorbance spectrum analysis and protein heating followed by high-performance liquid chromatography with high-resolution mass spectrometry (HPLC-HR-MS) (**Supplementary Fig. 3**). The combination of CaeB1 with CaeA1, CaeA2 and CaeA3 resulted in the CAE intermediate 2,2’-bipyridinyl-L-leucine (**1**), whose production however did not require CaeA4-associated, putative thioesterase activity (**Fig. 2a**). Consequently, the CAE NRPS-PKS assembly line requires the *trans* activity of flavoprotein CaeB1 in the assembly of picolinic acid, malonyl-*S*-CoA, L-cysteine and L-leucine. In the reaction mixture containing tris(2-carboxyethyl)phosphine (TCEP) as a reducing agent, we observed tris(2-carboxyethyl)phosphine-sulfide (**2**, **Fig. 2b**), which was indicative of H_2_S production. H_2_S was then trapped using compound **19** as a probe in the TCEP-free reaction mixture (**Fig. 2c**)^28^, confirming the involvement of dethiolation in the production of **1**.

### The *trans* flavoprotein oxidatively processes L-cysteinyl on PCP

Phylogenetically, CaeB1 is close to flavin-dependent dehydrogenases, particularly those acting on acyl thioester substrates (**Supplementary Fig. 3d**)^29,30^. This observation led to the hypothesis that CaeB1 oxidatively processes L-cysteinyl on PCP prior to its noncanonical incorporation into Ring **A** of CAEs. We thus focused on CaeA2 and analyzed its changes in PCP *S*-aminoacylation using a Bottom-Up protein MS strategy^31,32^. For method establishment, thiolated CaeA2 was subjected to protease hydrolysis, and peptide mixtures were analyzed by nanoLC-MS/MS to map the sequence containing the Ppant-modified L-serine residue. Extensive attempts revealed **SLGGD<u>S</u>IMGIQF_2042_VSR** (the relevant L-serine is underlined), a 15-aa sequence generated from complete trypsin treatment and partial chymotrypsin digestion (**Supplementary Fig. 4a**). This sequence can be detected by MS, largely due to its C-terminal alkaline residue, L-arginine, which facilitates peptide ionization. To eliminate the remaining chymotrypsin cleavage site and retain L-arginine, we engineered CaeA2 by mutating F2042 to L-leucine, L-isoleucine and L-valine based on residue conservation analysis (**Supplementary Fig. 2b**). The activity of each variant, i.e., CaeA2^F2042L^, CaeA2^F2042I^ or CaeA2^F2042V^, was then assayed by substituting for wild-type CaeA2 in the **1**-producing reaction mixture. In contrast to the mutations F2042I and F2042V, both of which caused a significant decrease in **1** production, F2042L had little effect on CaeA2 activity (**Supplementary Fig. 2b**). The CaeA2^F2042L^ variant can be completely digested using trypsin and chymotrypsin to efficiently produce **SLGGD<u>S</u>IMGIQL_2042_VSR**, facilitating analysis by HR-MS/MS to confirm the PCP sequence identity and covalently tethered substrate/intermediate (**Supplementary Fig. 4b**).

PCP *S*-aminoacylation-caused mass changes in **SLGGD<u>S</u>IMGIQL_2042_VSR** were then monitored (**Fig. 3**). The incubation of thiolated CaeA2^F2042L^ with L-cysteine yielded L-cysteinyl-*S*-CaeA2^F2042L^ (**3**) (**Supplementary Fig. 5a**), consistent with the activities of the A and PCP domains of the CaeA2 NRPS module. The presence of L-cysteinyl in CaeA2^F2042L^ was confirmed by treatment with iodoacetamide (IAM, a thiol derivatization reagent), which yielded derivative **4** (with an MS signal ∼30-fold stronger than that of **3**, **Fig. 3** (ii, left) and **Supplementary Fig. 5b**). Incubation of the above mixture with CaeB1 significantly lowered the production of **3** by ∼70% (**Fig. 3**, iii, left). Accordingly, treatment with IAM generated **5**, a derivative of (3-sulfhydryl)-pyruvoyl-*S*-CaeA2^F2042L^ (**6**) (**Fig. 3** (iii, middle) and **Supplementary Fig. 6a**), indicating that CaeB1 acts on PCP for L-cysteinyl transformation. This conclusion was supported by the observation of a + 3Da derivative of **5** when L-[1,2,3-^13^C_3_,^15^N]cysteine was used (**Supplementary Fig. 6**). Using L-[2,3,3-D_3_]cysteine as the substrate resulted in deuterium-unlabeled **5** (**Supplementary Fig. 6**), consistent with the notion that CaeB1 catalyzes the α,β-dehydrogenation of L-cysteinyl and converts **3** to dehydrocysteinyl-*S*-CaeA2^F2042L^ (**7**), which appears to be unstable and readily undergoes epimerization followed by hydrolysis to produce **6**. Proton exchange can occur at the Cβ position of **6**, leading to a loss of deuterium.

**Figure 3.**
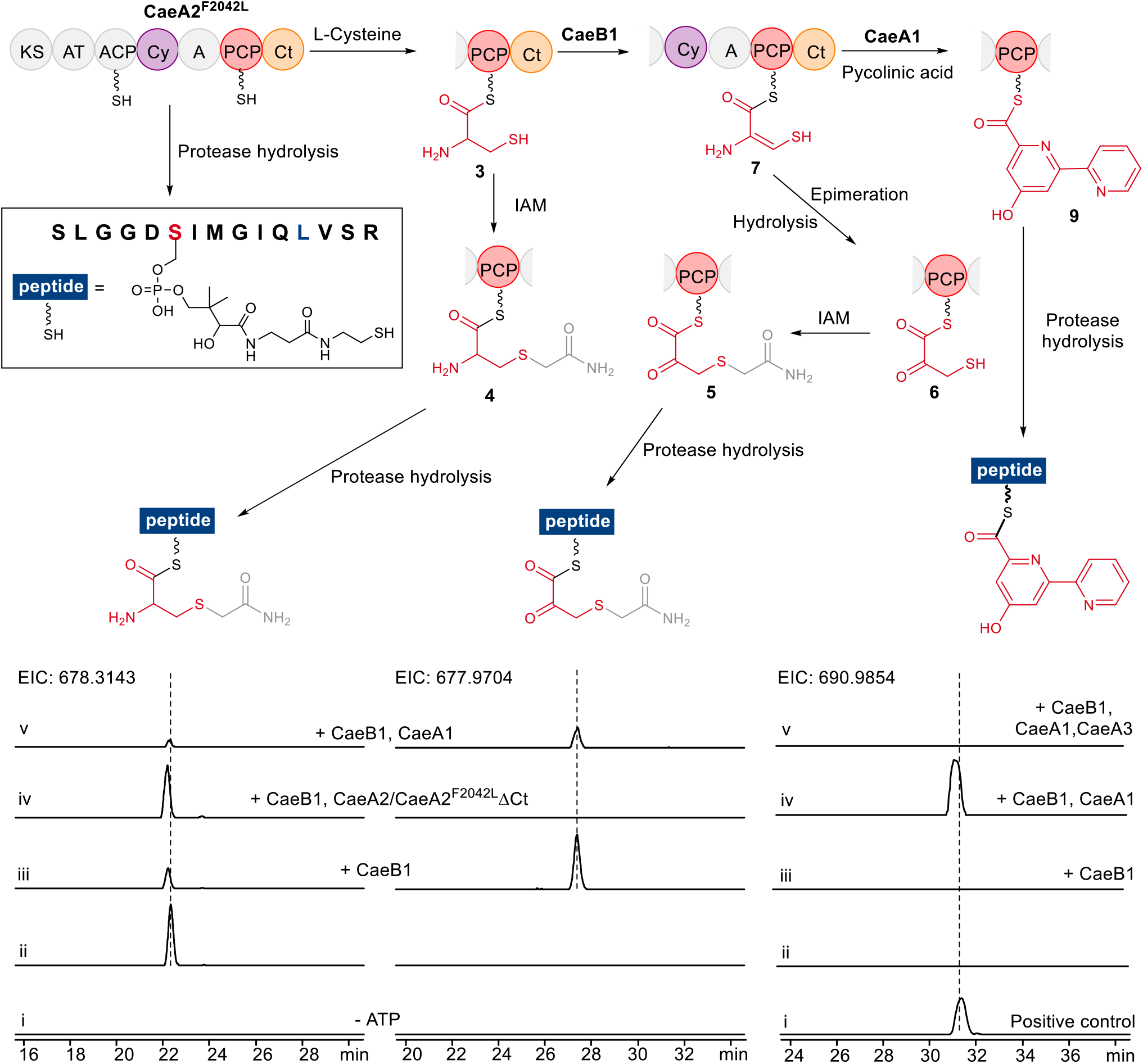
Examination of PCP-tethered intermediates during 2,2’-bipyridine formation (top) by nanoLC-MS/MS (below). The sequence **SLGGD<u>S</u>IMGIQL_2042_VSR** (in the rectangle) arises from the complete digestion of CaeA2^F2042L^ using trypsin and chymotrypsin. For details of the HR-MS/MS analyses, see **Figs. S4**∼**S6**, **S9** and **S11**. The two IAM-treated sequences (ESI *m/z* [M + 3H]^3+^ for the left, calcd. 678.3143; for the middle, calcd. 677.9704) come from L-cysteinyl-*S*-CaeA2^F2042L^ (**3**) and dehydrocysteinyl-*S*-CaeA2^F2042L^ (**7**), respectively, which were examined by incubating CaeA2^F2042L^ and L-cysteine in the absence (i, negative control) and presence (ii) of ATP, with CaeB1 (iii), CaeB1 and CaeA2^F2042L^ΔCt (replacing CaeA2^F2042L^) (iv), or CaeB1, CaeA1, picolinic acid and malonyl-*S*-CoA (v). The right sequence (ESI *m/z* [M + H]^+^, calcd. 690.9854) comes from 2,2’-bipyridinyl-*S*-CaeA2^F2042L^ (**9**). The positive control was prepared by incubating CaeA2^F2042L^ (with PCP in apo form) and 2,2’-bipyridinyl-*S*-CoA (i). **9** was examined by incubating CaeA2^F2042L^ and L-cysteine (ii), with CaeB1 (iii), CaeB1, CaeA1, picolinic acid and malonyl-*S*-CoA (iv), or CaeB1, CaeA1, CaeA3, picolinic acid, malonyl-*S*-CoA and L-leucine (v).

The sequence alignment of CaeB1 with various flavin-dependent dehydrogenases revealed potential key residues including E372 at the active site and S168 related to FAD binding (**Supplementary Fig. 3d**). These residues were mutated, generating CaeB1^E372L^ and CaeB1^S168A^. The former variant shares with the wild-type protein a light yellow color and an absorbance spectrum characteristic of FAD in oxidized form; in contrast, the latter variant was nearly colorless, indicating the loss of FAD-binding ability (**Supplementary Fig. 3**). CaeB1^E372L^ and CaeB1^S168A^ were individually used to replace wild-type CaeB1 in the **1**-producing reaction mixture. Both mutations abolished **1** production, and saturation with an excess of FAD compensated the mutation S168A only (**Supplementary Fig. 3**). These results ascertained that CaeB1 activity is FAD-dependent. Mechanistically, CaeB1 catalyzes α,β-dehydrogenation likely utilizing E372 as a base for L-cysteinyl α-deprotonation, followed by the elimination of a β-hydride equivalent to FAD to yield dehydrocysteinyl and FADH_2_ (**Supplementary Fig. 7**).

### The atypical NRPS Ct domain recruits *trans* flavoprotein activity

Notably, CaeB1 activity strictly depends on the Ct domain of CaeA2, a C-like domain that is shortened by ∼1/3 and lacks the catalytic L-histidine residue (**Supplementary Fig. 8**). Deleting this domain completely abolished the production of **1** (**Fig. 2a, v**). In the incubation of truncated CaeA2^F2042L^ΔCt with L-cysteine, L-cysteinyl-*S*-CaeA2^F2042L^ΔCt (**8**) was observed; however, the α,β-dehydrogenation of L-cysteinyl failed to occur when CaeB1 was introduced (**Fig. 3** (iv, left and middle) and **Supplementary Fig. 9**). We thus proposed that resembling the X domain of the NRPSs involved in glycopeptide biosynthesis^33^, Ct is necessary for recruiting the activity of the flavoprotein CaeB1 that functions *in trans*. To validate this hypothesis, we expressed the N-terminally thioredoxin (Trx)-tagged Ct domain in *E. coli* and measured its interaction with CaeB1 by isothermal titration calorimetry (ITC). Because CaeB1 is relatively unstable and tends to be precipitated during the measurement, we prepared the variant of this flavoprotein by fusion with maltose binding protein (MBP) at the N-terminus. ITC analysis revealed a *K*_d_ value of 0.53 ± 0.12 μM between MBP-fused CaeB1 variant and Trx-tagged Ct domain, indicating the existence of a moderate interaction (**Fig. 4**). In contrast, this interaction was not observed by titrating MBP-fused CaeB1 variant to control proteins, e.g., CaeA2ΔCt (**Supplementary Fig. 10a**). Consequently, the flavoprotein CaeB1 likely functions *in trans* through the interaction with the Ct domain of CaeA2, leaving the Cy domain in the atypical NRPS module as the catalyst for dehydrocysteinyl extension and subsequent heterocyclization.

**Figure 4.**
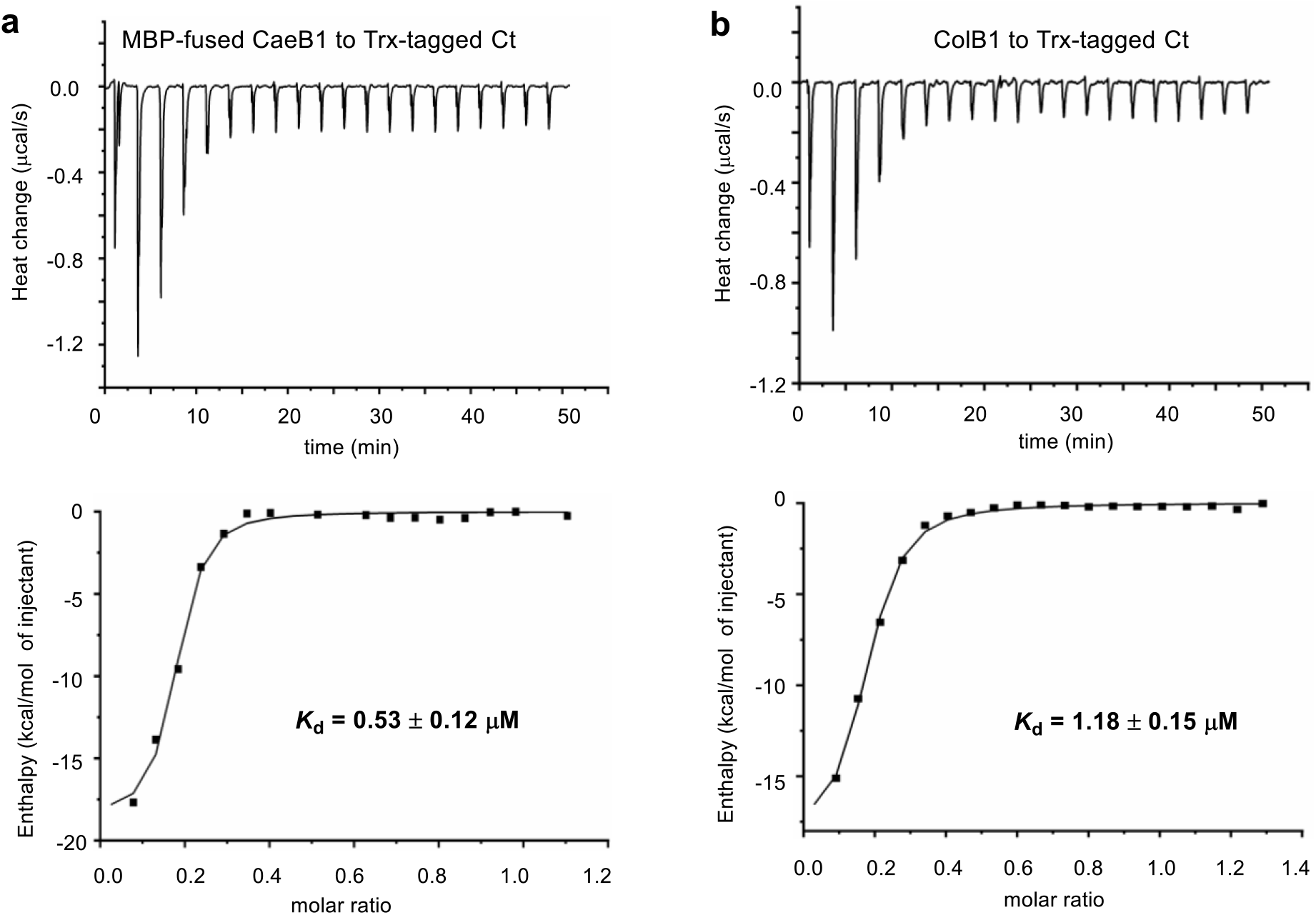
Measurement of the interactions of the Ct domain of CaeA2 with related flavoproteins by ITC. Raw dates were shown on top, and the integrated curves containing experimental points and the best fitting line obtained from the single binding site model were shown on bottom. This measurement was conducted in triplicate. For negative controls, see **Supplementary Fig. 10**. (**a**) Titrating MBP-fused CaeB1 to Trx-tagged Ct. (**b**) Titrating ColB1 to Trx-tagged Ct.

### 2,2’-Bipyridine formation proceeds in the atypical NRPS module

We then examined the elongating intermediate covalently tethered to the PCP domain of CaeA2 during **1** formation. The incubation of CaeA1, CaeA2^F2042L^ and CaeB1 with picolinic acid, malonyl-*S*-CoA and L-cysteine led to the accumulation of 2,2’-bipyridinyl-*S*-CaeA2^F2042L^ (**9**). In contrast, **9** was not observed in the presence of CaeA3 for **1** production (**Fig. 3** (iv and v, right) and **Supplementary Fig. 11a**), supporting that 2,2’-bipyridinyl is an intermediate formed in the atypical NRPS module of CaeA2 before L-leucine extension. The atypical NRPS Cy domain likely catalyzes the formation of 2,2’-bipyridinyl, because mutating the catalytic residue D1165 in its conserved motif DxxxxD(1165)xxS to L-alanine completely abolished the production of **1** (**Fig. 2a, vi**). H_2_S was observed during **9** formation (**Fig. 2b**), indicating that dethiolation occurred. The production of 2,2’-bipyridinyl in the atypical NRPS module of CaeA2 was further confirmed by HR-MS/MS analysis of the positive control of 2,2’-bipyridinyl-*S*-CaeA2^F2042L^, which was generated by incubating unthiolated CaeA2^F2042L^ with synthetic 2,2’-bipyridinyl-*S*-CoA in the presence of Sfp (**Fig. 3** (i, right) and **Supplementary Fig. 11b**). In addition, the combination of CaeA3 with truncated 2,2’-bipyridinyl-*S*-PCP_CaeA2_ (**10**, produced by thiolating the apo-form PCP_CaeA2_ domain with Sfp in the presence of 2,2’-bipyridinyl-*S*-CoA) and L-leucine produced **1** (**Supplementary Fig. 12**), validating that this single module NRPS accepts dethiolated 2,2’-bipyridinyl for L-leucine extension. Notably, 2,2’-bipyridinyl loses its chirality at Cα. This feature is consistent with the fact that CaeA3 possesses a ^D^C_L_ domain (**Supplementary Fig. 13**)^34^. A similar catalysis was observed in the biosynthesis of nocardicins^35^, in which an NRPS ^D^C_L_ domain mediates the assembly of a Cα−achiral intermediate with an L-amino acid for β-lactam formation.

### Timing of the dethiolation step in the formation of the CAE 2,2’-bipyridine core

Focusing on the timing of the dethiolation step, we attempted to dissect the specific 2,2’-bipyridine-forming process in the CAE assembly line (**Fig. 5**). Omitting CaeA1 for picolinic acid incorporation from the **1** or **9**-forming reaction mixture prevented H_2_S production (**Fig. 2b**), inconsistent with *route a* in which condensation follows the dethiolation of dehydrocysteinyl to 2-amino-allyl-*S*-CaeA2 (**11**). In addition, derivatives of the possible intermediate **11** (e.g., pyruvoyl-*S*-CaeA2, which might arise from epimerization and hydrolysis in a manner similar to the conversion of **7** to **6**) in this route were not observed based on careful HR-MS/MS analysis. Removing the substrate picolinic acid from the **1** or **9**-forming reaction mixtures led to similar results, supporting the hypothesis that the condensation with dehydrocysteinyl precedes its dethiolation. The Cy domain of CaeA2 shares high sequence homology to the counterparts known for thiazoline formation (**Supplementary Figs. 8** and **14**), in which related Cy domains display dual activity for L-cysteine or L-serine extension by forming an amide bond first and then cyclization via the nucleophilic addition of the –SH or –OH side chain of the newly incorporated residue onto the preceding carbonyl group to yield a five-membered thiazoline or azoline ring after H_2_O elimination^3,5^. However, the Cy domain of CaeA2 appears to be functionally distinct because it does not utilize unprocessed L-cysteinyl for condensation (**Figs. 2** and **3**). This domain likely follows CaeB1 activity and condenses the resulting reactive dehydrocysteinyl unit with the upstream intermediate for C-C bond formation through unprecedented Cβ-nucleophilic substitution, yielding picolinyl-acetyl-dehydrocysteinyl-*S*-CaeA2 (**12**, **Fig. 5**) before heterocyclization to form the six-membered pyridine ring. We then utilized L-[2,3,3-D_3_]cysteine in the **1**-forming reaction mixture. This assay led to the production of the + 1Da derivative of **1** (**Supplementary Fig. 15**), which excludes *route b* in which cyclization precedes dethiolation. As shown in this route, cyclization requires the epimerization of **12** to the enamine, which would eliminate the Cβ deuterium of the dehydrocysteinyl residue and result in unlabeled **1**. Instead, *route c* is favored based on this observation because the dethiolation of **12** to picolinyl-acetyl-dehydroalanyl-*S*-CaeA2 (**13**) facilitates cyclization to afford 2,2’-bipyridinyl and retain the Cβ deuterium (**Fig. 5**). For dethiolation, reduced FADH_2_ might serve as a hydride supplier and undergo oxidation to produce FAD for recycling, consistent with a net non-redox result from this process (**Supplementary Fig. 7a**).

**Figure 5.**
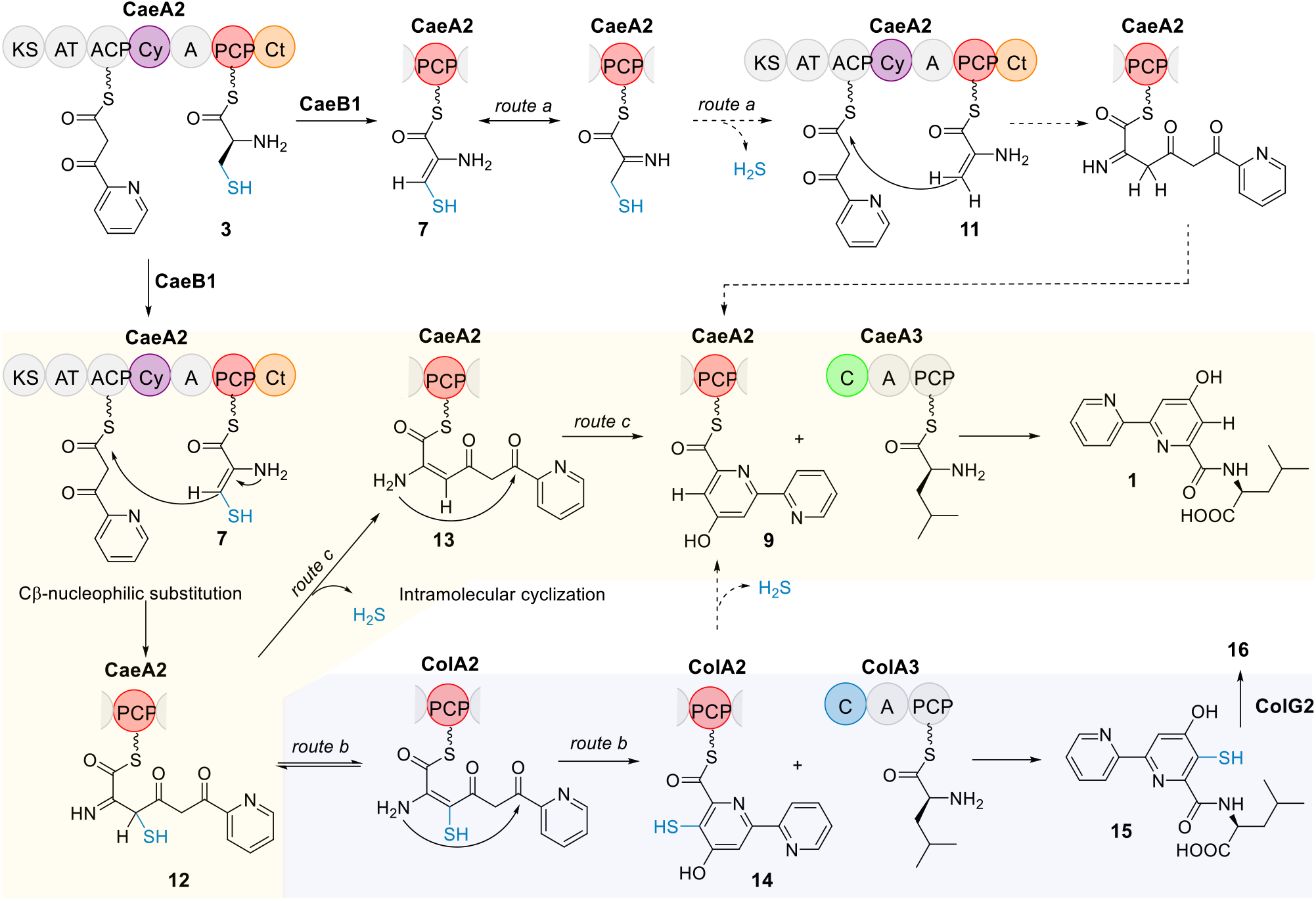
Proposed mechanisms for 2,2’-bipyridine formation. The PCP domain of CaeA2 is labeled in red. The C domains of CaeA3 and ColA3 are highlighted in green and blue, respectively. The favored pathways for CAEs and COLs are highlighted with light yellow and light purple backgrounds, respectively.

### 2,2’-Bipyridine assembly line provides both dethiolated and thiolated intermediates

Intriguingly, although *route b* is not favored in CAE biosynthesis, skipping the dethiolation step allows the condensation of 5-sulfhydryl-2,2’-bipyridinyl-*S*-CaeA2 (**14**) with L-leucine to produce 5-sulfhydryl-2,2’-bipyridinyl-L-leucine (**15**), the proposed off-loading intermediate in the biosynthesis of COLs (**Fig. 5**). Distinct from CAEs, COLs are 2,2’-bipyridine antibiotics possessing a sulfur-containing group (**Fig. 1**). In addition to ColA1a and ColA1b, ColA2 and ColA3, which share high sequence homology with the CAE-related modular synth(et)ases CaeA1, CaeA2 and CaeA3, respectively, the biosynthetic pathway of COLs involves ColB1, the homolog of the flavoprotein CaeB1 that also noncovalently binds a FAD cofactor (**Supplementary Fig. 3**), and thus are assumed to share a similar *trans* flavin-dependent NRPS-PKS assembly line for the formation of the different thiolated 2,2’-bipyridine core^14,25^. To determine the difference in 2,2’-bipyridine formation, we attempted to replace the CAE enzymes with their counterparts from the COL pathway in the **1**-producing reaction mixture. These counterparts did not include those for picolinyl priming (i.e., CaeA1 or ColA1a/ColA1b) and subsequent two-carbon elongation (i.e., the PKS module of CaeA2 or ColA2), which are believed to be identical in the biosynthesis of CAEs or COLs. The *trans* partners ColB1 and CaeB1 were confirmed to be interchangeable in the production of **1** (**Fig. 2a**, vii), supporting that they both share the *trans* flavin-dependent activity for oxidatively processing L-cysteinyl to dehydrocysteinyl on PCP. Consistent with this finding, ITC analysis showed that ColB1 can interact with the Ct domain of the CaeA2, with a moderate *K*_d_ value of 1.18 ± 0.15 μM (**Fig. 4**). Interestingly, changing the NRPS CaeA3 to ColA3 completely abolished **1** production (**Fig. 2a**, viii), indicating that ColA3 differs from CaeA3, even though they both mediate L-leucine extension.

*In vitro* testing the exchangeability between the PKS-NRPS hybrid proteins CaeA2 and ColA2 failed because ColA2 was highly resistant to various methods for soluble protein preparation. We thus turned to *in vivo* biochemical assays and examined this exchangeability in the COL-producing strain *Streptomyces roseosporus* (**Fig. 6**). A chimeric gene that encodes the variant harboring the PKS module of ColA2 and the NRPS module of CaeA2, *colA2_PKS-_caeA2_NRPS_*, was constructed and introduced into the previously developed, *ΔcolA2* mutant *S. roseosporus* strain that is incapable of producing COLs^14^. Remarkably, this complementation restored the production of COLs, supporting that the atypical NRPS modules of CaeA2 and ColA2 are functionally identical in terms of L-cysteinyl incorporation, *trans* CaeB1/ColB1 activity recruiting, condensation and heterocyclization during COL biosynthesis. These findings demonstrate the generality of both COLs and CAEs in 2,2’-bipyridine formation and particularly suggest that dethiolation can be skipped in the atypical NRPS module of CaeA2 to produce the thiolated 2,2’-bipyridine intermediate 5-sulfhydryl-2,2’-bipyridinyl. Notably, no dethiolated CAE-like products were observed in the *S. roseosporus* strain when *colA2* was replaced with *colA2_PKS-_caeA2_NRPS_* (**Fig. 6**). Most likely, the specificity of the L-leucine extension determines the fate of the sulfur as the substitution with the CaeA2 NRPS module does not change the product profile.

**Figure 6.**
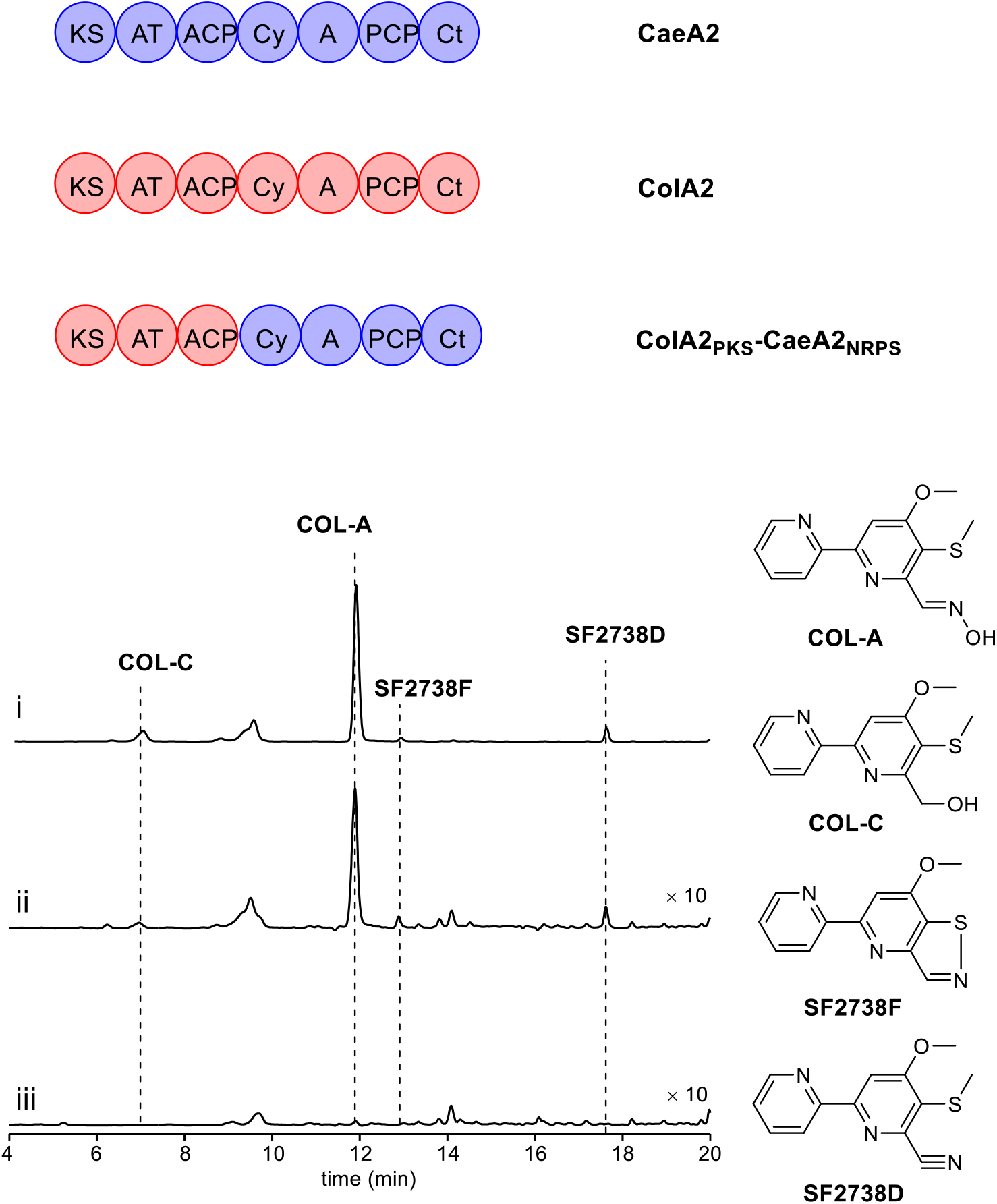
Domain organization of the NRPSs CaeA2, ColA2 and ColA2_PKS-_CaeA2_NRPS_ (Top) and HPLC analysis of COL-related products in *S. roseosporus* strains (below). i, the wild-type COL-producing strain (positive control); ii, the *ΔcolA2* mutant in which a chimeric gene *colA2_PKS-_caeA2_NRPS_* is expressed *in trans*; and iii, the *ΔcolA2* mutant (negative control).

### L-Leucine extension specializes 2,2’-bipyridine differentiation

Unlike 2,2’-bipyridinyl, sulfhydryl-2,2’-bipyridinyl formed in the assembly line is highly reactive and proved to be difficult for analysis by aforementioned HR-MS/MS approaches. To *in vitro* evidence this covalently tethered thiol intermediate, we attempted to examine the downstream off-loading intermediates by incubating CaeA1, CaeA2 and CaeB1 with ColA3 in the presence of the necessary substrates and ATP. Treating the reaction mixture with IAM revealed **17**, the derivative of the expected product 5-sulfhydryl-2,2’-bipyridinyl-L-leucine (**15**) (**Supplementary Fig. 16**). To further validate the production of **15**, we added ColG2, a COL pathway-specific methyltransferase^14,25^, into this reaction mixture. After L-leucine extension, ColG2 is believed to catalyze the *S*-methylation of the highly reactive thiol of intermediate **15** to produce the less reactive, previously characterized intermediate 5-methylmercapto-2,2’-bipyridinyl-L-leucine (**16**). Indeed, **16** was observed as the sole product (**Fig. 2a**, ix). During this *in vitro* conversion, the cycling of FADH_2_ to FAD can go through the formation of the oxidizing adduct FAD-4a-OOH, in agreement with a net oxidation occurring in sulfhydryl-2,2’-bipyridine formation (**Supplementary Fig. 7b**). In contrast, the combination of ColG2 with the complete CAE assembly line (i.e., CaeA1, CaeA2 and CaeA3 as well as CaeB1) failed to produce **16** and had little effect on the production of **1** (**Fig. 2a**, x), indicating that **15** cannot be generated in the presence of CaeA3. Furthermore, we narrowed the specificity of the upstream intermediate selection by CaeA3 and ColA3 to their C domains by domain swapping. Extensive attempts pointed to active CaeA3_C_ColA3_A-PCP_, the chimeric protein composed of the C domain of CaeA3 and the A-PCP didomain from ColA3. This protein was confirmed to be functionally identical to CaeA3 in the production of **1**, albeit with a lower yield (∼70% decrease) (**Fig. 2a**, xi). It cannot be used to replace ColA3 because **16** was not observed in the presence of ColG2. Clearly, the CAE and COL assembly lines are identical for 2,2’-bipyridine formation and can simultaneously provide the dethiolated and thiolated intermediates (**9** and **14**). The only exception is the C domain-conferred specificity in L-leucine extension, which specializes the pathway to the production of either CAEs or COLs.

## CONCLUSION

We demonstrate a common paradigm for 2,2’-bipyridine formation, which features a NRPS/PKS hybrid assembly line for sequential incorporation of the substrates picolinic acid, malonyl-*S*-CoA, L-cysteine and L-leucine with the association of a *trans* FAD-dependent partner (**Fig. 5**). This assembly line was first reconstituted *in vitro* for the biosynthesis of the dethiolated 2,2’-bipyridine antibiotics CAEs; then, the generality of 2,2’-bipyridine formation was validated in the biosynthesis of the distinct thiolated 2,2’-bipyridine antibiotics COLs, where the dethiolation step is skipped. In this assembly line, the atypical NRPS module Cy-A-PCP-Ct plays a central role in 2,2’-bipyridine formation. Following the A domain activity of this module for L-cysteine incorporation, the Ct domain recruits the activity of a flavoprotein that functions *in trans* for oxidatively processing L-cysteinyl on PCP. The Cy domain, which differs from the counterparts well-known for thiazoline formation, then condenses the resulting reactive dehydrocysteinyl extender unit with the upstream picolinyl-acetyl intermediate to afford a C-C bond via unusual Cβ-nucleophilic substitution, followed by intramolecular heterocyclization of the linear picolinyl-acetyl-dehydrocysteinyl intermediate to form the 2,2’-bipyridine core. Surprisingly, dethiolation can proceed or not between the condensation and heterocyclization steps, allowing for the production of both dethiolated and thiolated 2,2’-bipyridine intermediates in the atypical NRPS module. Subsequent L-leucine extension, which does not contribute any atoms in the final molecules, determines the sulfur fate and thus differentiates the biosynthesis of CAEs and COLs. Specifically, the C domain of the NRPS module C-A-PCP confers this selectivity for L-leucine extension and advances one of the two intermediates down a path to the 2,2’-bipyridine antibiotics with or without sulfur decoration. Therefore, whether dethiolation proceeds or is skipped appears to be not specifically controlled (i.e., not the key step) in the 2,2’-bipyridine assembly line because advancing the either intermediates (dethiolated or thiolated) strictly depends on the subsequent C domain selectivity for L-leucine extension. Based on this study, we provide a unique flavin-dependent paradigm in the assembly line, where cyclization reactions are usually difficult to be approached and thus remains poorly understood compared to those occurring at the off-line stage^3,4^. In addition, this study advances the understanding of the versatile functions of C domains in NRPS catalysis^36^. These domains include the gatekeeping C domain for sulfur fate determination, the Ct domain for *trans* flavin activity recruiting and, particularly, the unprecedented Cy domain for C-C bond formation and heterocyclization. The findings reported here further our appreciation of assembly line enzymology and will facilitate the design, development and utilization of enzymatic molecular machinery to address synthetic challenges arising from structurally-related complex polypeptides, polyketides and their hybrids.

**Supplementary Information** is available in the online version of the paper.

## Acknowledgements

We thank Dr. L. Yi (Beijing University of Chemical Technology) and Prof. Z. Xi (Nankai University) for providing the fluorescent probe in H_2_S examination. This work was supported in part by grants from NSFC (21750004, 21520102004, and 21621002), CAS (QYZDJ-SSW-SLH037 and XDB20020200), STCSM (17JC1405100), MST (2018ZX091711001-006-010) and K. C. Wang Education Foundation.

## SUPPLIMENTARY METHODS

### General materials and methods

Biochemicals and media were purchased from Sinopharm Chemical Reagent Co., Ltd. (China), Oxoid Ltd. (U.K.) or Sigma-Aldrich Co. LLC. (USA) unless otherwise stated. Restriction endonucleases were purchased from Thermo Fisher Scientific Co. Ltd. (USA). Chemical reagents were purchased from standard commercial sources. The bacterial strains, plasmids and primers used in this study are summarized in **Supplementary Tables 1, 2**, and **3**, respectively.

DNA isolation and manipulation in *E. coli* or actinobacteria were carried out according to standard methods^1,2^. PCR amplifications were carried out on an Applied Biosystems Veriti Thermal Cycler (Thermo Fisher Scientific Inc., USA) using either Taq DNA polymerase (Vazyme Biotech Co. Ltd, China) for routine verification or PrimeSTAR HS DNA polymerase (Takara Biotechnology Co., Ltd. Japan) for high fidelity amplification. Primer synthesis was performed at Shanghai Sangon Biotech Co. Ltd. (China). DNA sequencing was performed at Shanghai Majorbio Biotech Co. Ltd. (China).

High performance liquid chromatography (HPLC) analysis was carried out on an Agilent 1260 HPLC system (Agilent Technologies Inc., USA) equipped with a DAD detector. Semi-preparative HPLC was performed on an Agilent 1100 system equipped with a DAD detector (Agilent Technologies Inc., USA). HPLC Electrospray ionization MS (HPLC-ESI-MS) and tandem MS (MS/MS) for small molecules were performed on a Thermo Fisher LTQ Fleet ESI-MS spectrometer (Thermo Fisher Scientific Inc., USA), and the data were analyzed using Thermo Xcalibur software. NanoLC-MS/MS and MS/MS for peptides were performed on an EASY-nLC 1200 (Thermo Fisher Scientific Inc., USA) coupled with a Q Exactive HF mass spectrometer (Thermo Fisher Scientific Inc., USA), and the data were analyzed using pymzML, pFind^3^ and Thermo Xcalibur. High resolution ESI-MS (HR-ESI-MS) analysis for small molecules was carried out on an Agilent 6230B Accurate Mass TOF LC/MS System (Agilent Technologies Inc., USA), and the data were analyzed using Agilent MassHunter Qualitative Analysis software. NMR data were recorded on an Agilent 500 MHz PremiumCompact+ NMR spectrometer (Agilent Technologies Inc., USA).

### Protein expression and purification

The recombinant proteins CaeA1, PCP_CaeA2_, CaeB1, ColB1, ColG2, CaeA3, ColA3, and the chimeric NRPS protein CaeA3_C_ColA3_A-PCP_, were produced in N-terminal 6 × His-tagged forms, while CaeA2 and its variants (e.g., CaeA2^F2042L^, CaeA2^F2042I^, CaeA2^F2042V^, Ct _CaeA2_, CaeA2-ΔCt and CaeA2^F2042L^ΔCt) were expressed in C-terminal 8 × His-tagged forms or N-terminal 6 × His and Trx-tagged forms.

In general, the genes coding for the above proteins were amplified by PCR individually from the CAE-producing strain *Actinoalloteichus cyanogriseus* or the COL-producing strain *Streptomyces roseosporus*, both of which are listed in **Supplementary Table 1**, using the corresponding primers listed in **Supplementary Table 3**. The PCR products were first cloned into the vector pMD19-T for sequencing to confirm the fidelity and then into the expression vector pET-28a(+) (for N-terminal 6 × His-tagged proteins) or pET-37b(+) (for C-terminal 8 × His-tagged proteins). For expression of the chimeric NRPS protein CaeA3_C_ColA3_A-PCP_ (containing the C domain of CaeA3 and the A and PCP domains of ColA3), a 1.4 kb DNA fragment amplified by PCR from *A. cyanogriseus* using the primers CaeA3-C-For and CaeA3-C-Rev was cloned into pMD19-T, yielding pQL1046. Similarly, a 1.8 kb DNA fragment amplified by PCR from *S. roseosporus* using the primers ColA3-APCP-For and ColA3-APCP-Rev was cloned into pMD19-T, yielding pQL1047. The 1.4 kb NdeI-NheI fragment and the 1.8 kb NheI-HindIII fragment were recovered and co-ligated into the NdeI-HindIII site of pET-28a(+) to give pQL1048.

The resulting recombinant plasmids, which are listed in **Supplementary Table 2**, were transferred into *E. coli* BL21(DE3) for expression of CaeA2^F2042L^, PCP_CaeA2_, Trx-Ct _CaeA2_, CaeB1, ColG2 and ColB1 or into *E. coli* BAP1, an engineered strain capable of co-expressing the PPTase Sfp^4^, for expression of CaeA1, CaeA3, ColA3, and CaeA2 and its variants CaeA2^F2042L^, CaeA2^F2042I^, CaeA2^F2042V^, CaeA2-ΔCt, CaeA2^F2042L^ΔCt and CaeA3_C_ColA3_A-PCP_.

The culture of each *E. coli* transformant was incubated in Luria-Bertani (LB) medium containing 50 μg/mL kanamycin at 37°C and 220 rpm until the cell density reached 0.6-0.8 at OD_600_. Protein expression was induced by the addition of isopropyl-β-D-thiogalactopyranoside (IPTG) to the final concentration of 0.1 mM, followed by further incubation for 20 h at 25°C. The cells were harvested by centrifuging at 5000 rpm for 20 min at 4°C and were re-suspended in 30 mL of lysis buffer (50 mM Tris-HCl, 300 mM NaCl, 10% glycerol, pH 7.8). After disruption by FB-110X Low Temperature Ultra-pressure Continuous Flow Cell Disrupter (Shanghai Litu Mechanical Equipment Engineering Co., Ltd, China), the soluble fraction was collected and subjected to purification of each target protein using a HisTrap FF column (GE Healthcare, USA). The desired protein fractions, as determined by SDS-PAGE, were concentrated and desalted using a PD-10 Desalting Column (GE Healthcare, USA) according to the manufacturer’s protocols. The concentration of protein was determined by Bradford assay using bovine serum albumin (BSA) as the standard.

### Sequence analysis

Open reading frames (ORFs) were identified using the FramePlot 4.0beta program (http://nocardia.nih.go.jp/fp4/). The deduced proteins were compared with other known proteins in the databases using available BLAST methods (http://blast.ncbi.nlm.nih.gov/Blast.cgi). Amino acid sequence alignments were performed using Vector NT1 and ESPript 3.0 (http://espript.ibcp.fr/ESPript/ESPript/).

### Chemical synthesis of 4-hydroxy-2,2’-bipyridine-6-carboxyloyl-*S*-CoA (2,2’-bipyidinyl-*S*-CoA)

The precursor 4-hydroxy-2,2’-bipyridinyl-6-carboxylic acid hydrobromide (**17**) was synthesized according to the method described previously^4^.

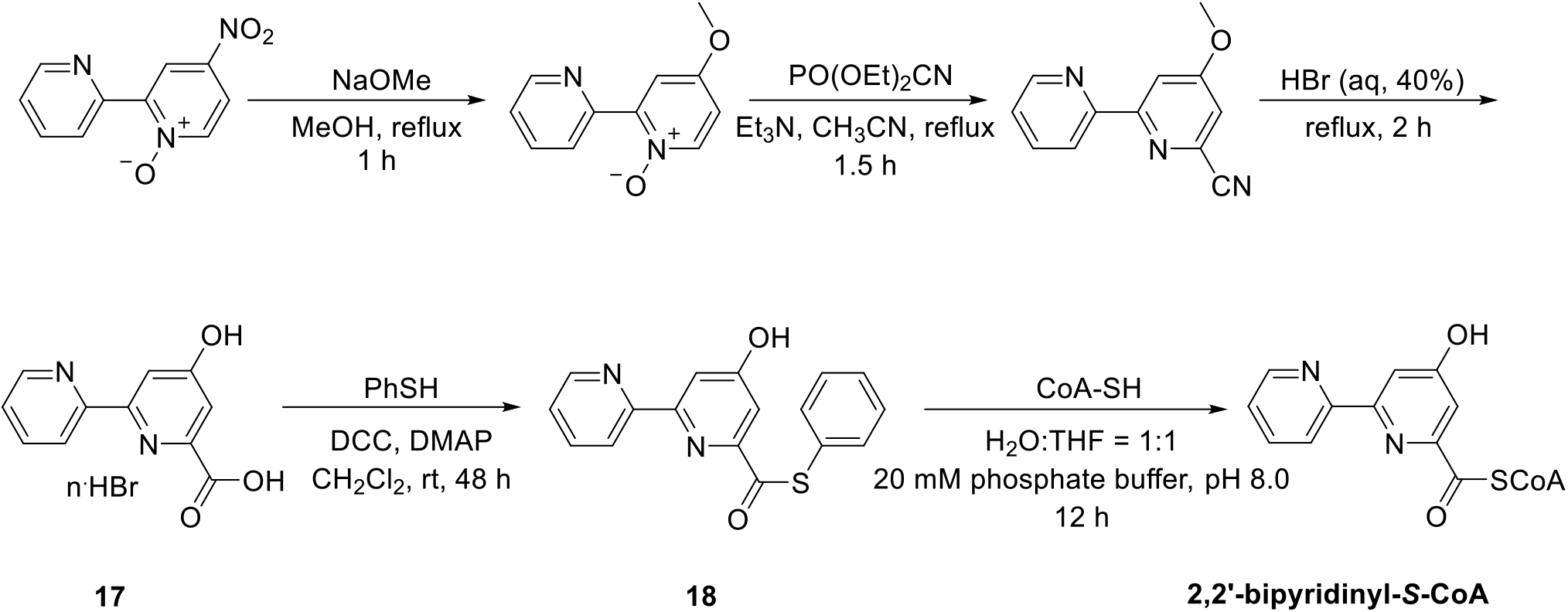

4-Hydroxy-2,2’-bipyridinyl-6-carboxylic acid hydrobromide (415 mg) was suspended in 15 mL of anhydrous CH_2_Cl_2_. Then, 721 mg of dicyclohexylcarbodiimide (DCC, 3.5 mmol), 0.58 mL of thiophenol (5.6 mmol) and 427 mg of 4-dimethylaminopyridine (DMAP, 3.5 mmol) were added into this CH_2_Cl_2_ suspension stirred at 0°C under argon atmosphere. The mixture was stirred at room temperature for 48 h before adding 0.3 mL of AcOH. After filtering, the organic phase of the reaction mixture was treated with saturated aqueous solution of NaHCO_3_ and brine, and subsequent concentrated to dryness under reduced pressure. The resulting residue was re-dissolved in EtOAc, washed with 5% (w/v) citric acid solution, and filtered over diatomite. The organic phase was then dried over anhydrous Na_2_SO_4_ overnight, filtered, and concentrated under reduced pressure. Further purification was carried out by chromatography on silica gel using 2:1 *n*-hexane/EtOAc as the eluent, followed by concentration to yield *S*-phenyl 4-hydroxy-2,2’-bipyridinyl-6-carbothioate (**18**, 50.2 mg). for the purified compound, ^1^H NMR (500 MHz, DMSO-*d*_6_,) *δ* 11.48 (s, 1H), 8.72 (ddd, *J* = 4.3, 1.5, 0.8 Hz, 1H), 8.49 (d, *J* = 7.9 Hz, 1H), 8.07 (d, *J* = 2.3 Hz, 1H), 8.04 (td, *J* = 7.7, 1.8 Hz, 1H), 7.57 – 7.48 (m, 6H), 7.30 (d, *J* = 2.3 Hz, 1H); and ^13^C NMR (125 MHz, DMSO-*d*_6_) *δ* 191.7, 166.9, 157.1, 154.2, 152.4, 149.9, 138.1, 135.3, 129.9, 129.7, 128.6, 125.3, 121.2, 112.1, 108.5. ESI-HRMS Calcd. for C_17_H_13_N_2_O_2_S^+^ 309.0698 [M+H]^+^, found 309.0696.

Next, 30 mg of CoA-SH (0.039 mmol, Sangon Biotech, China) was dissolved in 1.8 mL phosphate buffer (40 mM, pH = 8.0), and the pH was adjusted to 8.0 with 1 M NaOH, followed by the addition of the solution of 18 mg of the above obtained *S*-phenyl 4-hydroxy-2,2’-bipyridinyl-6-carbothioate in 1.8 mL THF, stirring at room temperature under argon atmosphere for 12 h. After removal of THF under reduced pressure, the mixture was washed with ether (to remove the unreacted thioester), and then subjected to semi-preparation by HPLC to give 2,2’-bipyridinyl-*S*-CoA as a white powder (8 mg). This semi-preparation was carried out on an Aglient Zorbax column (SB-C18, 5 μm, 9.4 x 250 mm, Agilent Technologies Inc., USA) by gradient elution of solvent A (H_2_O containing 5 mM NH_4_Ac) and solvent B (CH_3_CN) at a flow rate of 3 mL/min over a 30-min period as follows: T = 0 min, 5% B; T = 2 min, 5% B; T = 20 min, 90% B; T = 25 min, 90% B; T = 26 min, 5% B and T = 30 min, 5% B (mAU at 254 nm). For purified 2,2’-bipyridinyl-*S*-CoA, ESI-HRMS Calcd. for C_17_H_13_N_2_O_2_S^+^ 966.1682 [M+H]^+^, found 966.1670.

The CAE 2,2’-bipyridine intermediate **1** and the COL 2,2’-bipyridine intermediate **16** were synthesized using the methods previously reported^4^.

### Site-specific mutation of CaeA2 and its truncated proteins

Rolling cycle PCR amplification (using the primers listed in **Supplementary Table 3**) followed by subsequent DpnI digestion was conducted according to the standard procedure of the Mut Express II Fast Mutagenesis Kit (Vazyme Biotech Co. Ltd, China). The yielded recombinant plasmids were listed in **Supplementary Table 2**. Each mutation was confirmed by Sanger sequencing. CaeA2^F2042L^, CaeA2^F2042L^ΔCt, CaeA2^F2042I^, and CaeA2^F2042V^ were expressed in *E.coli* BAP1. All the mutant proteins were purified to homogeneity, and then concentrated according to the procedures for the native proteins described above.

### Determination of the flavin cofactor

Each protein (CaeB1 or ColB1) solution at the concentration of 1 mg/ml was incubated at 100°C for 5 min for denaturation and then subjected to HPLC-DAD analysis on an Agilent Zorbax column (SB-C18, 5 μm, 4.6 x 250 mm, Agilent Technologies Inc., USA) by gradient elution of solvent A (H_2_O containing 20 mM ammonium acetate) and solvent B (CH_3_CN) at a flow rate of 1 mL/min over a 35-min period as follows: T = 0 min, 5% B; T = 2 min, 5% B; T = 20 min, 90% B; T = 25 min, 90% B; T = 30 min, 5% B and T = 35, 5% B (δ at 448 nm), using standard FAD as control. The supernatant of CaeB1 or ColB1 was subjected to ESI-HRMS analysis to confirmed the identity of FAD (ESI *m/z* [M+H]^+^, calcd. 786.1644; found 786.1593).

### Reconstitution of the 2,2’-bipyridine assembly line *in vitro*

The initial reaction was conducted at 30°C for 1 h in a 100 μL reaction mixture containing 50 mM Tris-HCl (pH 7.5), 1 mM TCEP, 10 mM MgCl_2_, 1 mM picolinic acid, 1 mM malonyl-CoA, 100 μM L-cysteine, 1 mM L-leucine, 10 μM CaeA1, 1 μM CaeA2, 1 μM CaeA3, and 4 mM ATP. To determine the necessary *trans* partner, CaeB1, CaeA4 or both of them were added into the above mixture, respectively, with the final concentration of 1 μM for each protein. Each reaction was quenched with 100 μL of CH_3_CN after incubation. To examine the production of the dethiolated 2,2’-bipyridine intermediate **1**, reaction mixtures were subjected to HPLC analysis on an Agilent Zorbax column (SB-C18, 5 μm, 4.6 x 250 mm, Agilent Technologies Inc., USA) using a DAD detector, by gradient elution of solvent A (H_2_O containing 0.1% TFA) and solvent B (CH_3_CN containing 0.1% TFA) at a flow rate of 1 mL/min over a 35-min period as follows: T = 0 min, 5% B; T = 2 min, 5% B; T = 20 min, 90% B; T = 25 min, 90% B; T = 30 min, 5% B and T = 35, 5% B (δ at 315 nm). For HPLC-ESI-MS analysis, TFA was replaced by 0.1% formic acid.

To evaluate the participation of the enzymes and substrates in the production of **1**, each component was omitted from the 100 μL reaction mixture that contains 50 mM Tris-HCl (pH 7.5), 1 mM TCEP, 10 mM MgCl_2_, 1 mM picolinic acid, 1 mM malonyl-CoA, 100 μM L-cysteine, 1 mM L-leucine, 10 μM CaeA1, 1 μM CaeA2, 1 μM CaeA3, 1 μM CaeB1, and 4 mM ATP. To examine O_2_ dependence, gas exchange for O_2_ elimination was conducted in an anaerobic glovebox overnight before incubation under anaerobic conditions. To evaluate the mutation effects of CaeA2 on **1** production, CaeA2 was replaced with its variants CaeA2^F2042L^, CaeA2^F2042I^, CaeA2^F2042V^, CaeA2-ΔCt or CaeA2^F2042L^ΔCt. To mechanistically trace the transformation process, L-[1,2,3-^13^C_3_,^15^N]cysteine and L-[2,3,3-D_3_]cysteine were used to replace unlabeled L-cysteine, respectively. To evaluate the changeability of the enzymes in the reaction mixture, 1) CaeB1 was replaced with ColB1 (with the final concentration of 10 μM); and 2) CaeA3 was replaced with ColA3 and CaeA3_C_ColA3_A-PCP_, respectively. Reactions are conducted at 30°C for 1 h and then quenched with 100 μL of CH_3_CN. The production of **1** was monitored by HPLC or HPLC-ESI-MS under conditions as described above.

For H_2_S examination during **1** production, the reaction was conducted at 30°C for 1 h in the 100 μL, TCEP-involving reaction mixture that contains 50 mM Tris-HCl (pH 7.5), 1 mM TCEP, 10 mM MgCl_2_, 1 mM picolinic acid, 1 mM malonyl-CoA, 100 μM L-cysteine, 1 mM L-leucine, 10 μM CaeA1, 1 μM CaeA2, 1 μM CaeA3, 1 μM CaeB1, and 4 mM ATP. Then, the TCEP derivative **2** was analyzed by HPLC-ESI-MS under conditions as described above. Alternatively, H_2_S examination was conducted in the reaction mixture where TCEP was omitted, in the presence of **19** (1 μM), a dual-reactable fluorescent probe used for highly selective and sensitive detection of biological H_2_S ^5^. The reaction of **19** with Na_2_S to yield **20** serves as the control reaction. **20** was examined by HPLC-FLD under conditions as described above with excitation at 370 nm and relative emission at 450 nm.

To examine whether the CAE assembly line provides the thiolated intermediate in COL biosynthesis, ColG2 (with the final concentration of 1 μM) was added into the 100 μL reaction mixture that contains 50 mM Tris-HCl (pH 7.5), 1 mM TCEP, 10 mM MgCl_2_, 1 mM picolinic acid, 1 mM malonyl-CoA, 100 μM L-cysteine, 1 mM L-leucine, 10 μM CaeA1, 1 μM CaeA2, 1 μM CaeB1, 1 μM CaeA3 (ColA3 or CaeA3_C_ColA3_A-PCP_), and 4 mM ATP. The production of the thiolated 2,2’-bipyridine intermediate **16** was examined by HPLC or HPLC-ESI-MS under conditions as described above.

For sulfhydryl-2,2’-bipyridinyl-L-leucine (**15**) examination, the reaction was conducted at 30°C for 1 h in the 50 μL reaction mixture contains 50 mM Tris-HCl (pH 7.5), 1 mM TCEP, 10 mM MgCl_2_, 1 mM picolinic acid, 1 mM malonyl-CoA, 100 μM L-cysteine, 1 mM L-leucine, 10 μM CaeA1, 1 μM CaeA2, 1 μM ColA3, 1 μM CaeB1, and 4 mM ATP. The reaction mixture was treated with 1% SDS and 5 mM TCEP at 55°C for 30 min to release the free thiol of **15**, and then was incubated with 25 mM iodoacetamide (IAM) at 30°C for 30 min. The production of **17** was examined by HPLC or HPLC-ESI-MS under conditions as described above.

### *In vitro* assays of PCP *S*-aminoacylation on CaeA2 by nanoLC-MS/MS

The CaeA2 recombinant protein that was purified from *E.coli* BAP1 was subjected to complete or partial protease hydrolysis with trypsin, Glu-C or chymotrypsin as well as a variety of their combinations to map **SLGGD<u>S</u>IMGIQF_2042_VSR** of CaeA2, the MS-detectable sequence that contains the Ppant-modified active-site L-serine residue (underlined). The digestion mixtures were filtrated using Microcon YM-10 (MilliporeSigma, USA) by centrifugation and stored at − 80°C before analysis. For nanoLC-MS/MS analysis, each sample was loaded on a trap column (75 μm i.d., 2 cm, C18, 5 μm, 100 Å, Thermo Fisher Scientific Inc., USA) for online desalting, and then was separated using a reversed phase column (75 μm i.d., 10.2 cm, C18, 3 μm, 120 Å, New Objective Inc., USA) by gradient elution of solvent A (H_2_O containing 0.1% formic acid) and solvent B (80% CH_3_CN containing 0.1% formic acid) at a flow rate of 300 nL/min over a 1.5 h period as follows: T = 0 min, 18% B; T = 45 min, 45% B; T = 50 min, 100% B; and T = 90 min, 100% B. For MS analysis, the nano-ESI voltage and capillary temperature were set at 2.2 kV and 275°C, respectively. The MS data were acquired in data-dependent mode. Each full-scan MS (*m/z* 350−2000, resolution of 60 k) was followed with 10 HCD MS/MS scans (normalized collision energy of 33, resolution of 15 k) for the most intense precursor ions. The maximum ion injection time for MS and MS/MS were 50 and 45 ms, and the auto gain control target for MS and MS/MS were 3 × 10^6^ and 5 × 10^4^, respectively. The dynamic-exclusion time was set as 40 s.

To obtain the sequence **SLGGD<u>S</u>IMGIQF_2042_VSR**, the 50 μL solution that contains 50 mM Tris-HCl (pH 7.5) and 5 μM CaeA2 was treated with 4 μg of trypsin (sequencing grade, Promega Corp., USA) at 30°C for 20 min (leading to complete digestion) and then with 0.6 μg of chymotrypsin (sequencing grade, Promega Corp., USA) at 30°C for 10 min (leading to partial digestion that retains the C-terminal sequence **F_2042_VSR**). To obtain the engineered sequence **SLGGD<u>S</u>IMGIQL_2042_VSR**, the CaeA2^F2042L^ recombinant protein that was purified from *E.coli* BAP1 and its variants different in *S*-(amino)acylation underwent complete digestion with trypsin (4 μg) and chymotrypsin (0.6 μg) at 30°C for 20 min.

To identify the target sequence that contains the Ppant-modified active-site L-serine residue, the raw MS data was processed using the pFind software by setting Ppant as a variable posttranslational modification. The selected sequence was then validated by HR-MS/MS analysis of the parent ion, its associated fragmented peptide ions and particularly the characteristic Ppant ejection ion^6^.

To prepare L-cysteinyl-*S*-CaeA2^F2042L^ (**3**), the reaction was conducted at 30°C for 10 min in a 50 μL reaction mixture containing 50 mM Tris-HCl (pH 7.5), 1 mM TCEP, 10 mM MgCl_2_, 100 μM L-cysteine, 5 μM CaeA2^F2042L^ (or CaeA2^F2042L^ΔCt) and 4 mM ATP. To prepare (3-sulfhydryl)-pyruvoyl-*S*-CaeA2^F2042L^ (**6**), the reaction was conducted at 30°C for 10 min in a 50 μL reaction mixture containing 50 mM Tris-HCl (pH 7.5), 1 mM TCEP, 10 mM MgCl_2_, 100 μM L-cysteine, 5 μM CaeA2^F2042L^ (or CaeA2^F2042L^ΔCt), 1 μM CaeB1 and 4 mM ATP. To prepare 2,2’-bipyridinyl-*S*-CaeA2^F2042L^ (**9**), the reaction was conducted at 30°C for 10 min in a 50 μL reaction mixture containing 50 mM Tris-HCl (pH 7.5), 1 mM TCEP, 10 mM MgCl_2_, 1 mM picolinic acid, 1 mM malonyl-CoA, 100 μM L-cysteine, 1 μM CaeA1, 1 μM CaeB1, 5 μM CaeA2^F2042L^ (or CaeA2^F2042L^ΔCt) and 4 mM ATP. For isotope labeling, L-[1,2,3-^13^C_3_,^15^N]cysteine and L-[2,3,3-D_3_]cysteine were used to replace unlabeled L-cysteine, respectively. For thiol derivatization, each reaction mixture was treated with 10 mM iodoacetamide (IAM) at 30°C for 2 min. Protease digestion and subsequent nanoLC-MS/MS analyses were conducted using approaches as described above.

To probe potential intermediates, all MS/MS data of the derivatives from the sequence (**SLGGD<u>S</u>IMGIQL_2042_VSR)** were extracted from raw MS data using script written by Python (https://github.com/billpb610/AlFinder/blob/master/AlFinderS2.py). The script also recorded information about precursor ions, fragment iond and Ppant ejection ions for further analysis.

### *In vitro* preparation of 2,2’-bipyridinyl-*S*-CaeA2^F2042L^ (9)

To prepare **9** as a positive control in *S*-aminoacylation assays, the reaction was conducted at 30°C for 30 min in a 50 uL reaction mixture containing 50 mM Tris-HCl (pH 7.5), 1 mM TCEP, 10 mM MgCl_2_, 100 μM 2,2’-bipyridinyl-S-CoA, 5 μM Sfp, and 10 μM CaeA2^F2042L^ in apo-form. Protease digestion and subsequent nanoLC-MS/MS analyses were conducted using approaches as described above.

### *In vitro* preparation of 2,2’-bipyridinyl-*S*-PCP_CaeA2_ (10)

The reaction was conducted at 30°C for 1 h in an 80 μL reaction mixture containing 62.5 mM Tris^-^HCl (pH = 7.5), 1.25 mM TCEP, 12.5 mM MgCl_2_, 1 mM 2,2’-bipyridinyl-*S*-CoA, 100 μM PCP_CaeA2_ (derived from wild-type CaeA2) in apo-form, and 4 μM Sfp.

To produce the dethiolated 2,2’-bipyridine intermediate **1**, the reaction mixture was combined with the following components in a new 100 μL reaction mixture: 1 mM L-leucine, 5 μM CaeA3, and 4 mM ATP. After incubation at 30°C for 2 h, the reaction was quenched with 100 μL of CH_3_CN for HPLC or HPLC-ESI-MS analysis under conditions as described above.

### Measurement of protein-protein interactions by isothermal titration calorimetry (ITC)

ITC was performed with MicroCal-ITC200 (Malvern) at 25°C. A 400 μl aliquot of 60 μM Ct or CaeA2ΔCt was placed in the stirred cell, and 120 μl aliquot of 400 μM MBP-fused CaeB1 or ColB1 was prepared in the syringe. All the recombinant proteins were prepared by Ni-affinity chromatography followed by size-exclusion chromatography (SEC) for purification, and then were exchanged in the 50 mM Tris-HCl (PH = 7.5) buffer containing 100 mM NaCl, 0.1 mM TCEP and 5 % glycerol. The titration was performed as follows: 1 μl of protein in the syringe over 0.8 s for the first injection, followed by 19 injections of 2 μl protein in the stirred cell at 120 s intervals. The heat of reaction per injection was determined by integration of the peak areas using the MicroCal-PEAQ-ITC software, which provides the best-fit values for the heat of binding (ΔH), the stoichiometry of binding (N), and the dissociation constant (*K*_d_). The heats of dilution were determined by injecting flavoprotein alone into the buffer and were subtracted from the corresponding experiments before curve fitting.

### Analysis of the functional exchangeability between CaeA2 and ColA2 *in vivo*

A 4.4 kb DNA fragment amplified by PCR from the CAE-producing *A. cyanogriseus* strain using the primers cae-CO-for and cae-Ct-rev was cloned into pMD19-T, yielding pQL1053. After sequencing to confirm the fidelity, this PCR product was recovered by digestion with BglII-EcoRV and then utilized to replace the 4.4 kb BglII-EcoRV fragment of pQL1022^7^, a pSET152 derivative previously constructed for *colA2* expression under the control of *PermE\** (the constitutive promoter for expressing the erythromycin-resistance gene in *Saccharopolyspora erythraea*). The resulting recombinant plasmid pQL1054, which carries the chimeric gene *col/caeA2* coding for the hybrid protein that harbors the PKS module from ColA2 and the NRPS module from CaeA2, was transferred by conjugation into the *ΔcolA2 S. roseosporus* mutant strain^7^, yielding the recombinant strain QL2006. The fermentation of QL2006 and the examination of CAE or COL production were conducted according the methods described previously^7^.

## SUPPLIMENTARY FIGURES

**Supplementary Figure 1.**
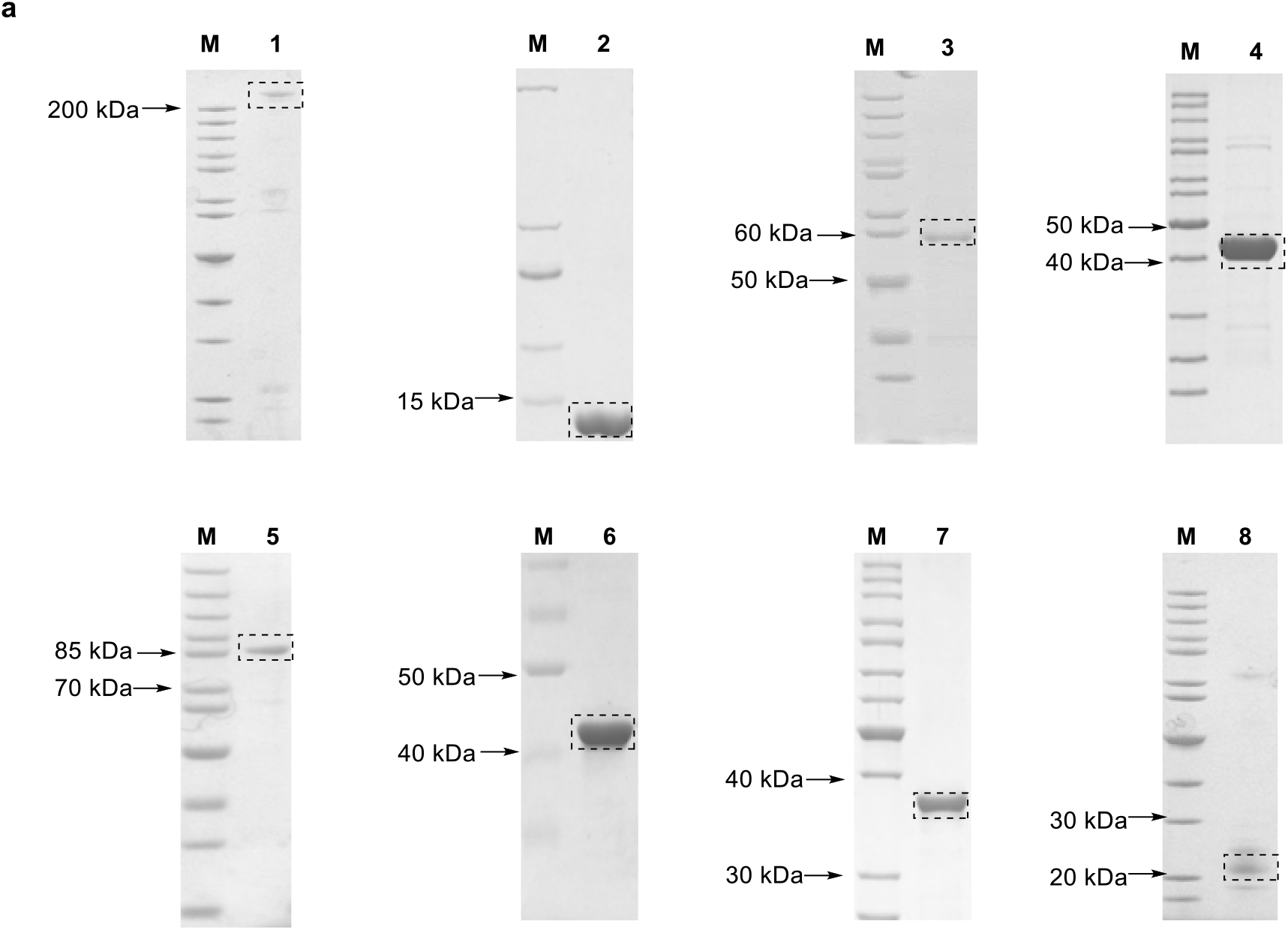
Coomassie-stained SDS-PAGE analysis of purified recombinant proteins. Lane M, protein standard. The target proteins are indicated in the dashed rectangles. (a) Proteins produced in *E. coli* BL21(DE3). Lane 1, CaeA2^F2042L^ (269 kDa); Lane 2, PCP_CaeA2_ (derived from CaeA2, 13 kDa); Lane 3, Trx-tagged Ct_CaeA2_ (58 kDa), Lane 4, CaeB1 (43 kDa); Lane 5, MBP-fused CaeB1 (87 kDa), Lane 6, ColB1 (43 kDa); Lane 7, ColG2 (39 kDa); and Lane 8, CaeA4 (25 kDa).

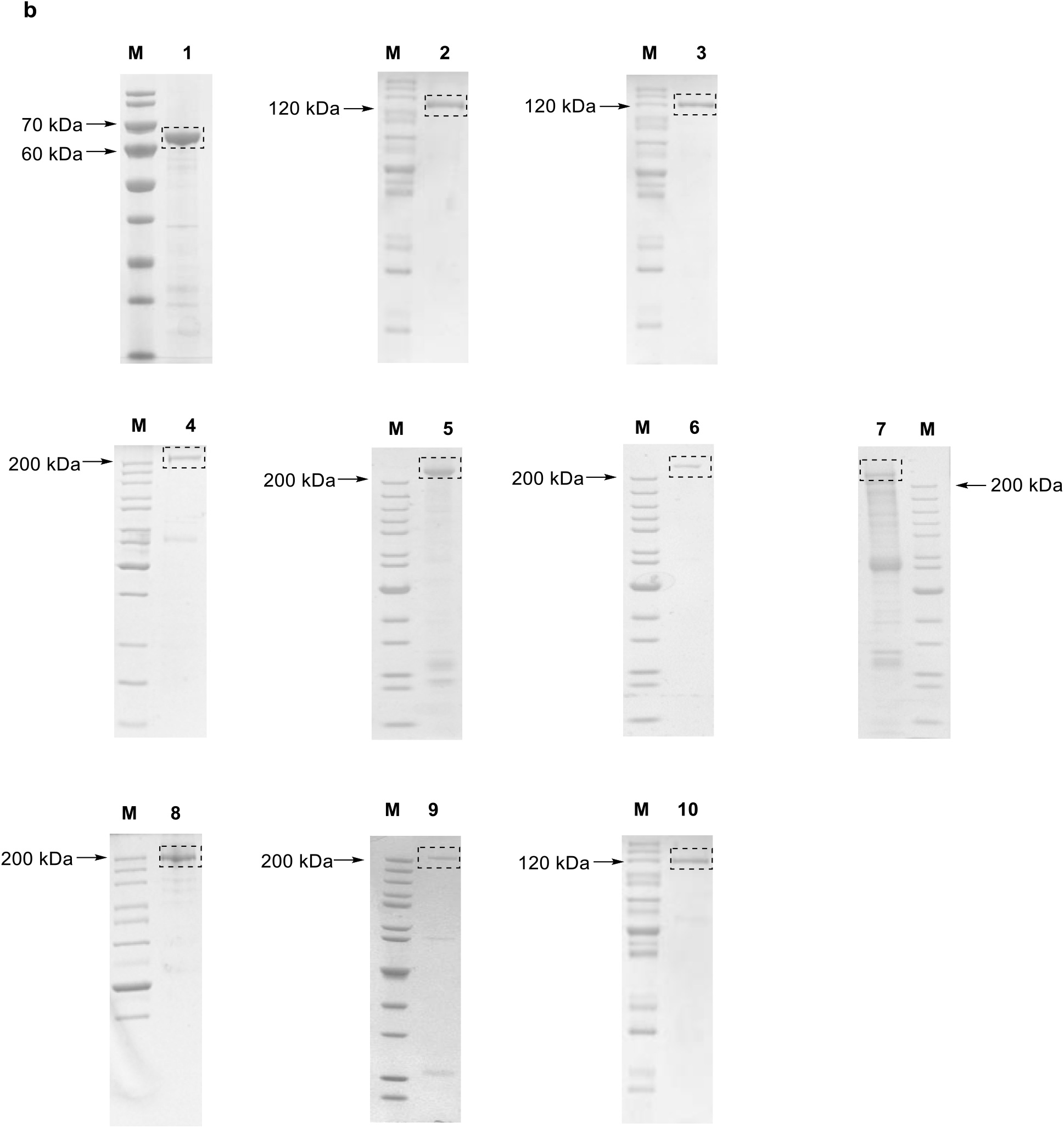
(b) Proteins produced in *E. coli* BAP1. Lane 1, CaeA1 (69 kDa); Lane 2, CaeA3 (118 kDa); Lane 3, ColA3 (121 kDa); Lane 4, CaeA2 (269 kDa); Lane 5, CaeA2^F2042L^ (269 kDa); Lane 6, CaeA2^F2042I^ (269 kDa); Lane 7, CaeA2^F2042V^ (269 kDa); Lane 8, CaeA2-ΔCt (225 kDa); Lane 9, CaeA2^F2042L^ΔCt (225 kDa); and Lane 10, CaeA3_C_ColA3_A-PCP_ (120 kDa).

**Supplementary Figure 2.**
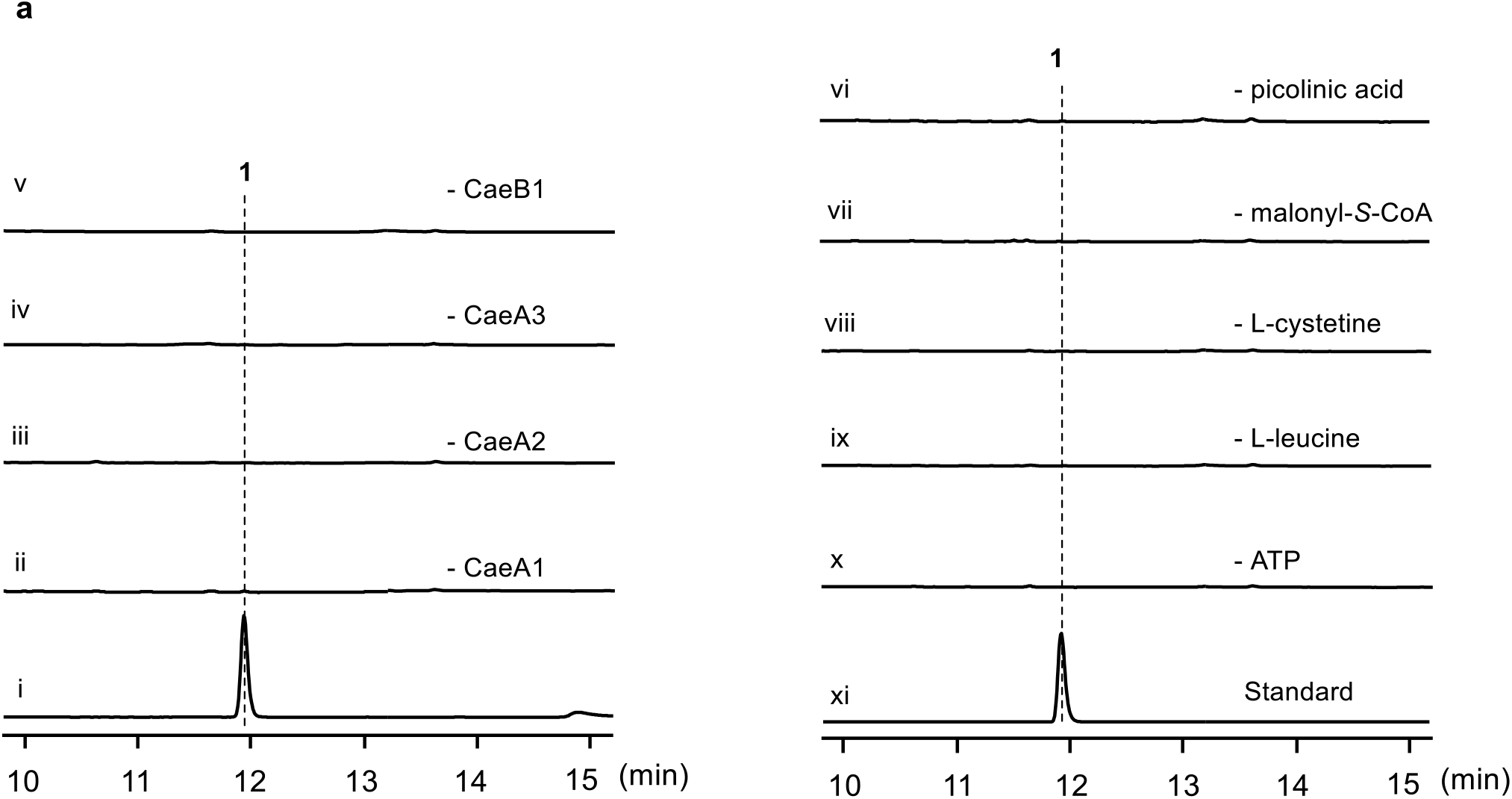
Supplementary data for *in vitro* reconstitution of the 2,2’-bipyridine assembly line. (a) Determination of component necessity in the production of the CAE 2,2’-bipyridine intermediate **1**. Tested reactions were derived from the combination of CaeA1, CaeA2, CaeA3 and CaeB1 with picolinic acid, malonyl-*S*-CoA, L-cysteine, L-leucine, and ATP (i), and included those lacking the enzyme CaeA1 (ii), CaeA2 (iii), CaeA3 (iv) or CaeB1 (v) and the substrate picolinic acid (vi), malonyl-*S*-CoA (vii), L-cysteine (viii), L-leucine (ix) or ATP (x, for the activity of the A domains), respectively. Synthesized **1** was used as the stabdard (xi).

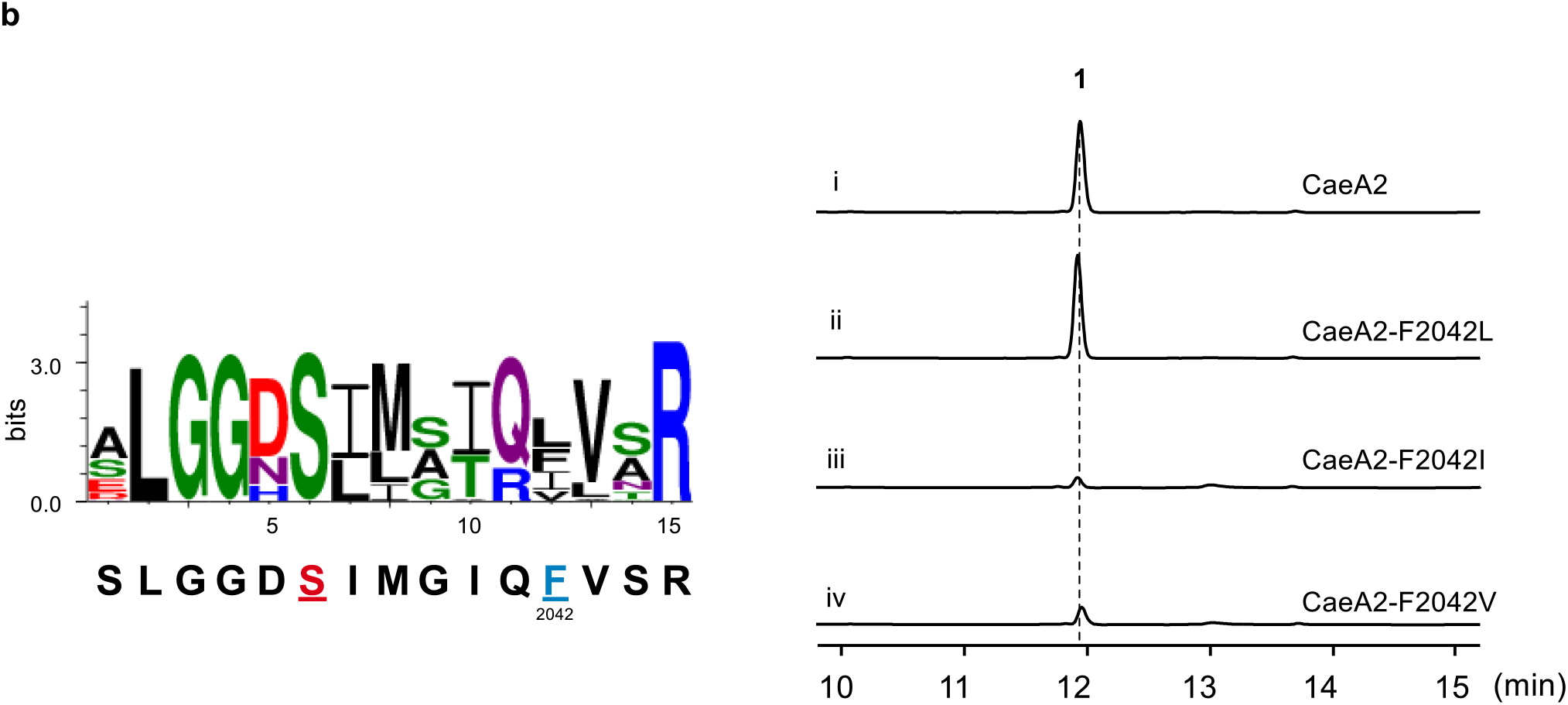
(b) Activity assays of CaeA2 variants in the production of **1**. The CaeA2 variants, i.e., CaeA2^F2042L^, CaeA2^F2042V^ and CaeA2^F2042I^, were designed according to residue conservative analysis (left), which was conducted by comparing the 15-aa sequence **SLGGD<u>S</u>IMGIQ<u>F_2042_</u>VSR** that is derived from the PCP domain of CaeA2 (the Ppant-modified active-site L-serine residue and the target L-phenylalanine residue are highlighted in color and underlined) with the corresponding sequences of 500 homologous PCP domains or proteins from NCBI NR database. Tested reactions were derived from the combination of CaeA1, CaeA2, CaeA3 and CaeB1 with picolinic acid, malonyl-*S*-CoA, L-cysteine, L-leucine, and ATP (i), and included those in which CaeA2 was replaced with CaeA2^F2042L^ (ii), CaeA2^F2042V^ (iii) and CaeA2^F2042I^ (iv), respectively.

**Supplementary Figure 3.**
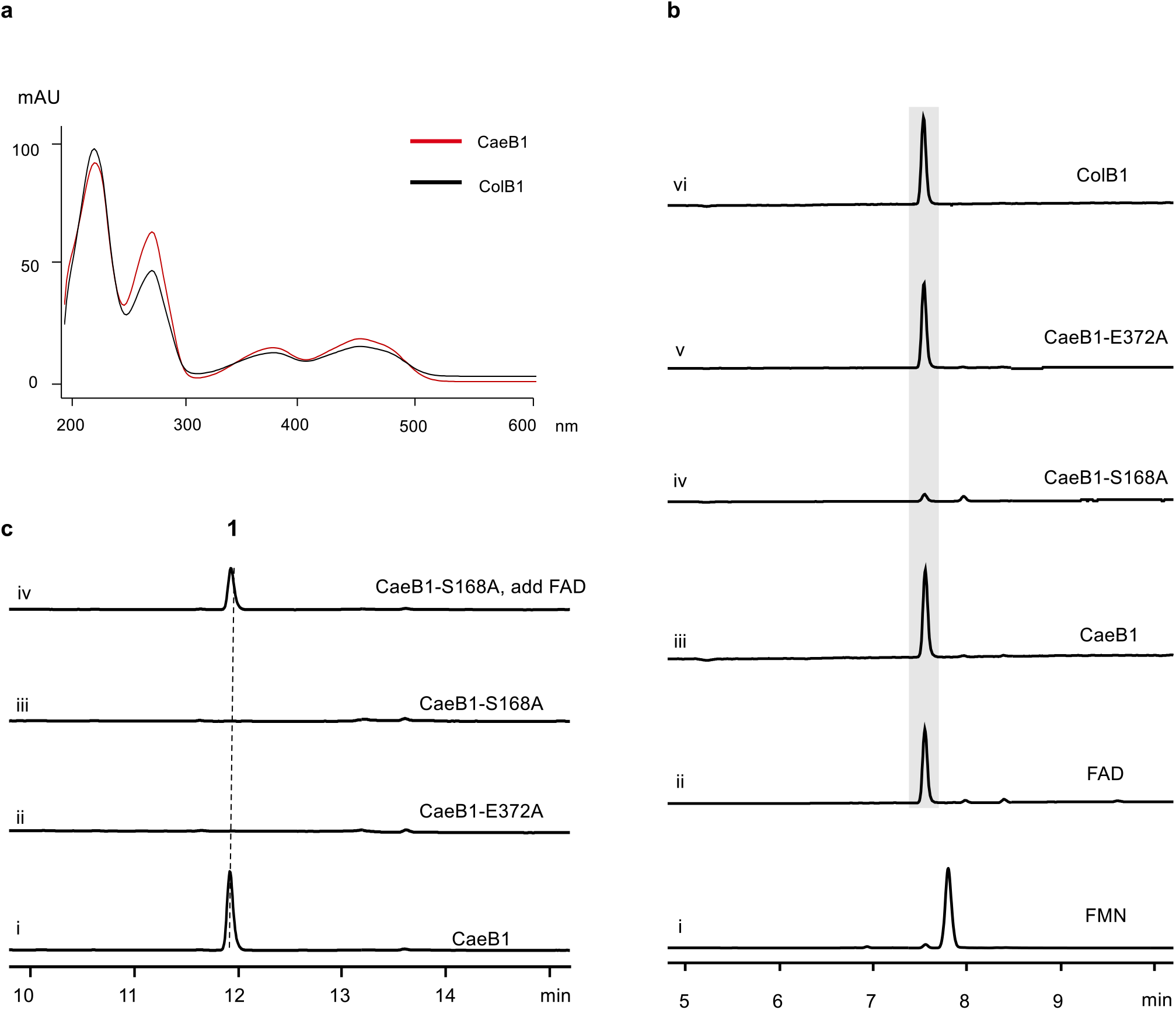
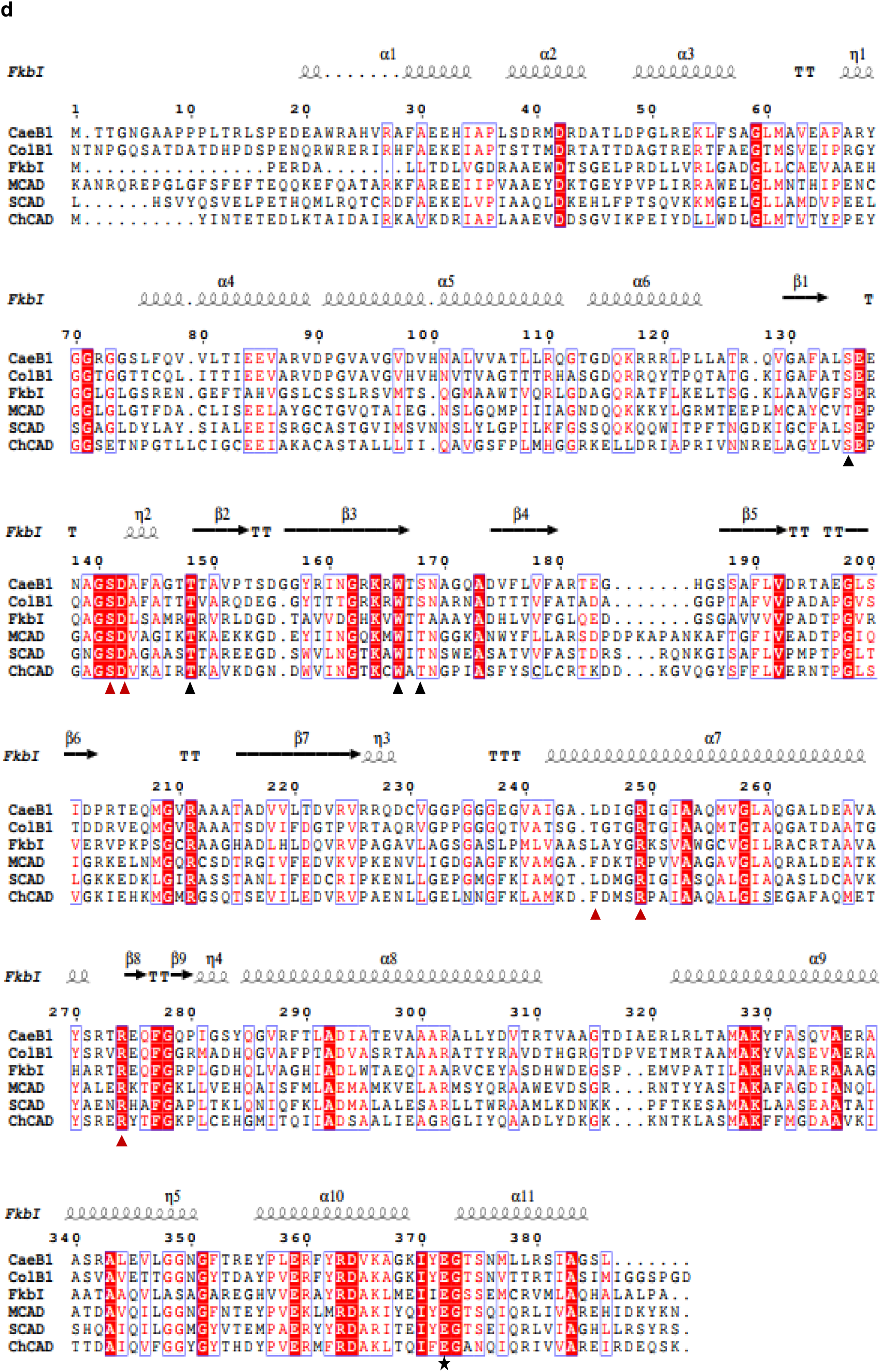
Analysis of the flavoproteins CaeB1 and ColB1. (a) UV spectra of the recombinant proteins CaeB1 and ColB1 that were purified from *E. coli* BL21(DE3). (b) Determination of flavin cofactors associated with CaeB1 and ColB1. (i) authentic FMN; (ii) authentic FAD, (iii) boiled CaeB1, (iv) boiled CaeB1-S168A, (v) boiled CaeB1-E372A, and (vi) boiled ColB1. For examination by HPLC, UV absorbance at 375 nm. (c) Activity assays of CaeB1 variants in the production of **1**. Tested reactions were derived from the combination of CaeA1, CaeA2, CaeA3 and CaeB1 with picolinic acid, malonyl-*S*-CoA, L-cysteine, L-leucine, and ATP (i), and included those in which CaeB1 was replaced with CaeB1-E372A (ii), CaeB1-S168A (iii), and CaeB1-S168A with excess of FAD (iv), respectively. (d) Sequence alignment of CaeB1 and ColB1 with various FAD-dependent dehydrogenases. The homologs include FkbI (1R2G) in FK520 biosynthesis^8^, medium chain acyl-CoA dehydrogenase (MCAD, 1T9G_A)^9^, short chain acyl-CoA dehydrogenase (SCAD, 1JQI_A)^10^ and cyclohexanecarboxyl-CoA dehydrogenase (ChCAD, ABC76100.1)^11^. Residues important for dehydrogenase activity, FAD-binding and phosphopentatheine binding are indicated by star, black triangle and red triangle, respectively.

**Supplementary Figure 4.**
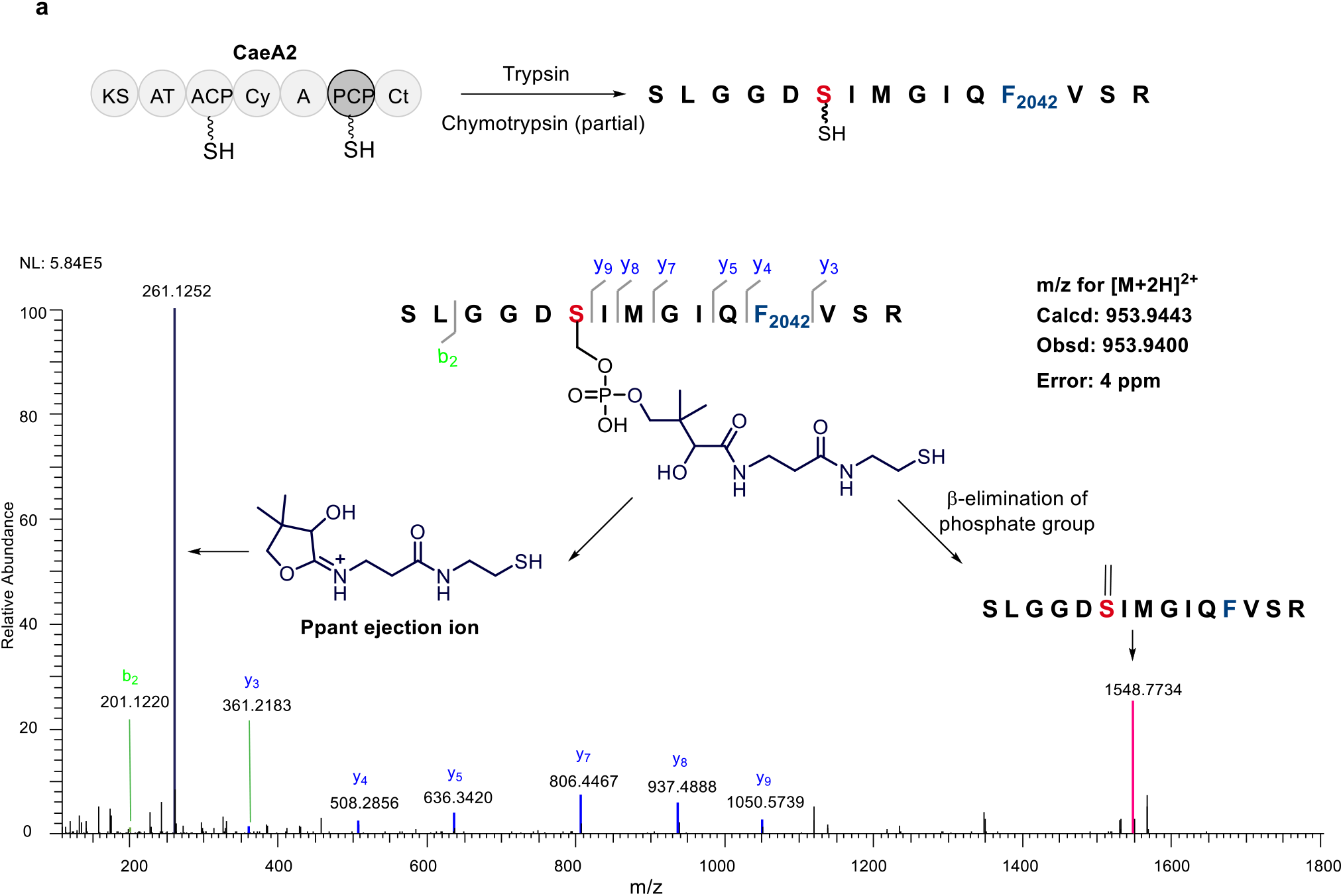
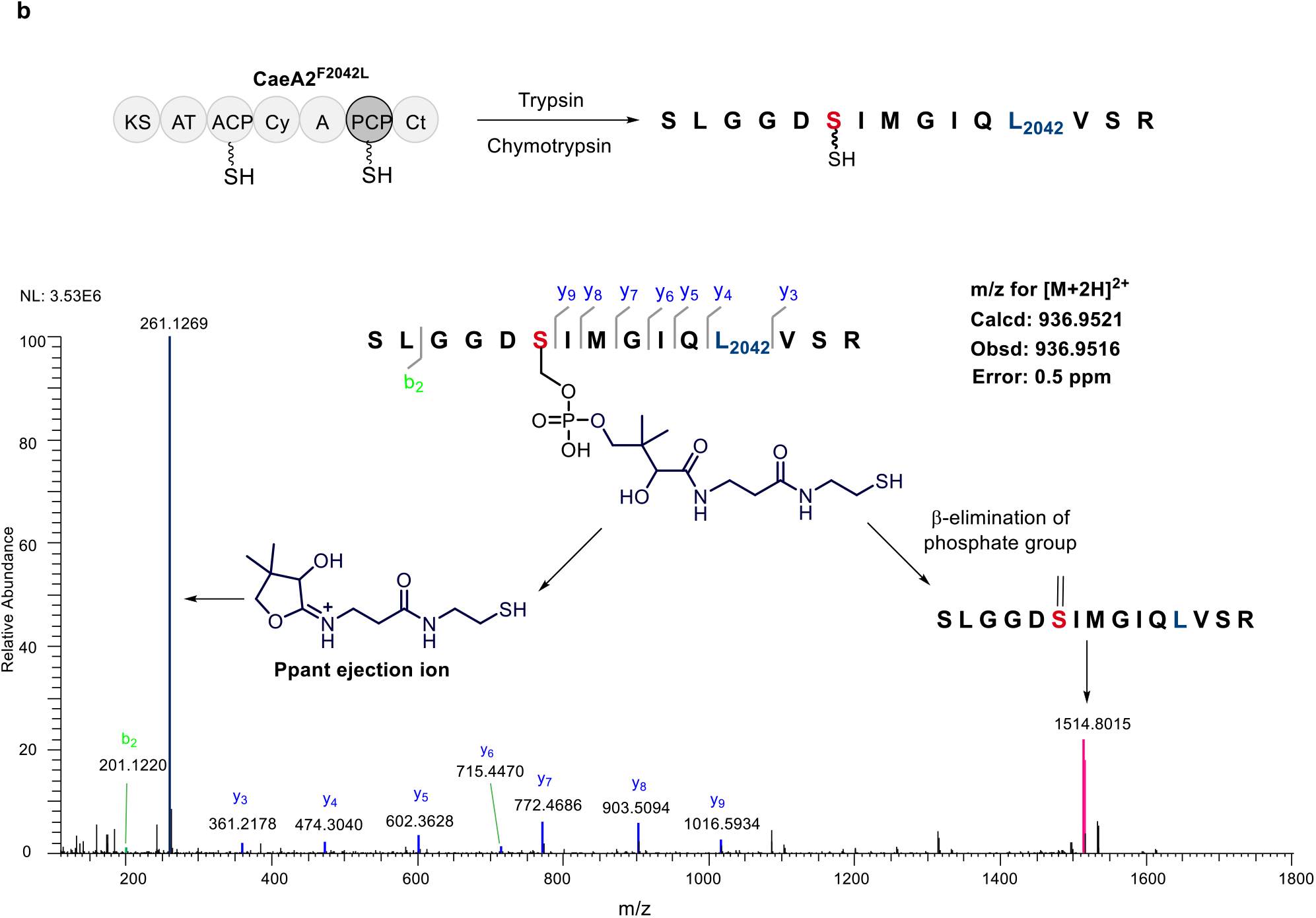
Determination of the 15-aa sequence that is derived from the PCP domain of CaeA2 and contains the Ppant-modified active-site L-serine residue (red) by nanoLC-MS/MS. (**a**) Ppant-modified CaeA2 by treatment with trypsin (complete digestion) and chymotrypsin (partial digestion). (**b**) Ppant-modified CaeA2^F2042L^ by treatment with trypsin (complete digestion) and chymotrypsin (complete digestion).

**Supplementary Figure 5.**
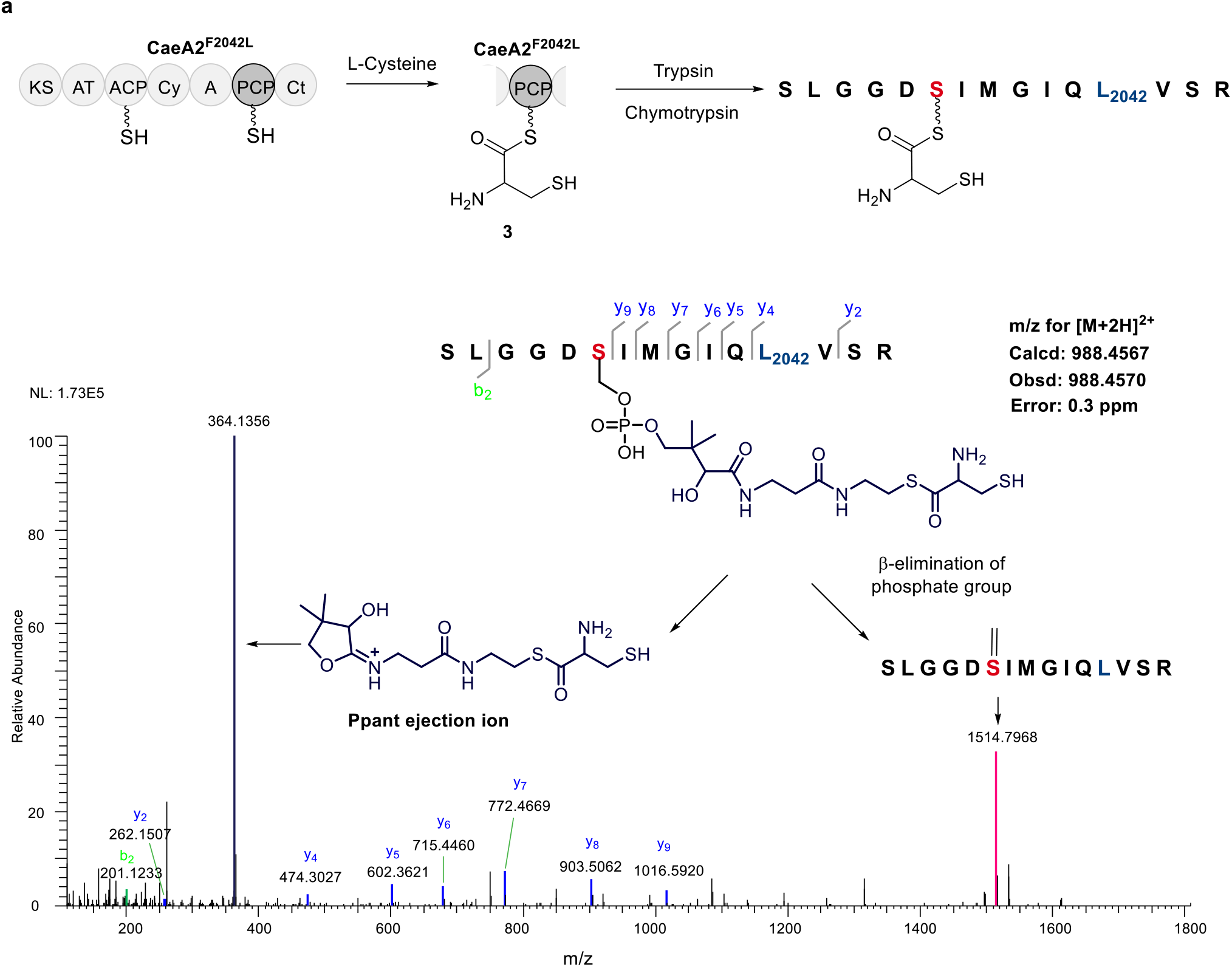
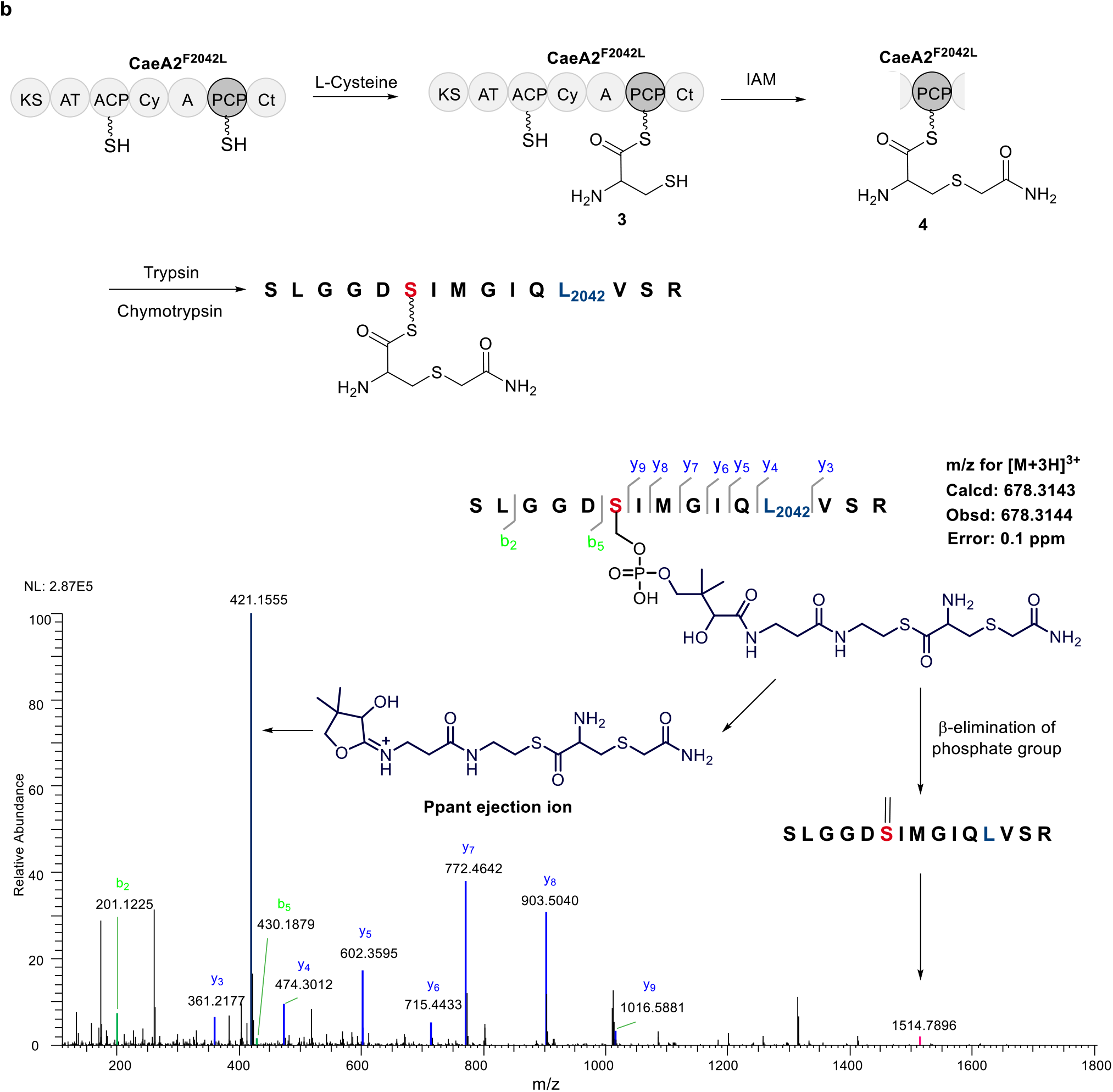
Characterization of L-cysteinyl-*S*-CaeA2^F2042L^ (**3**) by nanoLC-MS/MS following complete digestion with trypsin and chymotrypsin. (**a**) Incubation of thiolated CaeA2^F2042L^ and L-cysteine. (**b**) Treatment of with IAM after the incubation of thiolated CaeA2^F2042L^ and L-cysteine.

**Supplementary Figure 6.**
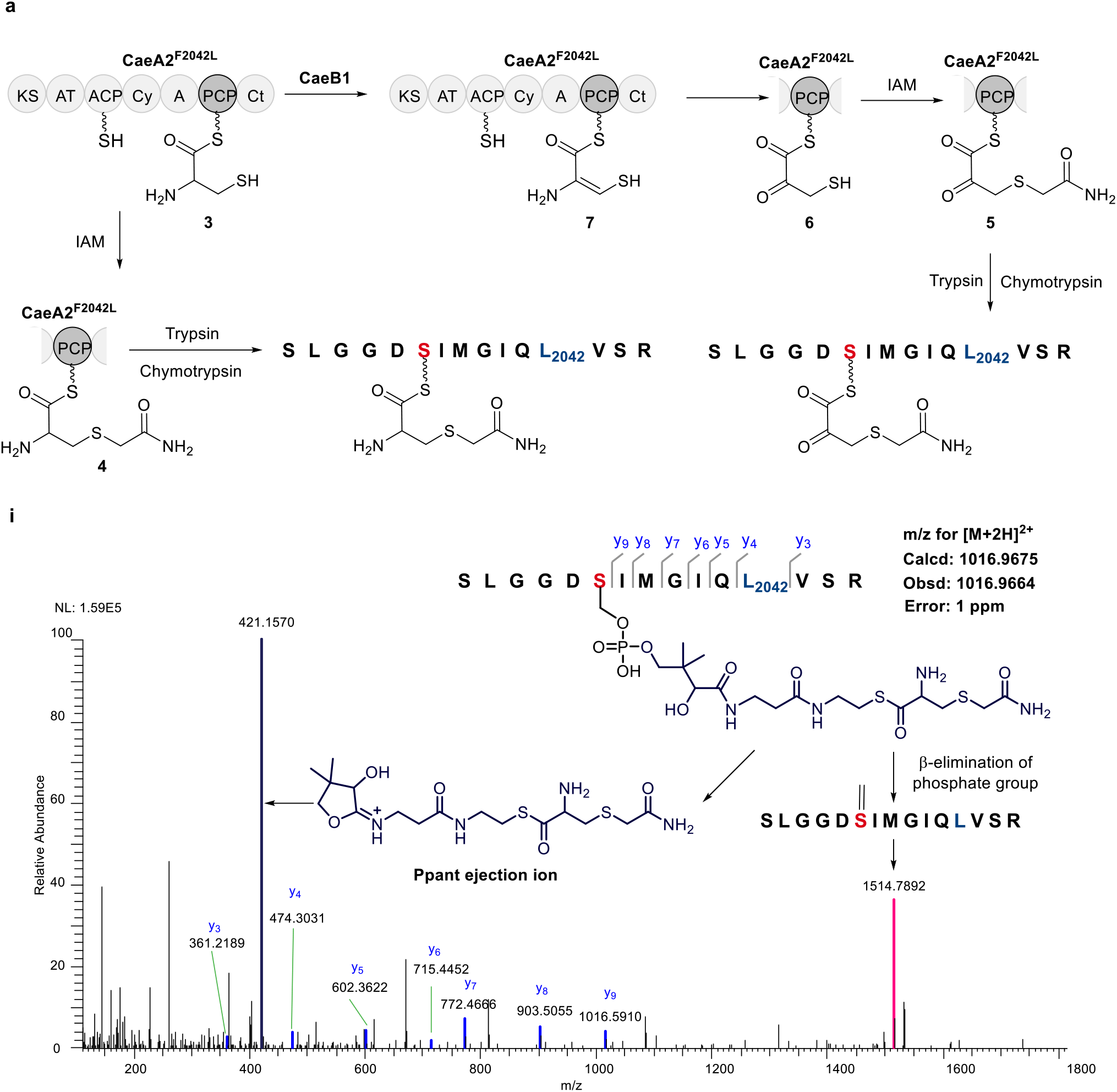
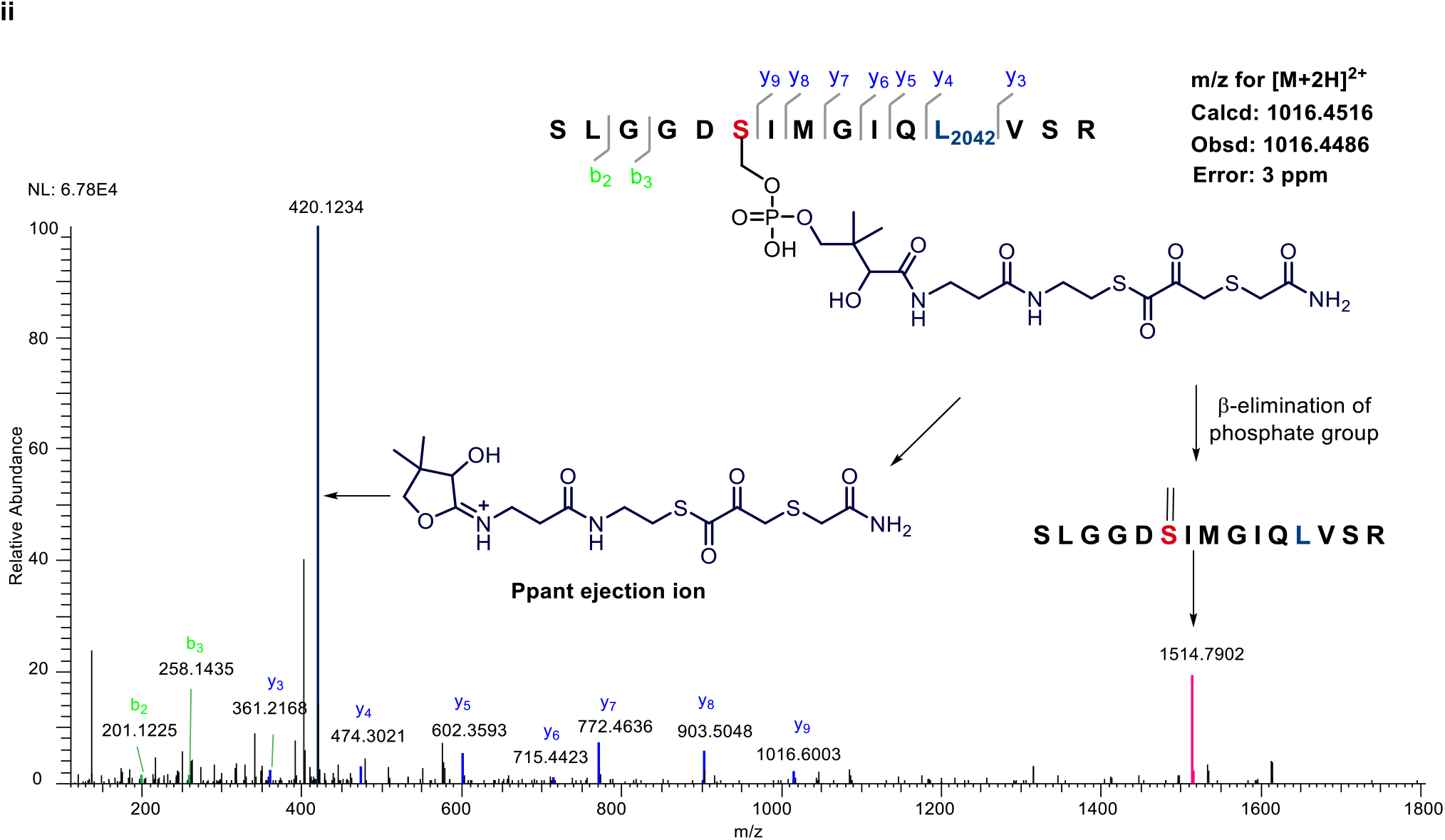
Characterization of (3-sulfhydryl)-pyruvoyl-*S*-CaeA2^F2042L^ (**6**) by nanoLC-MS/MS following treatment with IAM and subsequent complete digestion with trypsin and chymotrypsin. (a) Incubation of thiolated CaeA2^F2042L^, CaeB1 and L-cysteine. MS-detectable products include **4** (i) and **5** (ii).

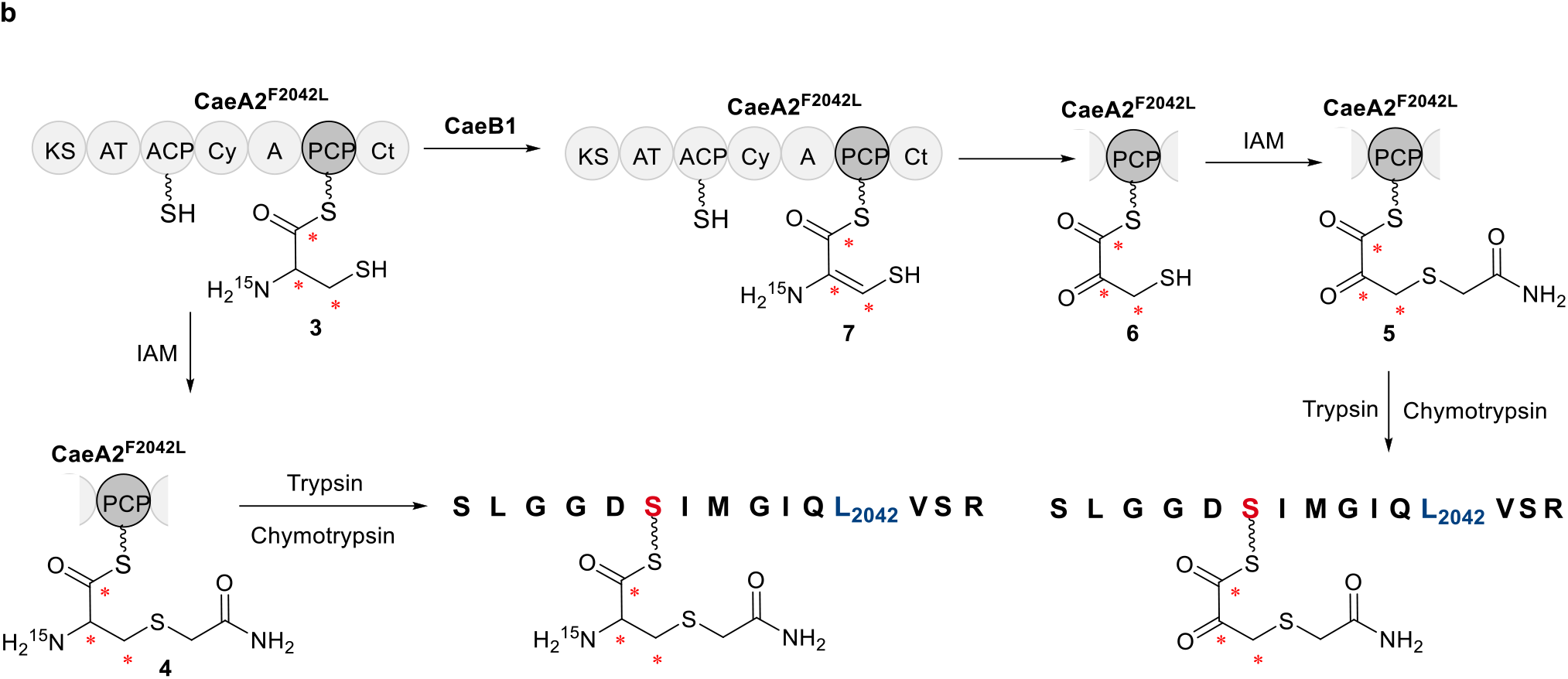
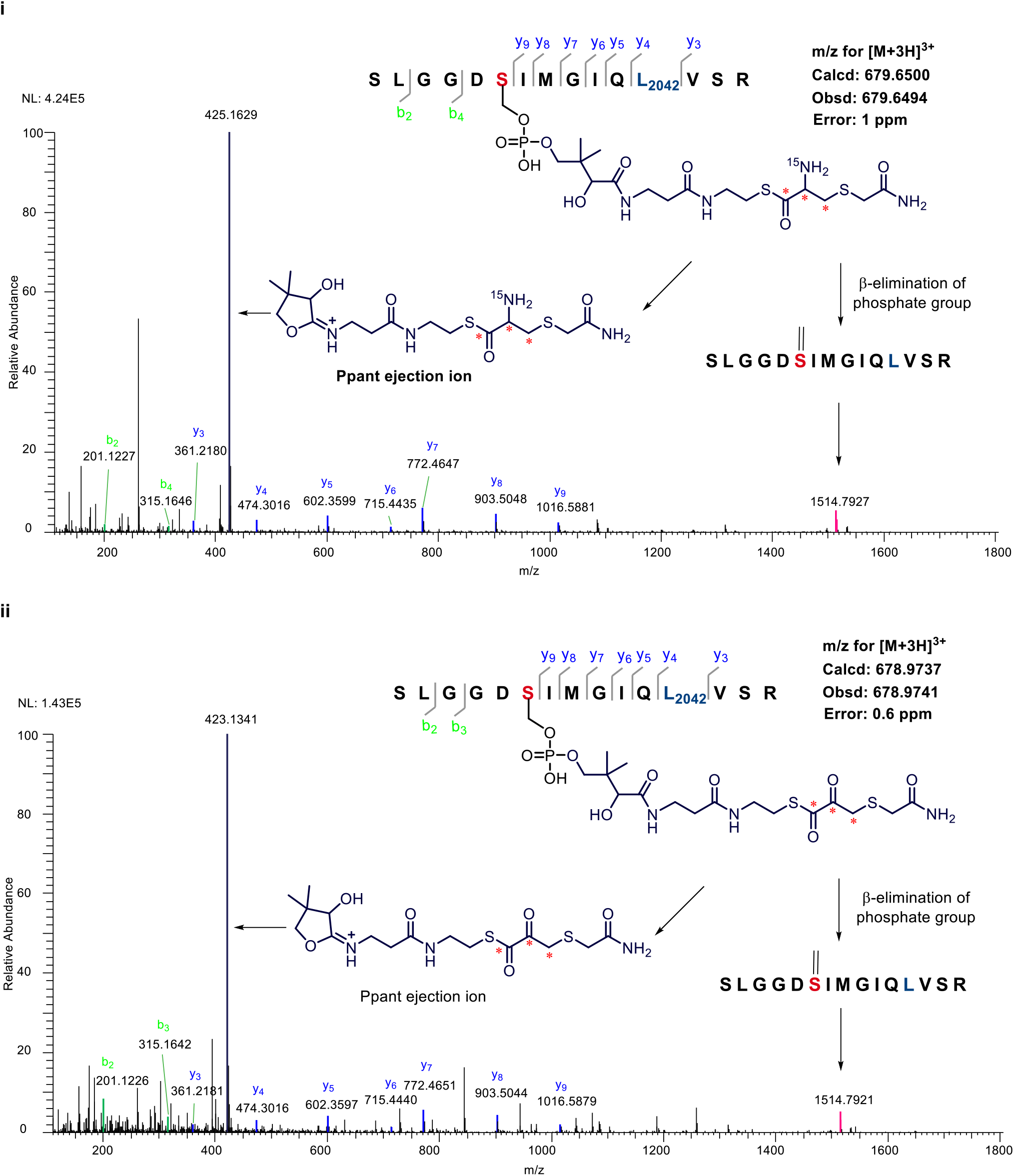
(b) Incubation of thiolated CaeA2^F2042L^, CaeB1 and L-[1,2,3-^13^C_3_,^15^N]cysteine. MS-detectable products include ^15^N and/or ^13^C labelled **4** (i) and **5** (ii).

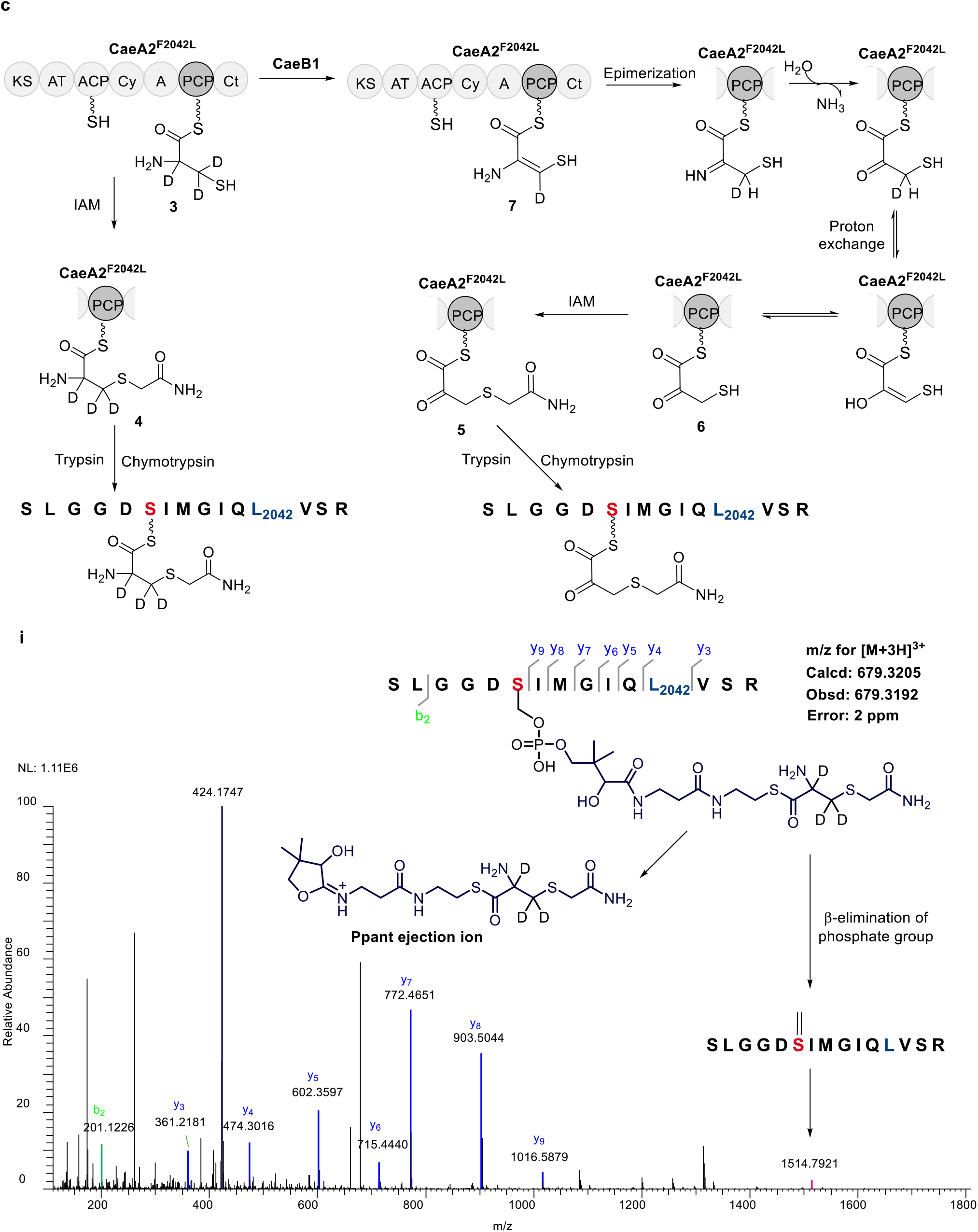
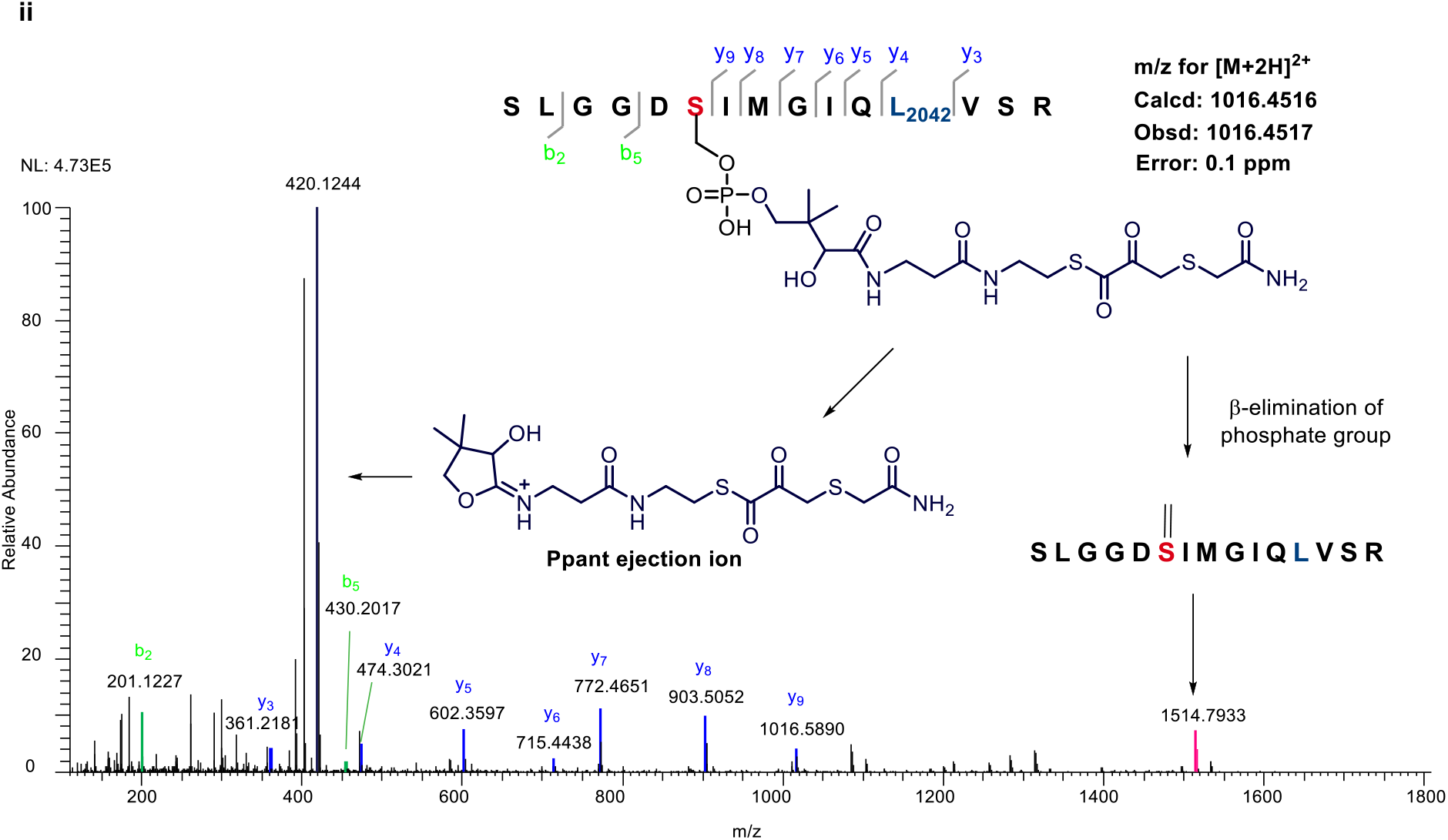
(c) Incubation of thiolated CaeA2^F2042L^, CaeB1 and L-[2,3,3-D_3_]cysteine. MS-detectable products include labelled **4** (i) and unlabeled **5** (ii).

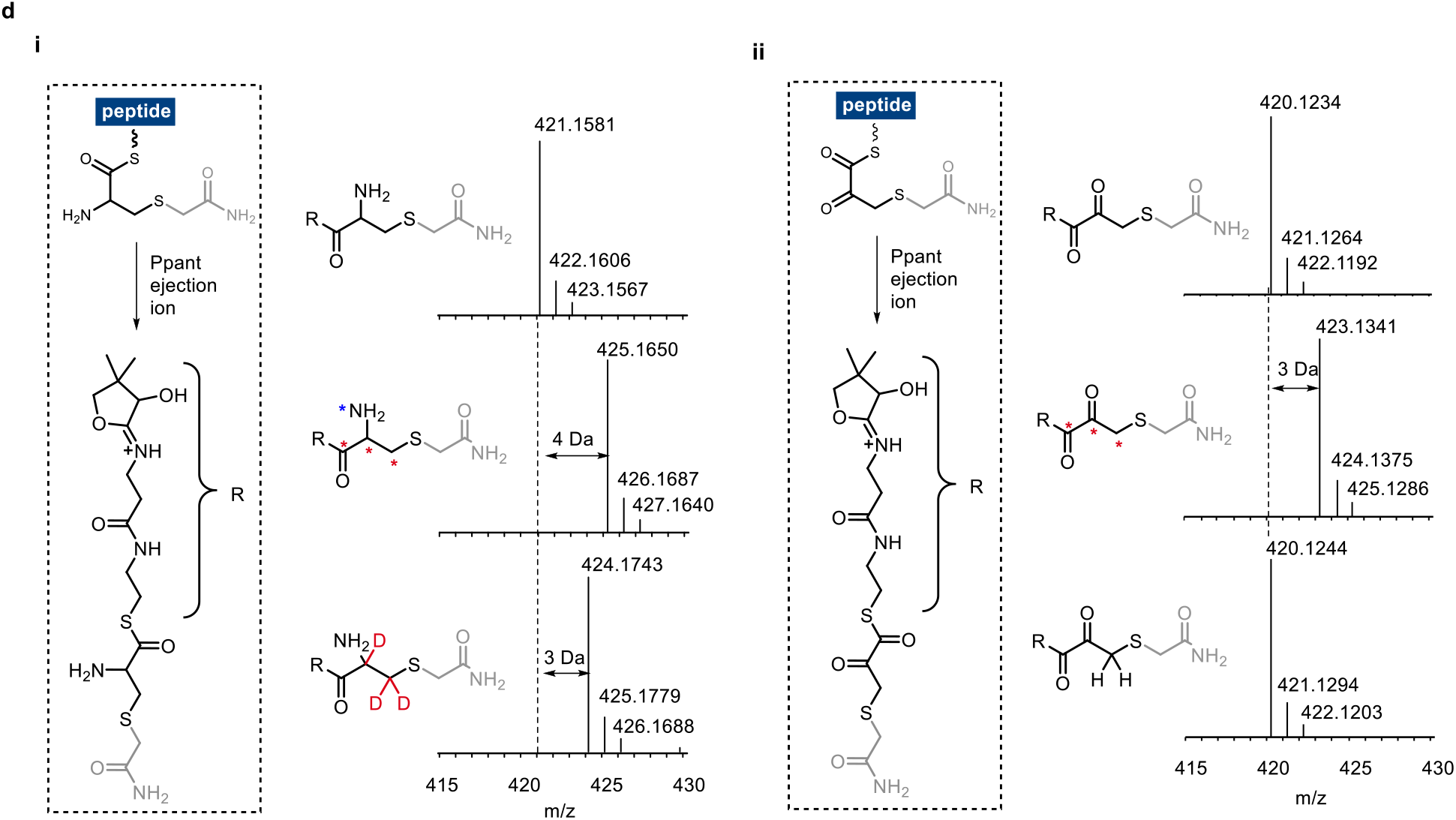
(d) MS comparison of **4** (i) and **5** (ii) in their associated Ppant ejection products, using L-cysteine (top), L-[1,2,3-^13^C_3_,^15^N]cysteine (middle) and L-[2,3,3-D_3_]cysteine (down) as the substrates, respectively.

**Supplementary Figure 7.**
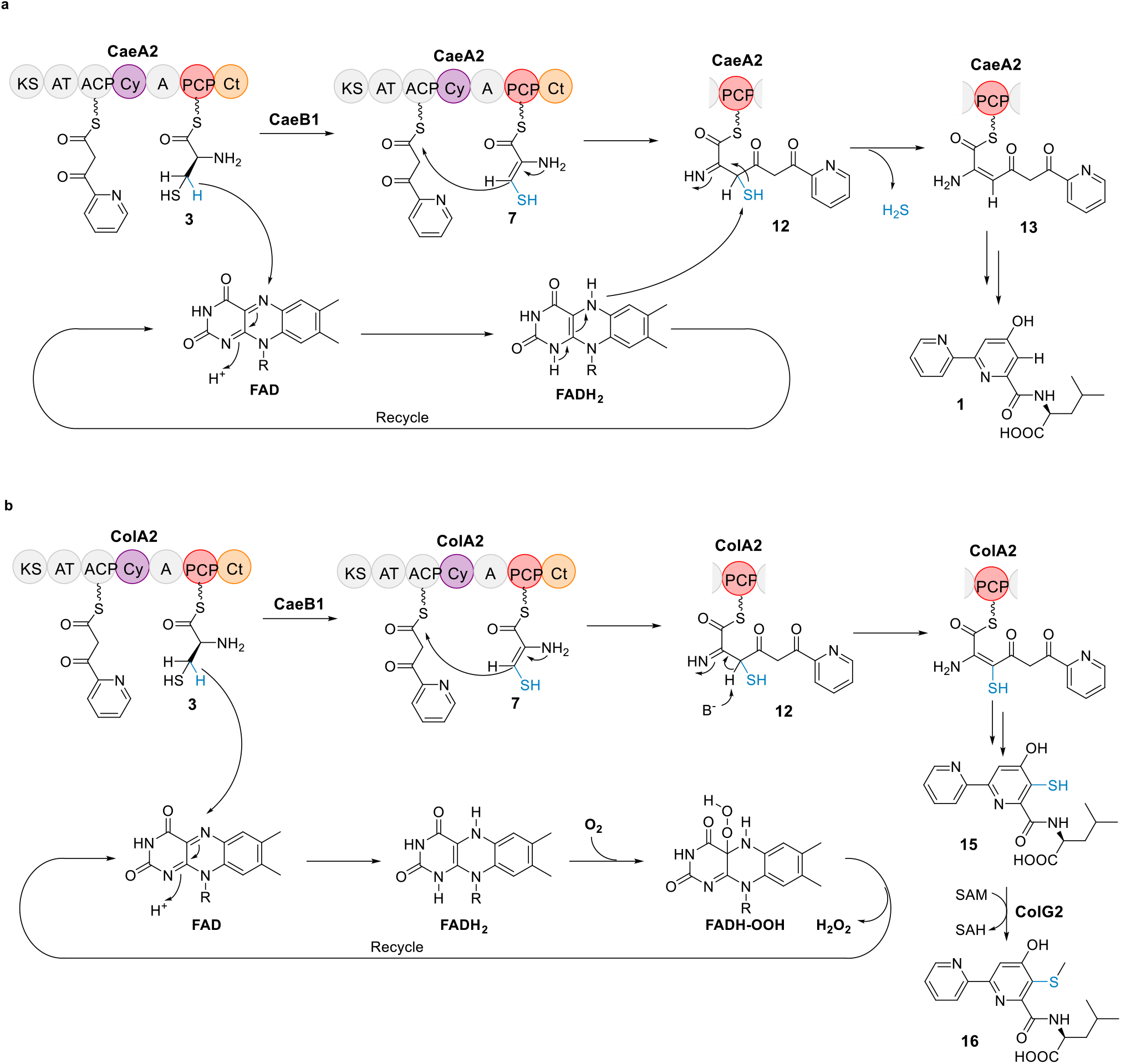
Proposed mechanisms for recycling FAD in the CAEs (**a**) and COLs (**b**) biosynthetic pathways, respectively.

**Supplementary Figure 8.**
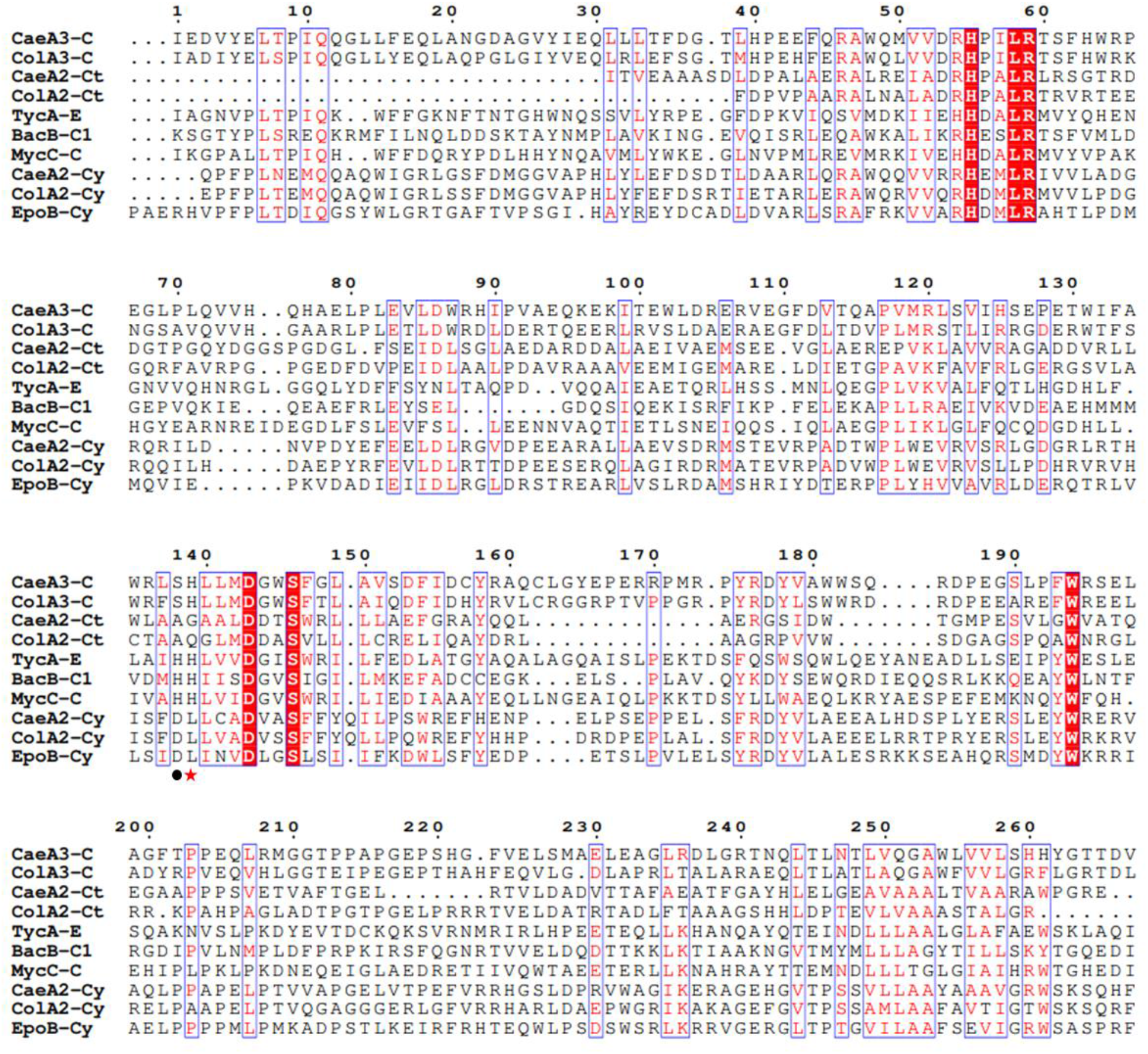
Sequence analysis of the C domains of CaeA3 and ColA3, and the Cy and Ct domains of CaeA2 and ColA2 For comparison, the homologs include the epimerization (E) domain of the NRPS TycA (610-913 aa, AAC45928.1) in tyrocidine biosynthesis^12^, the first C domain of the NRPS BacB (72-361 aa, AAC06347.1) in bacitracin biosynthesis^13^, the C domain of the NRPS MycC (855-1156 aa, AAF08797.1) in mycosubtilin biosynthesis^14^, and the Cy domain of the NRPS EpoB (71-364 aa, ADB12489.1) in epothilone biosynthesis^15^. The boundaries of each domain were determined by Pfam. The active site residue L-histine of C domains is indicated by red star, and the active site residue L-glutamic acid of Cy domains is indicated by black dot.

**Supplementary Figure 9.**
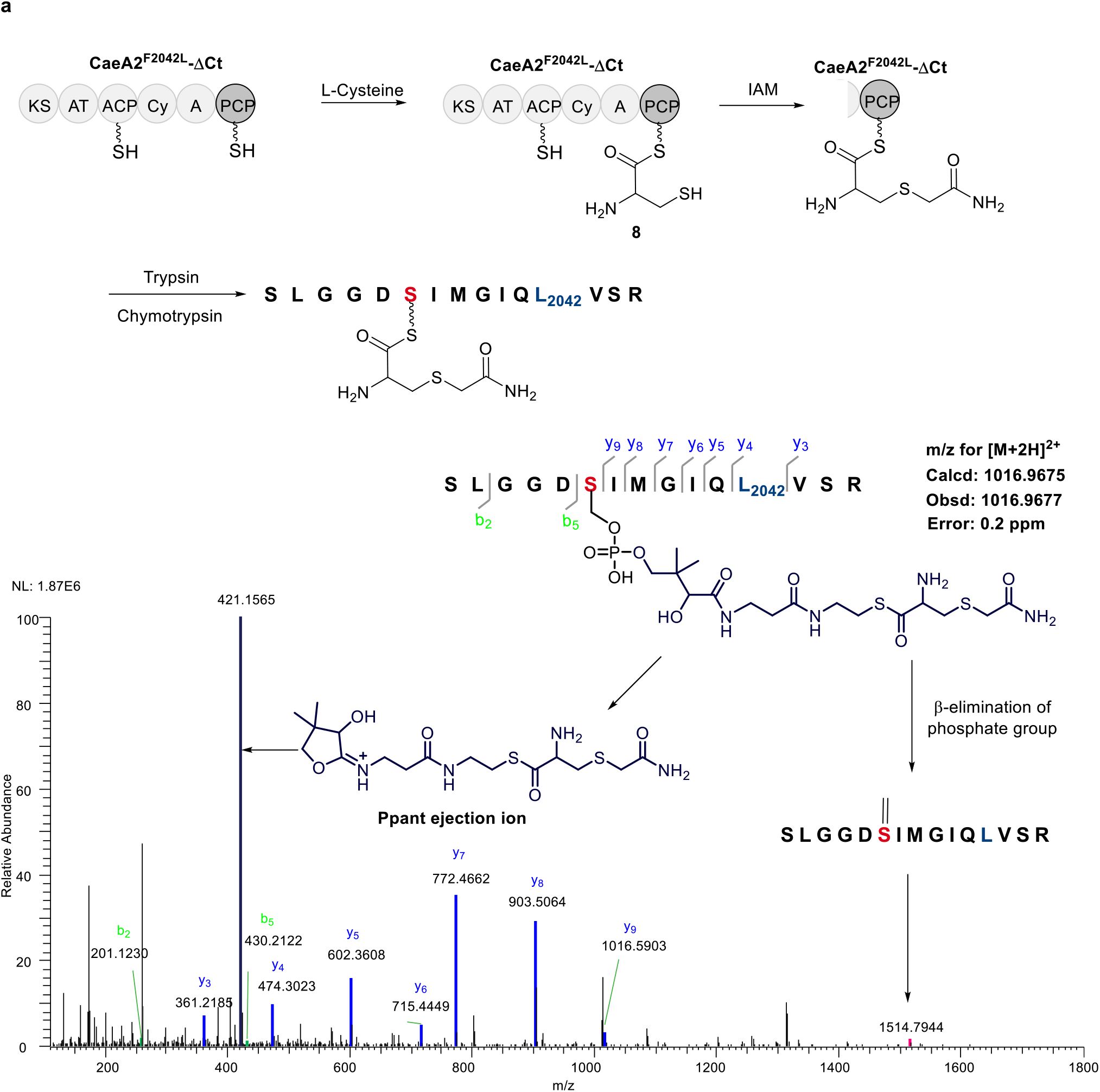
Characterization of L-Cysteinyl-*S*-CaeA2^F2042L^ΔCt (**8**) by nanoLC-MS/MS following treatment with IAM and subsequent complete digestion with trypsin and chymotrypsin. (a) Incubation of truncated CaeA2^F2042L^ΔCt and L-cysteine.

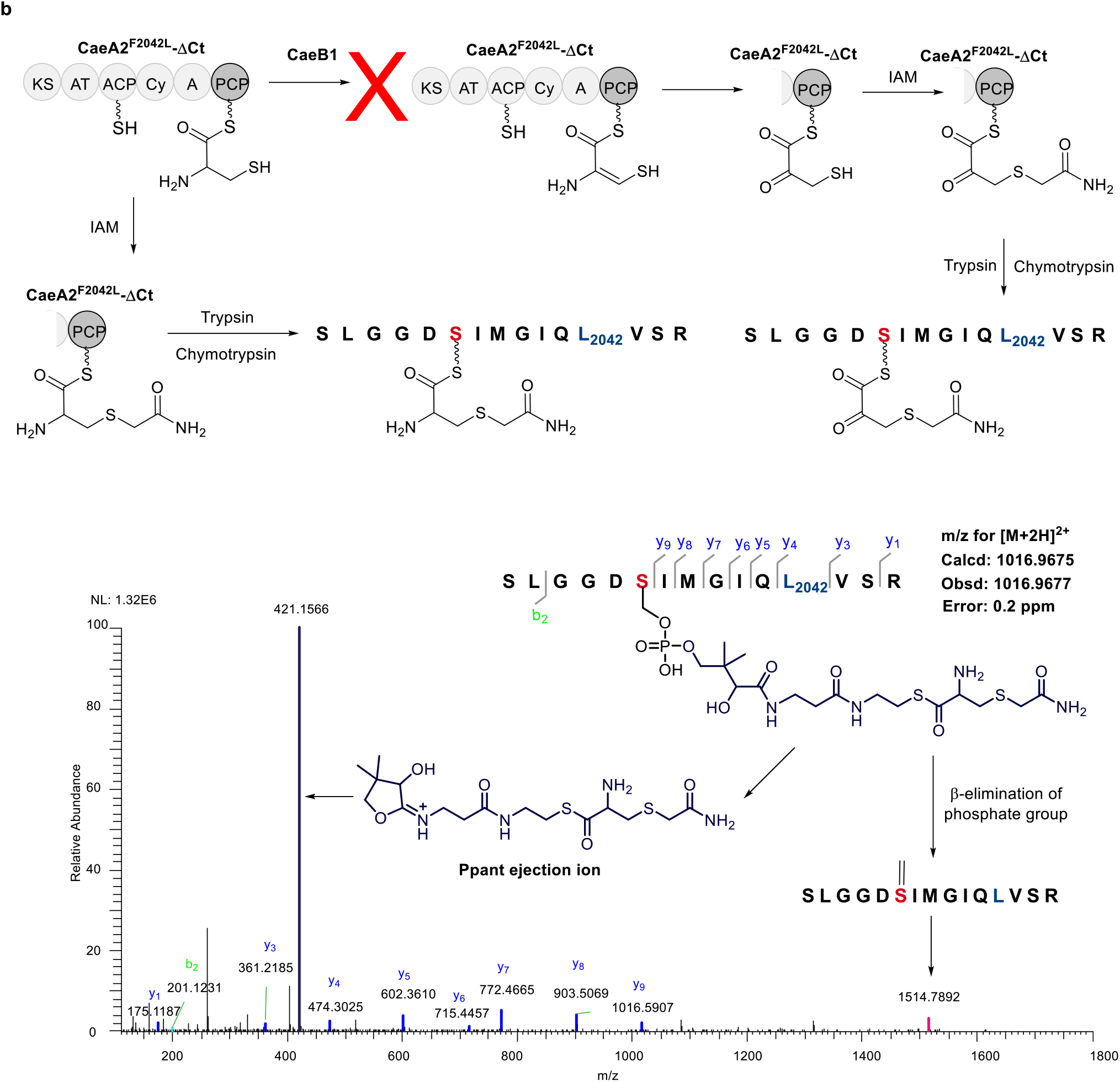
(b) Incubation of CaeB1 with truncated CaeA2^F2042L^ΔCt and L-cysteine.

**Supplementary Figure 10.**
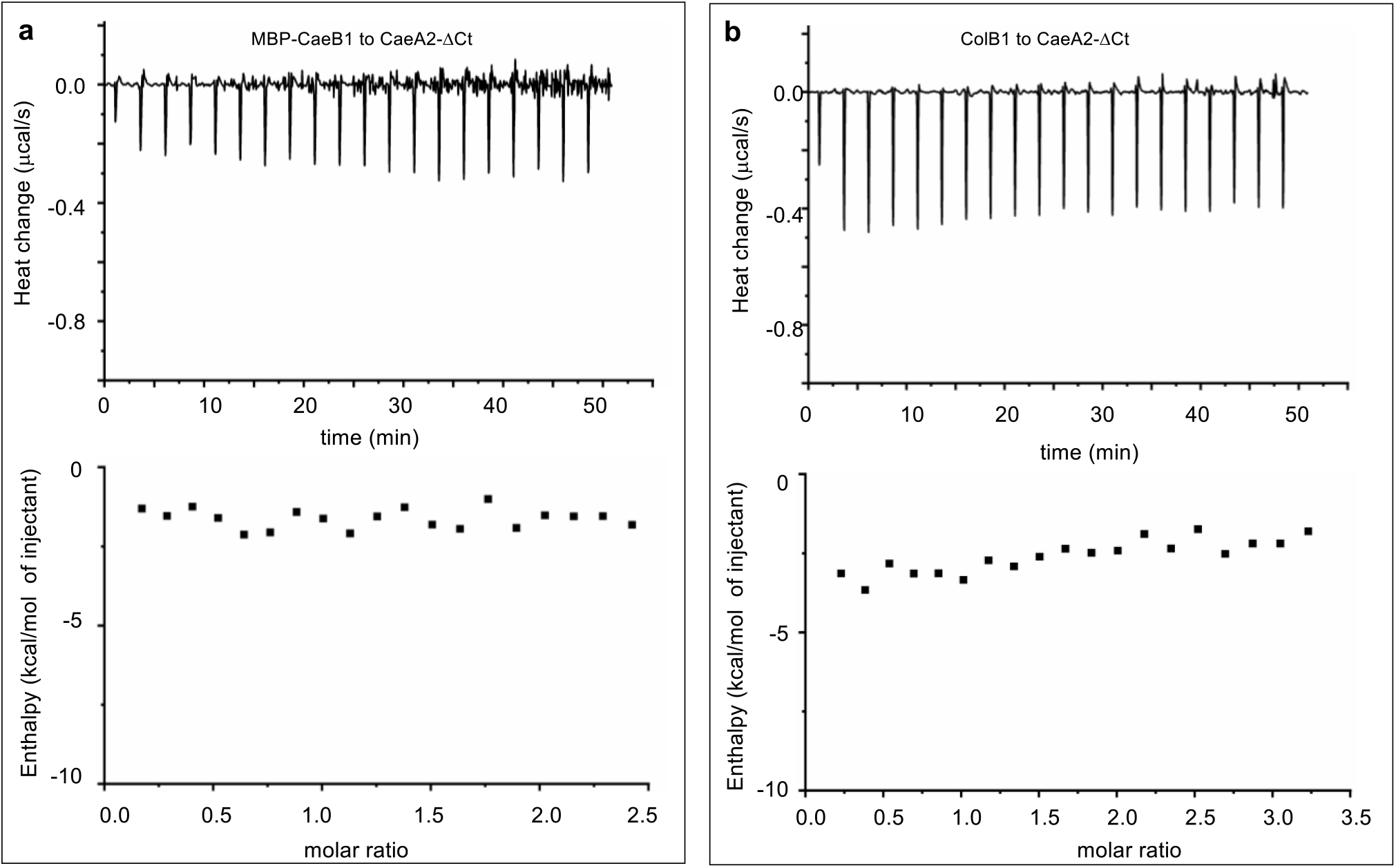
Measurement of the interactions of truncated CaeA2-ΔCt with related flavoproteins by ITC. Raw dates were shown on top, and the integrated curves containing experimental points and the best fitting line obtained from the single binding site model were shown on bottom. (**a**) Titrating MBP-fused CaeB1 to CaeA2-ΔCt. (**b**) Titrating ColB1 to CaeA2-ΔCt.

**Supplementary Figure 11.**
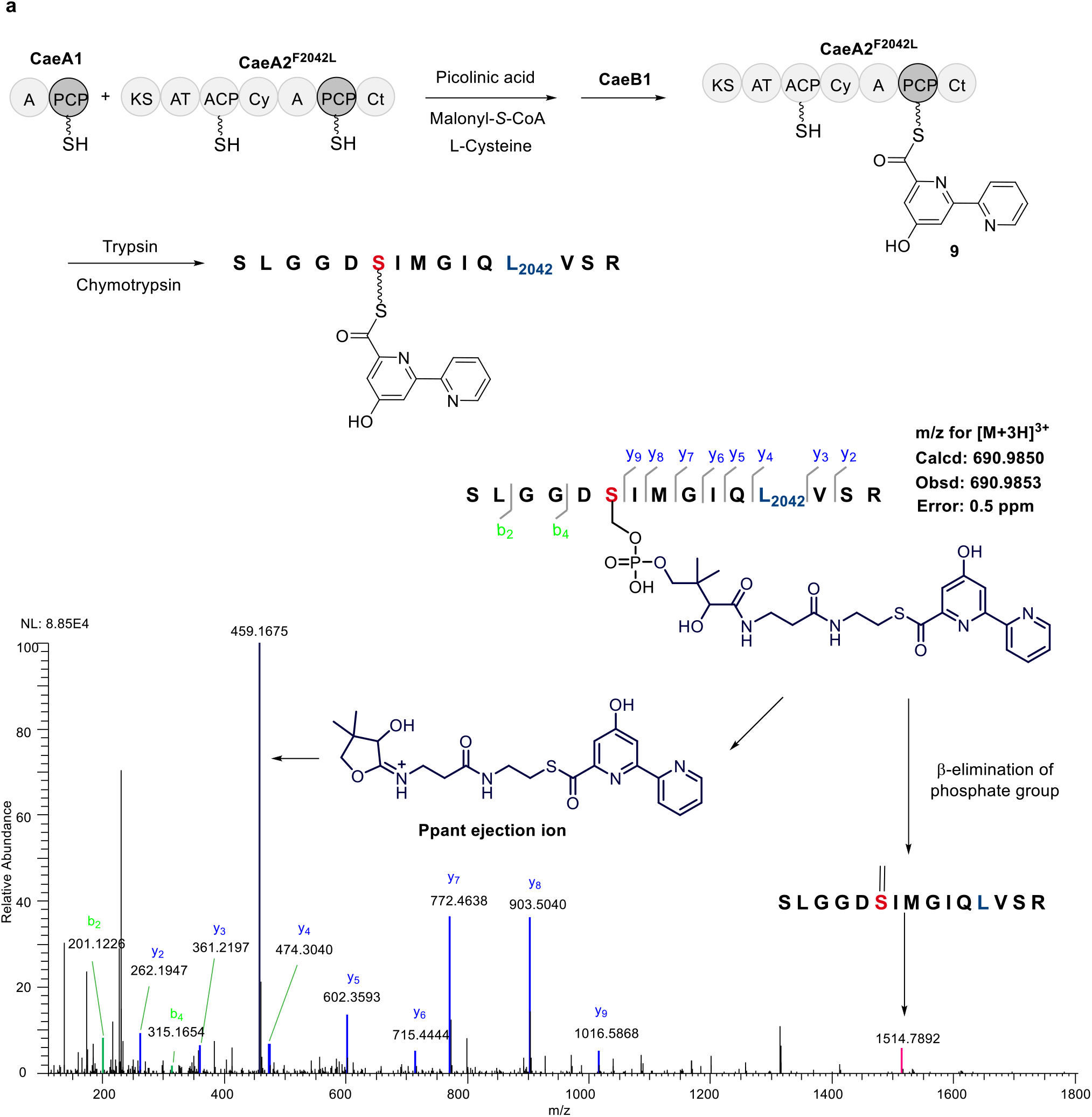
Characterization of 2,2’-bipyridinyl-*S*-CaeA2^F2042L^ (**9**) by nanoLC-MS/MS following complete digestion with trypsin and chymotrypsin. (a) Incubation of CaeA1, CaeA2^F2042L^ and CaeB1 with picolinic acid, malonyl-*S*-CoA and L-cysteine.

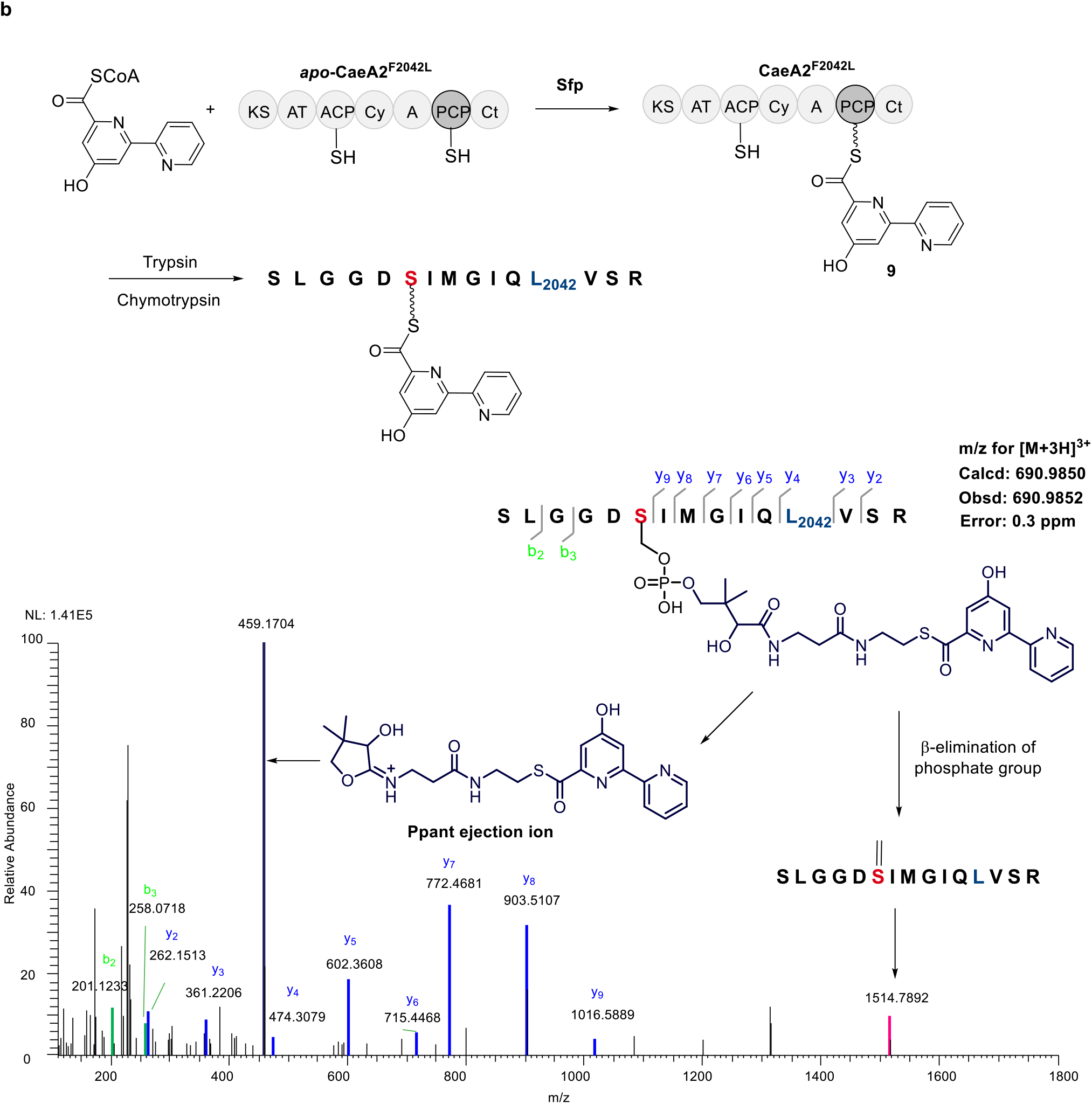
(b) Incubation of Ppant-unmodified CaeA2^F2042L^ (produced in *E. coli* BL21(DE3)) with synthesized 2,2’-bipyridinyl-*S*-CoA in the presence of Sfp.

**Supplementary Figure 12.**
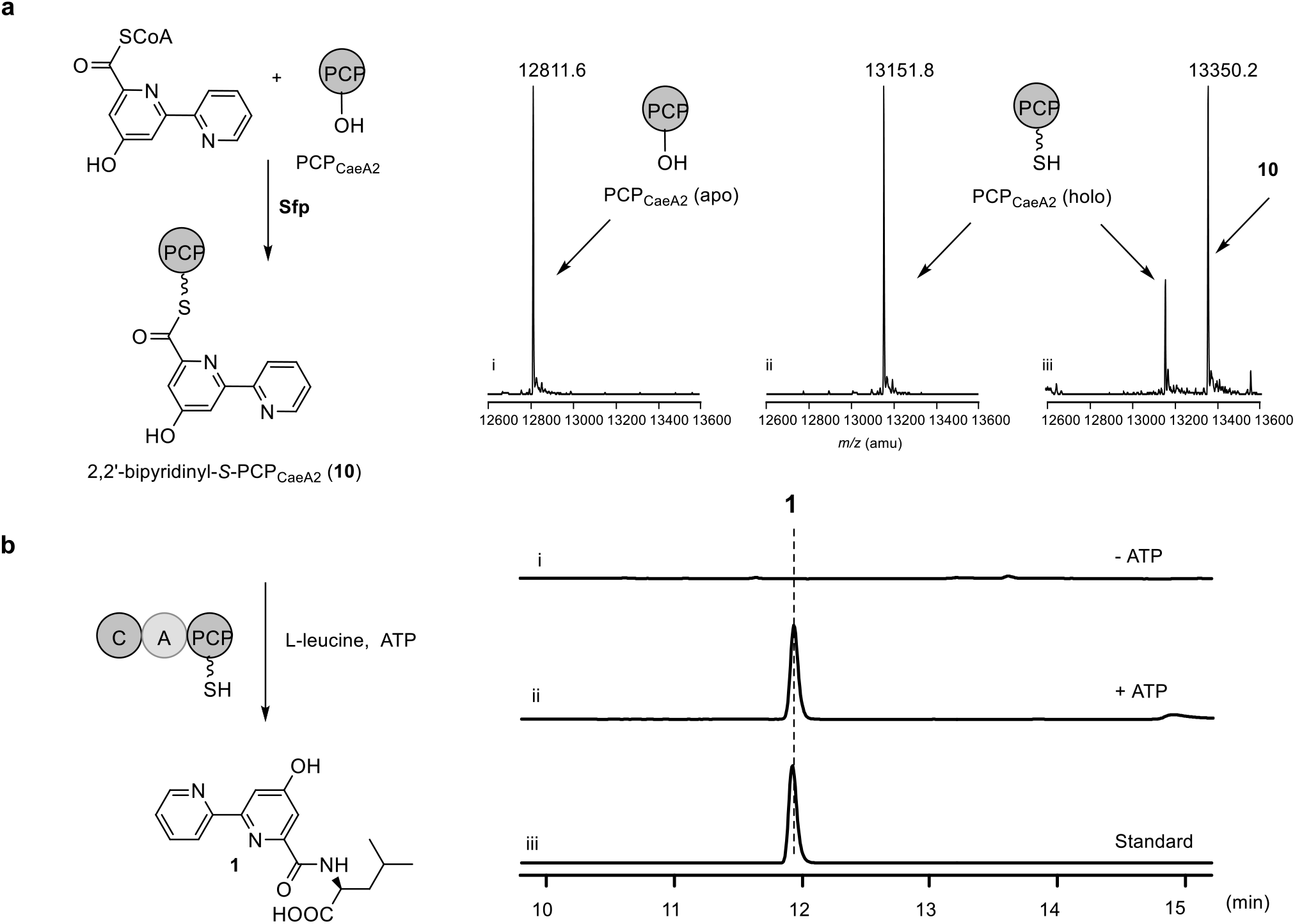
Validation of 2,2’-bipyridinyl-*S*-Ppant as an intermediate in the production of **1**. (**a**) Preparation of 2,2’-bipyridinyl-*S*-PCP_CaeA2_ (**10**) by the incubation of PCP_CaeA2_ in apo form (produced in *E. coli* BL21(DE3)) with synthesized 2,2’-bipyridinyl-*S*-CoA in the presence of Sfp. PCP_CaeA2_ in both apo (i) and holo (ii) forms and 2,2’-bipyridinyl-*S*-PCP_CaeA2_ (iii) were examined by HR-MS. (**b**) Incubation of prepared 2,2’-bipyridinyl-*S*-PCP_CaeA2_ with CaeA3 and L-leucine in the absence (i) and presence (ii) of ATP, the standard of **1** (iii).

**Supplementary Figure 13.**
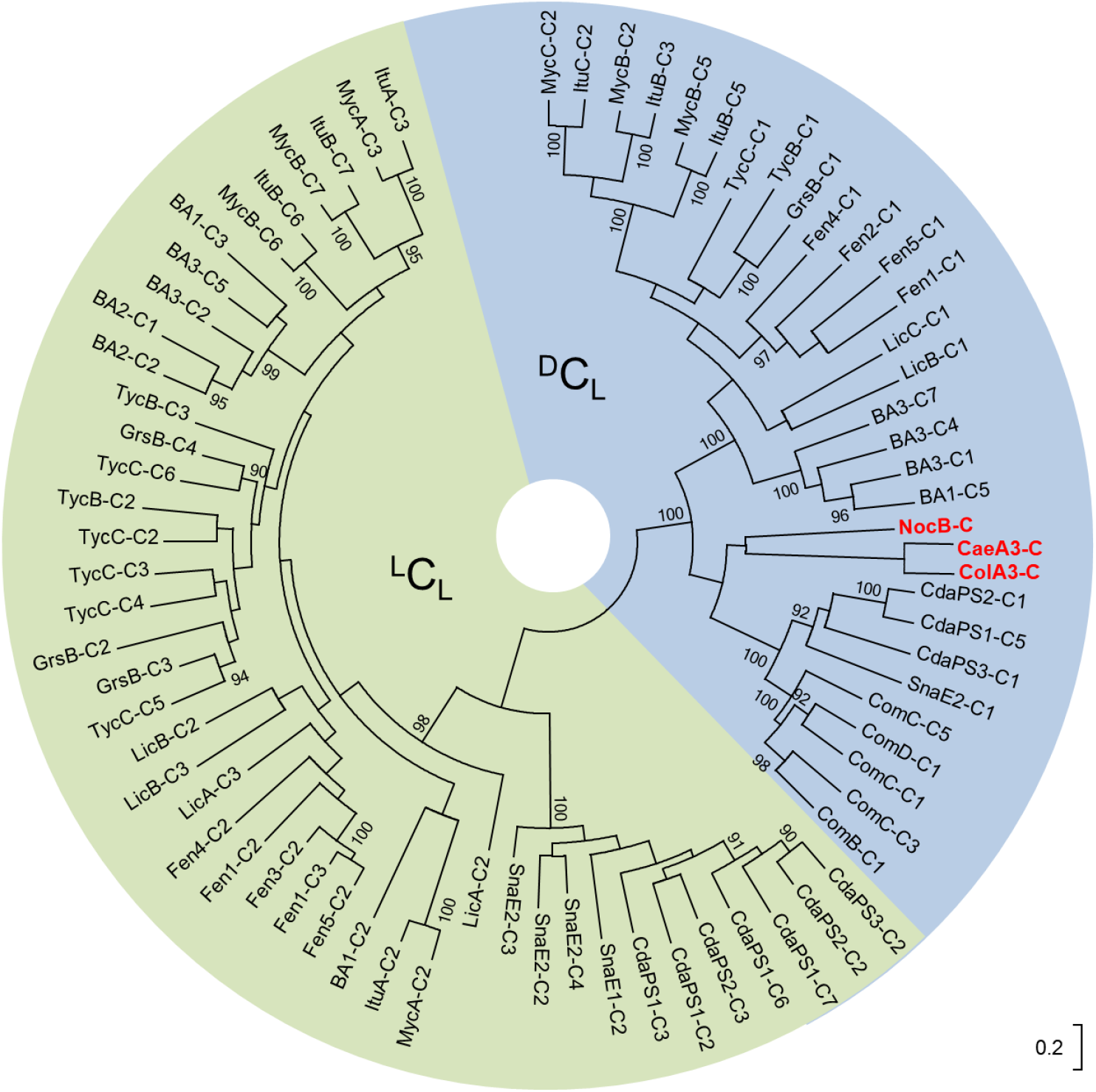
Phylogenetic analysis of the C domains of NRPS CaeA3 and ColA3 in substrate stereo-chemistry. The evolutionary distances were computed using the p-distance method. The support for grouping the clades ^L^C_L_ (green) and ^D^C_L_(blue) is indicated by bootstrap value. The C domains of CaeA3, ColA3 and the C domain of NocB that mediates β-lactam formation in nocardicin biosynthesis^16^ are shown in red. The homologous C domains arise from the NRPSs BA1, BA2 and BA3 in bacitracin biosynthesis^13^; CdaPS1, CdaPS2 and CdaPS3 in calcium-dependent antibiotic biosynthesis^17^; ComB, ComC and ComD in complestatin biosynthesis^18^; Fen1, Fen2, Fen3, Fen4 and Fen5 in fengycin bioysynthesis^19^; GrsB in gramicidin biosynthesis^20^; ItuA, ItuB and ItuC in iturin biosynthesis^21^; LicA, LicB and LicC in lichenicin biosynthesis^22^; MycA, MycB and MycC in mycosubtilin biosynthesis^23^; SnaE1 and SnaE2 in pristinamycin biosynthesis^24^; and TycB and TycC in tyrocidine biosynthesis^12^. The sequences were downloaded from the database of NaPDoS, in which the biochemical function and substrate stereo-chemistry of these related C domains have been confirmed^25,26^.

**Supplementary Figure 14.**
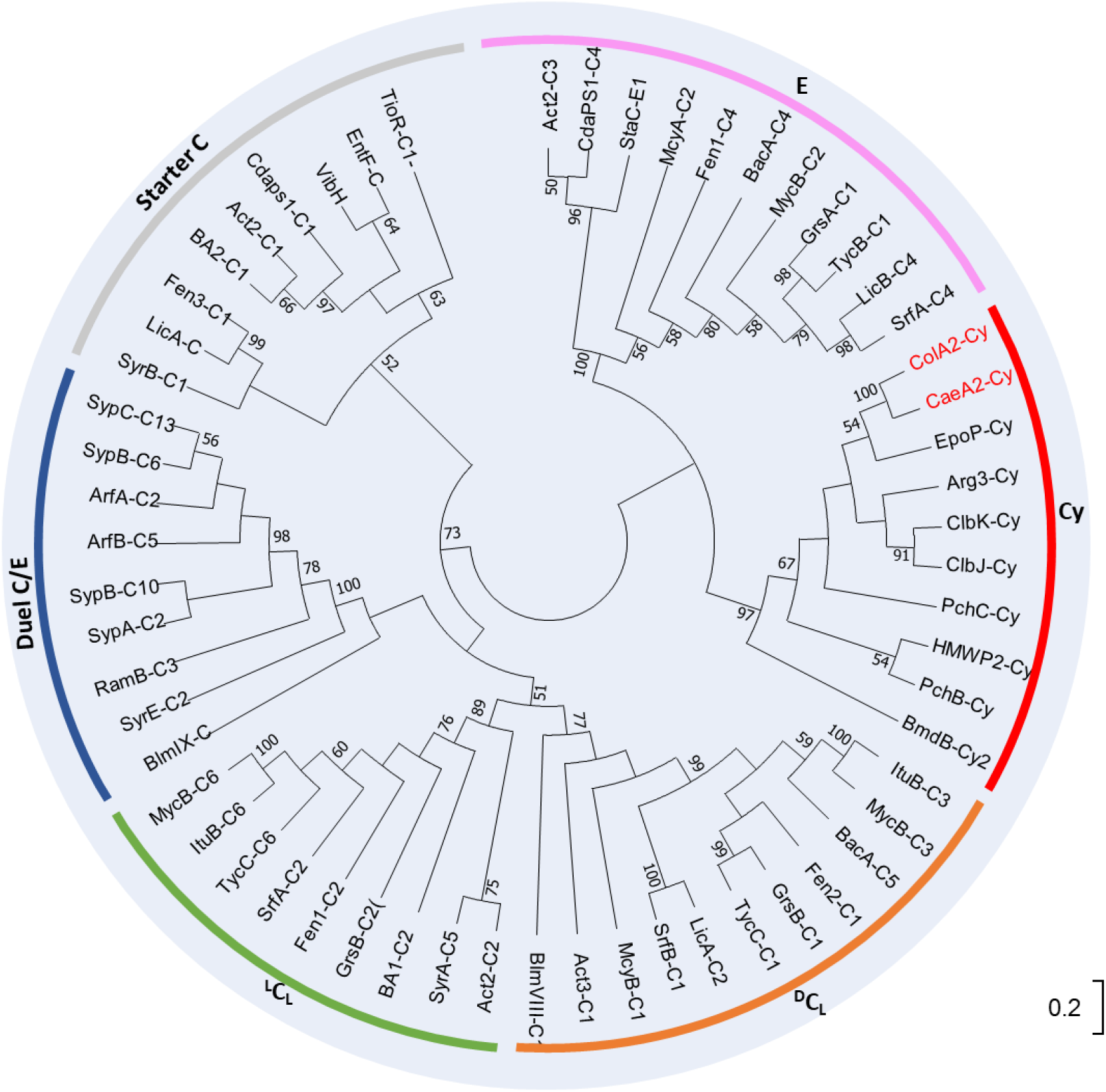
Phylogenetic analysis of the Cy domains of the NRPSs CaeA2 and ColA2. The evolutionary distances were computed using the p-distance method. The support for grouping the C domain subtypes is indicated by bootstrap value. The Cy domains of CaeA2 and ColA2 are shown in red. Members of the C domain subtypes (^L^C_L_, ^D^C_L_, Starter C, Epimerization (E), Duel E/C and Heterocyclization (Cy) domains) include those of the NRPSs Act2 and Act3 in actinomycin biosynthesis^27^; ArfA and ArfB in arthrofactin biosynthesis^28^, Arg3 in argyrins biosynthesis^29^; BA1 and BA2 in bacitracin biosynthesis^13^; BmdB in bacillamide biosynthesis^30^; BlmVIII and BlmIV in bleomycin biosynthesis^31^; ClbJ and ClbK in colibactin biosynthesis^32^; CdaPS1 in calcium-dependent antibiotic biosynthesis^17^; EpoP in epothilone biosynthesis^15^; EntF in enterobactin biosynthesis^33^; Fen1, Fen2 and Fen3 in fengycin bioysynthesis^19^; GrsA and GrsB in gramicidin biosynthesis^20^; HMWP2 in yersiniabactin biosynthesis^34^; ItuB in iturin biosynthesis^21^; LicA and LicB in lichenicin biosynthesis^22^; McyA and McyB in microcystin biosynthesis^35^; MycB in mycosubtilin biosynthesis^23^; PchB and PchC in pyochelin biosynthesis^36^; RamB in ramoplanin biosynthesis^37^; SrfA and SrfB in surfactin biosynthesis^38^; SypA, SypB and SypC in syringopeptin biosynthesis^39^; StaC in A47934 biosynthesis^40^; SyrA, SyrB and SyrE in syringomycin biosynthesis^41^; TycB and TycC in tyrocidin biosynthesis^12^; TioR in thiocoraline biosynthesis^42^; VibH in vibriobactin biosynthesis^43^. The sequences were downloaded from the database of NaPDoS, in which the biochemical function of these related C domains have been characterized^25,26^.

**Supplementary Figure 15.**
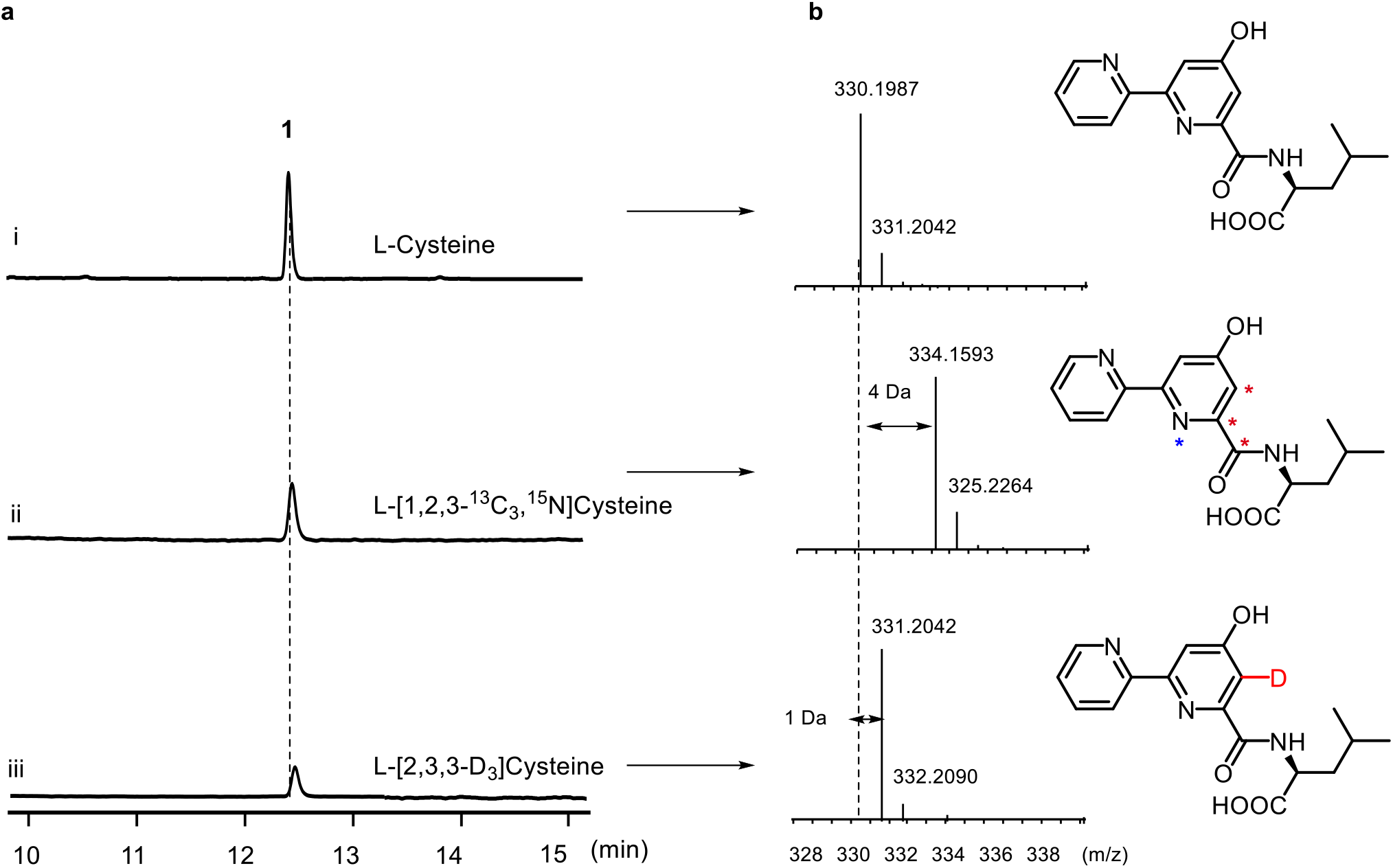
Comparison in the production of the CAE 2,2’-bipyridine intermediate **1** using the substrate L-cysteine (i), L-[1,2,3-^13^C_3_,^15^N]cysteine (ii) and L-[2,3,3-D_3_]cysteine (iii), respectively. (**a**) HPLC analysis. (**b**) HR-MS analysis.

**Supplementary Figure 16.**
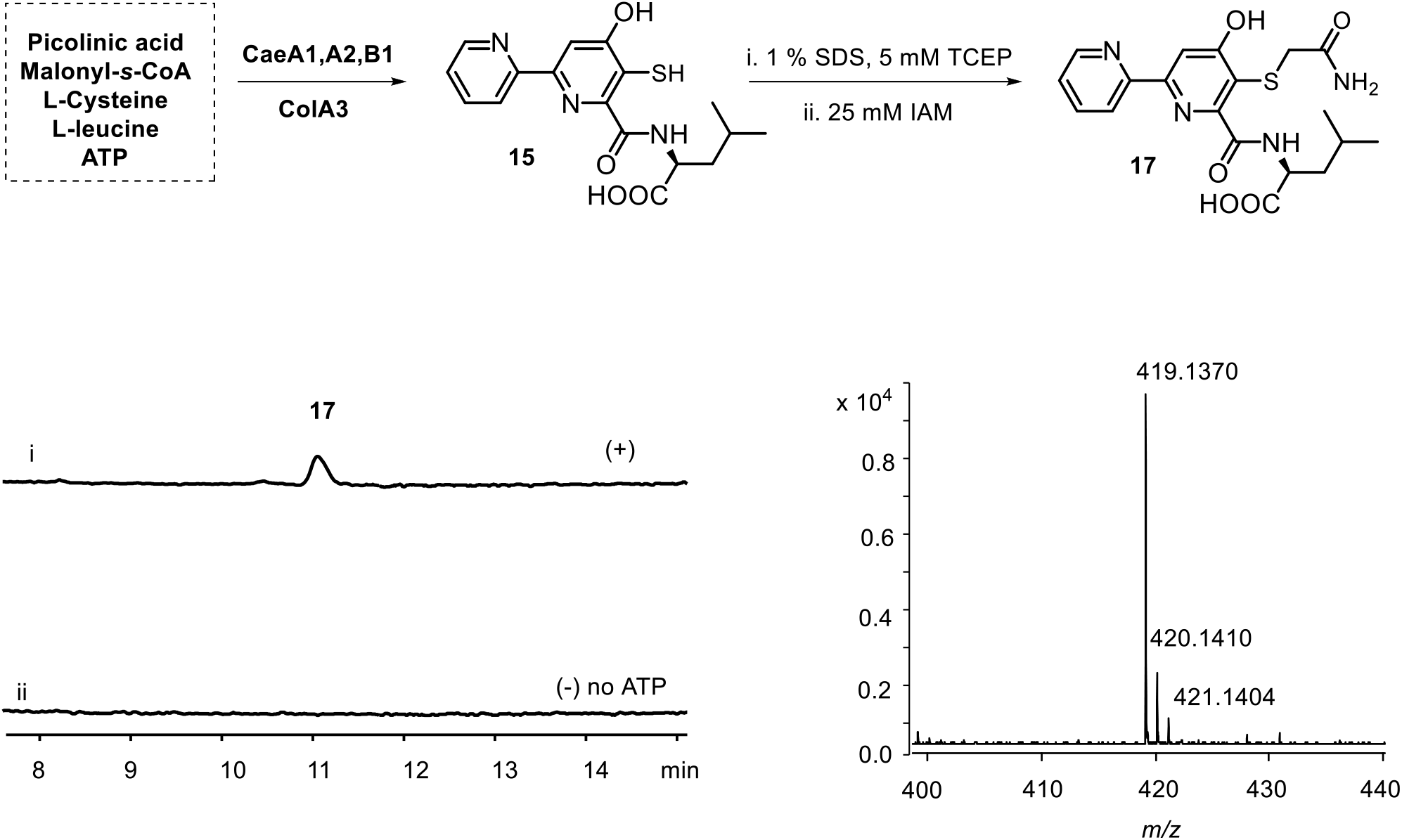
Examination of thiol intermediate 5-sulfhydryl-2,2’-bipyridinyl-L-leucine (**15**) by IAM derivatization. **15** was produced in the reactions where CaeA1, CaeA2, CaeB1and ColA3 were combined with picolinic acid, malonyl-*S*-CoA, L-cysteine and L-leucine in the presence (i) or absence (ii) of ATP. The reaction mixtures were treated with IAM to produce **17**, which were then analyzed by HPLC (left, λ = 315 nm) and HR-MS (right, [M + H]^+^ *m/z*: cald. 419.1389 for C_19_H_22_O_5_N_4_S, obs. 419.1370).

## SUPPLEMENTARY TABLES

**Table S1.**
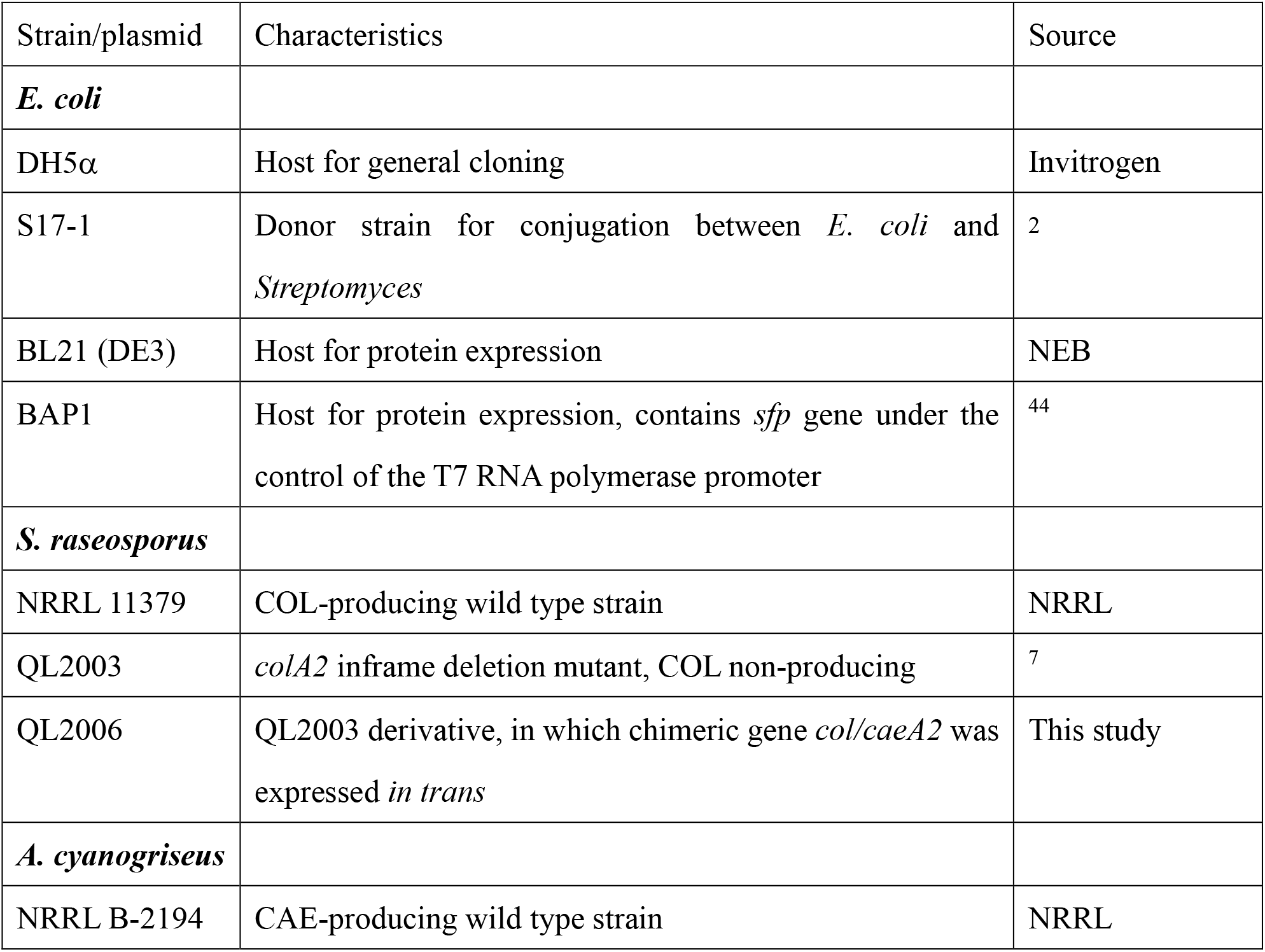
Bacterial strains in this study.

**Table S2.**
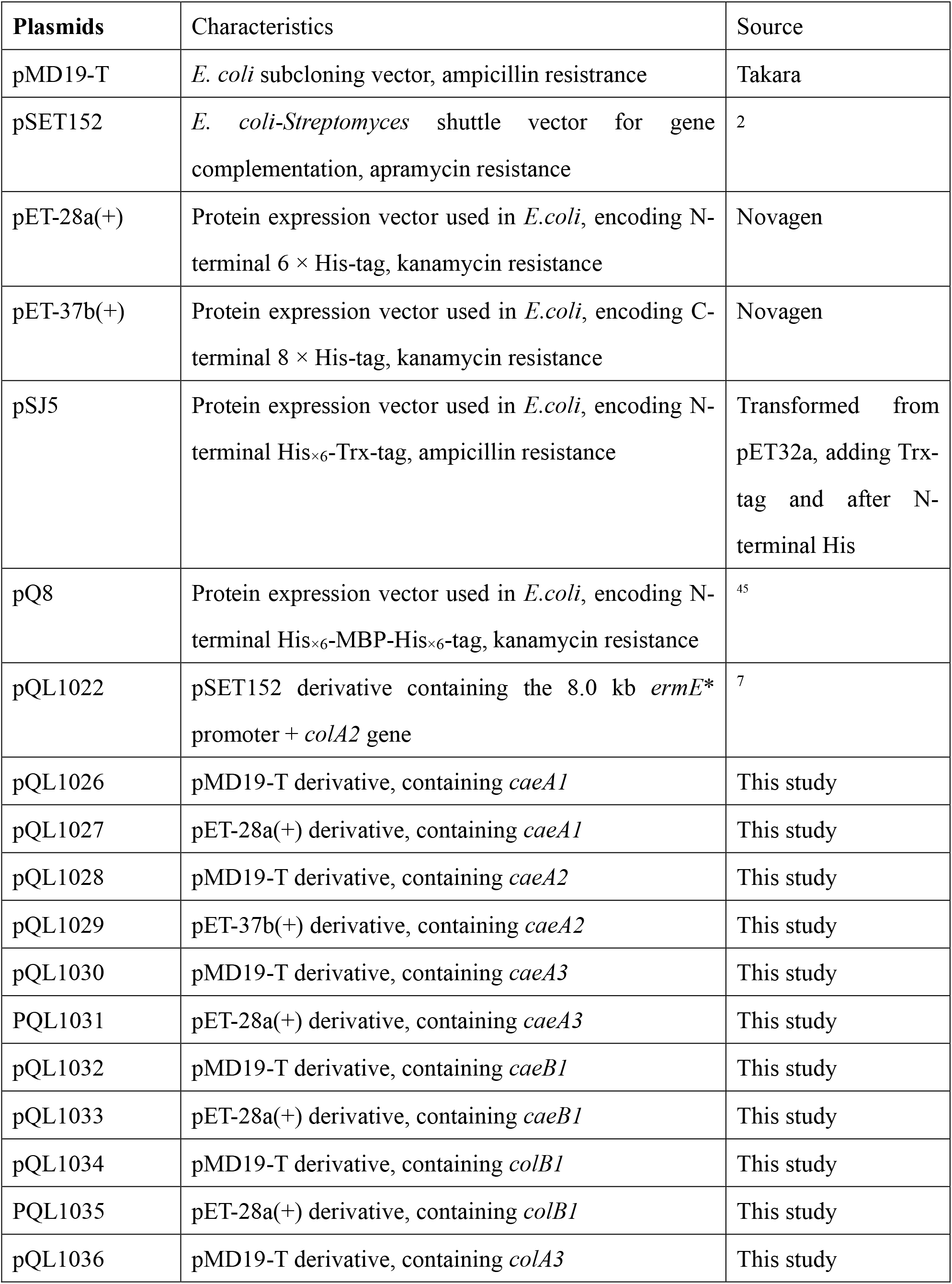
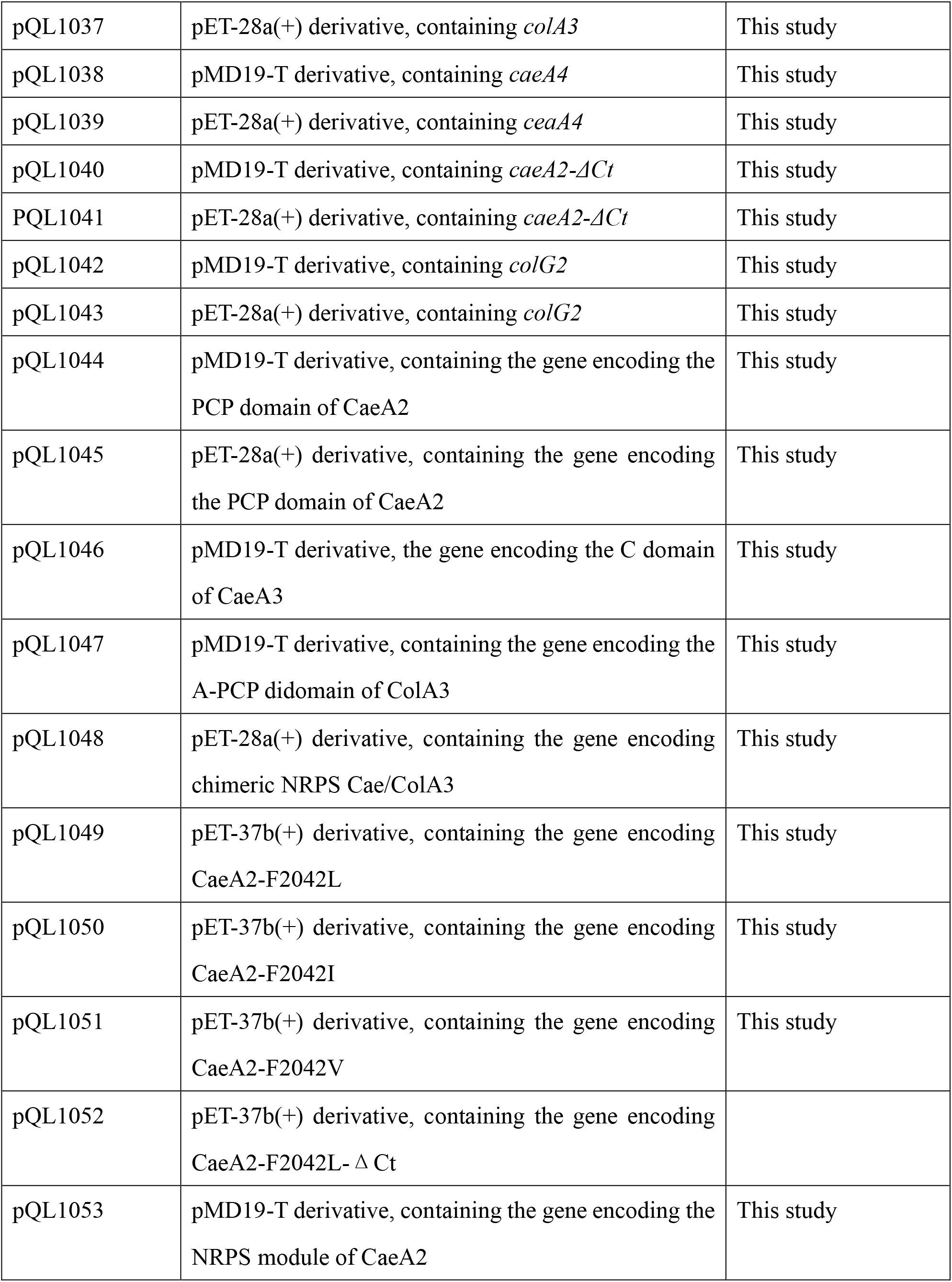
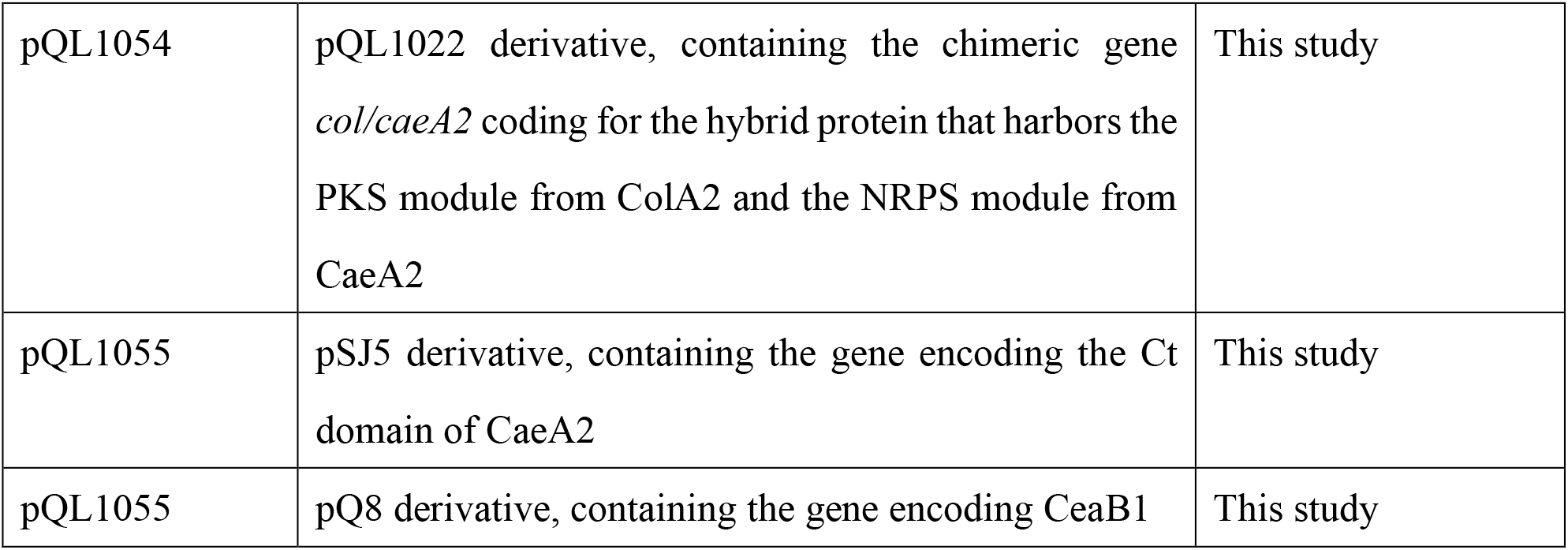
Plasmids in this study.

**Table S3.**
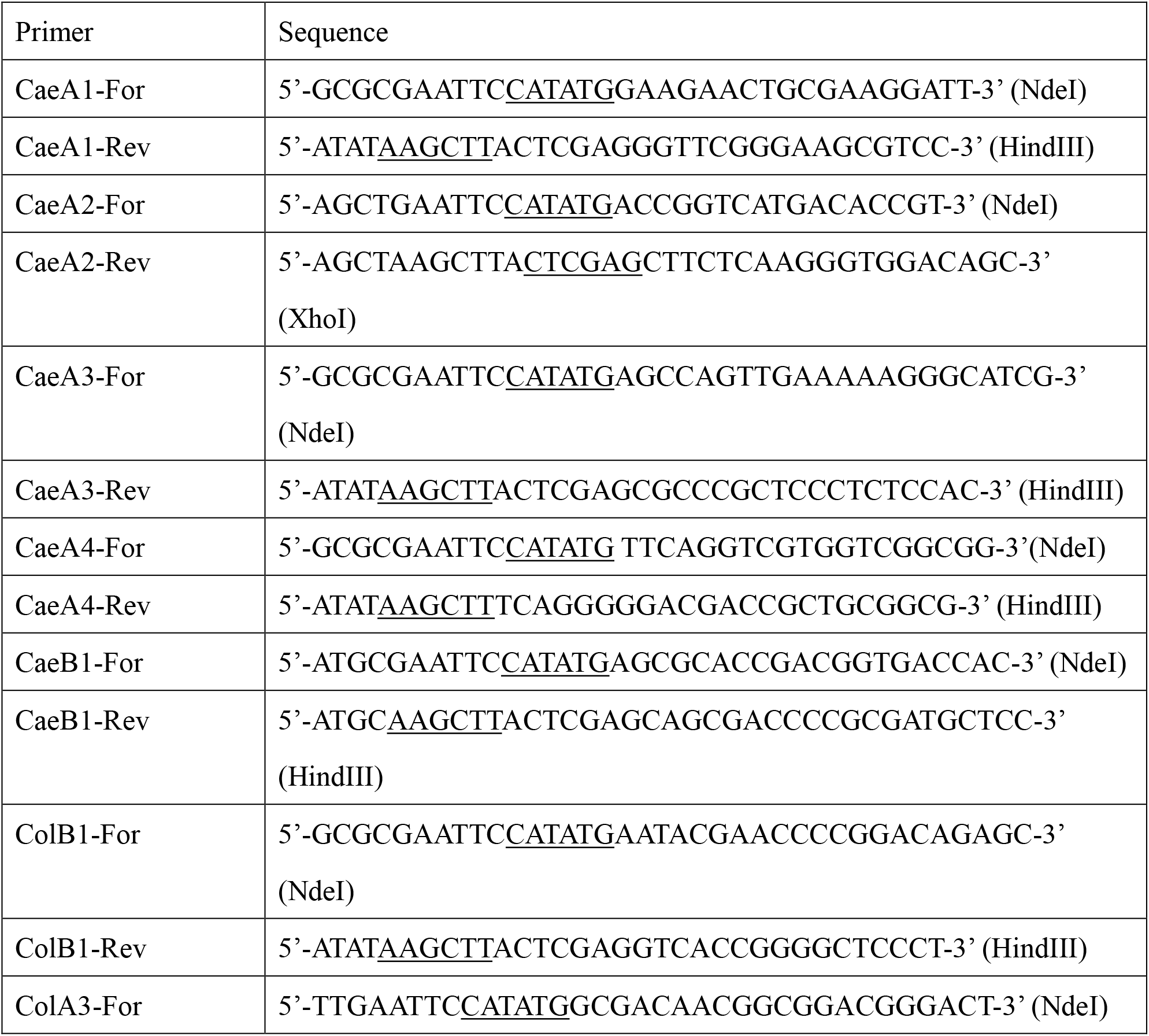
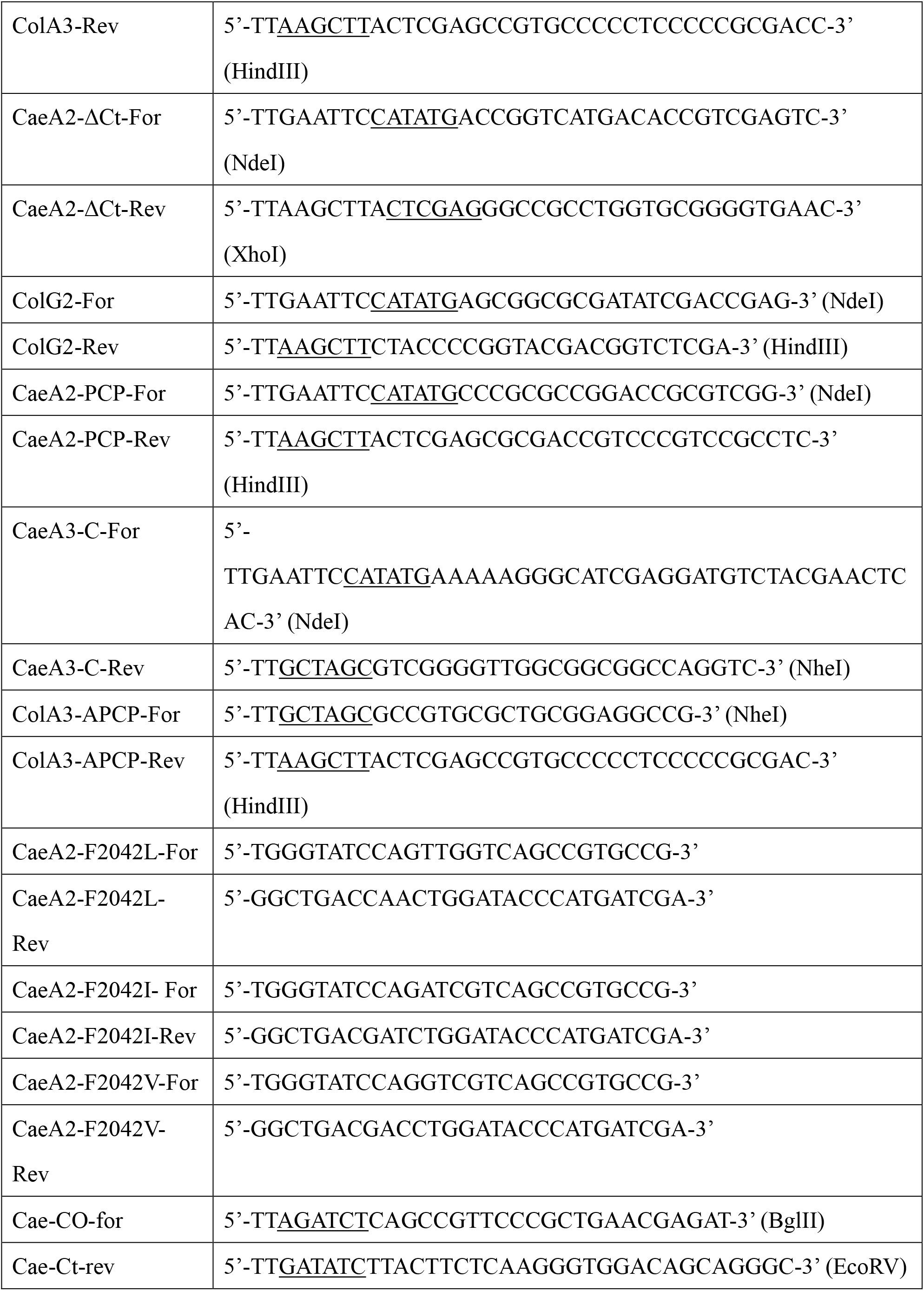
Primers used in this study. The sequences of restriction enzymes are underlined.

**Table S4.**
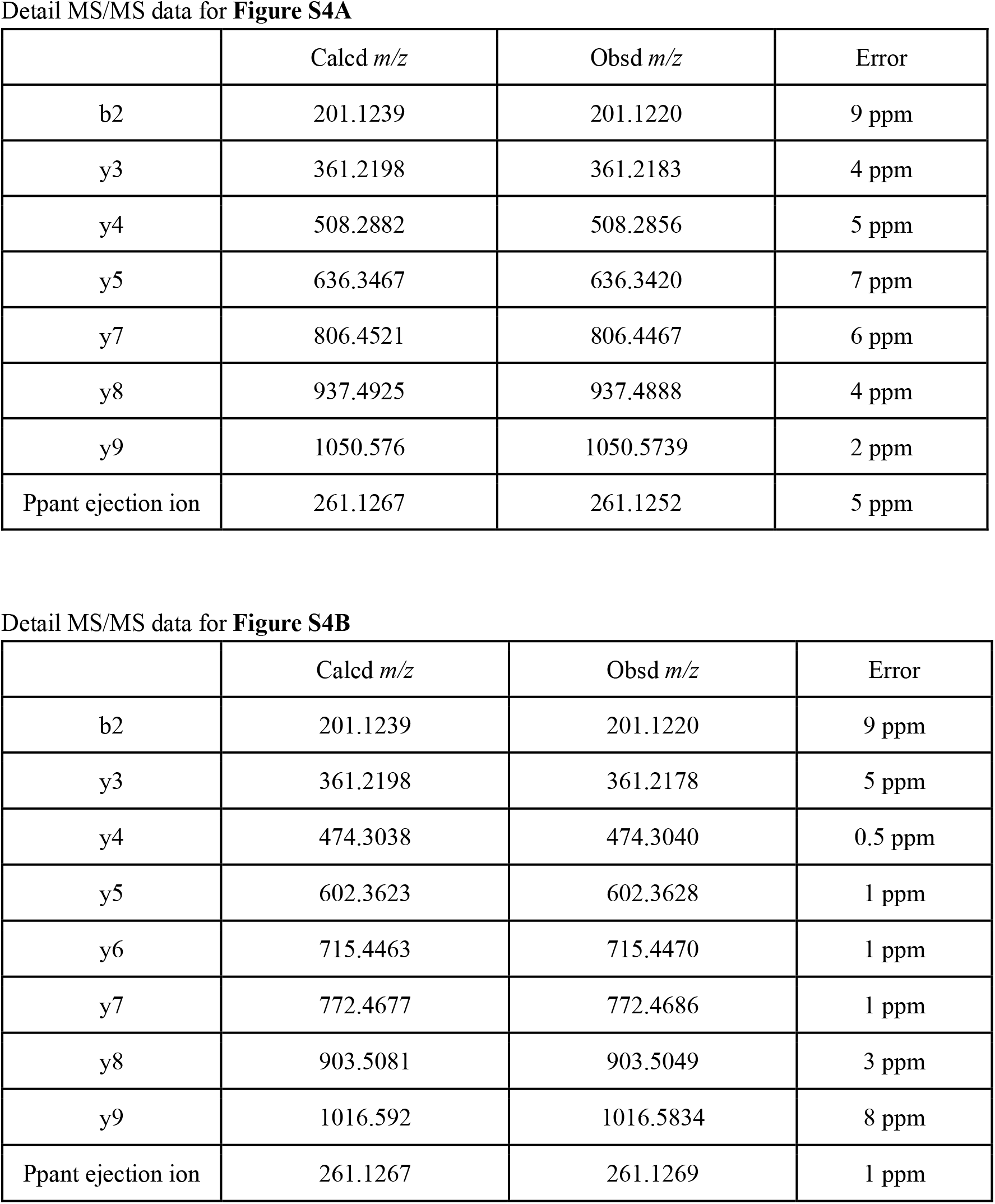
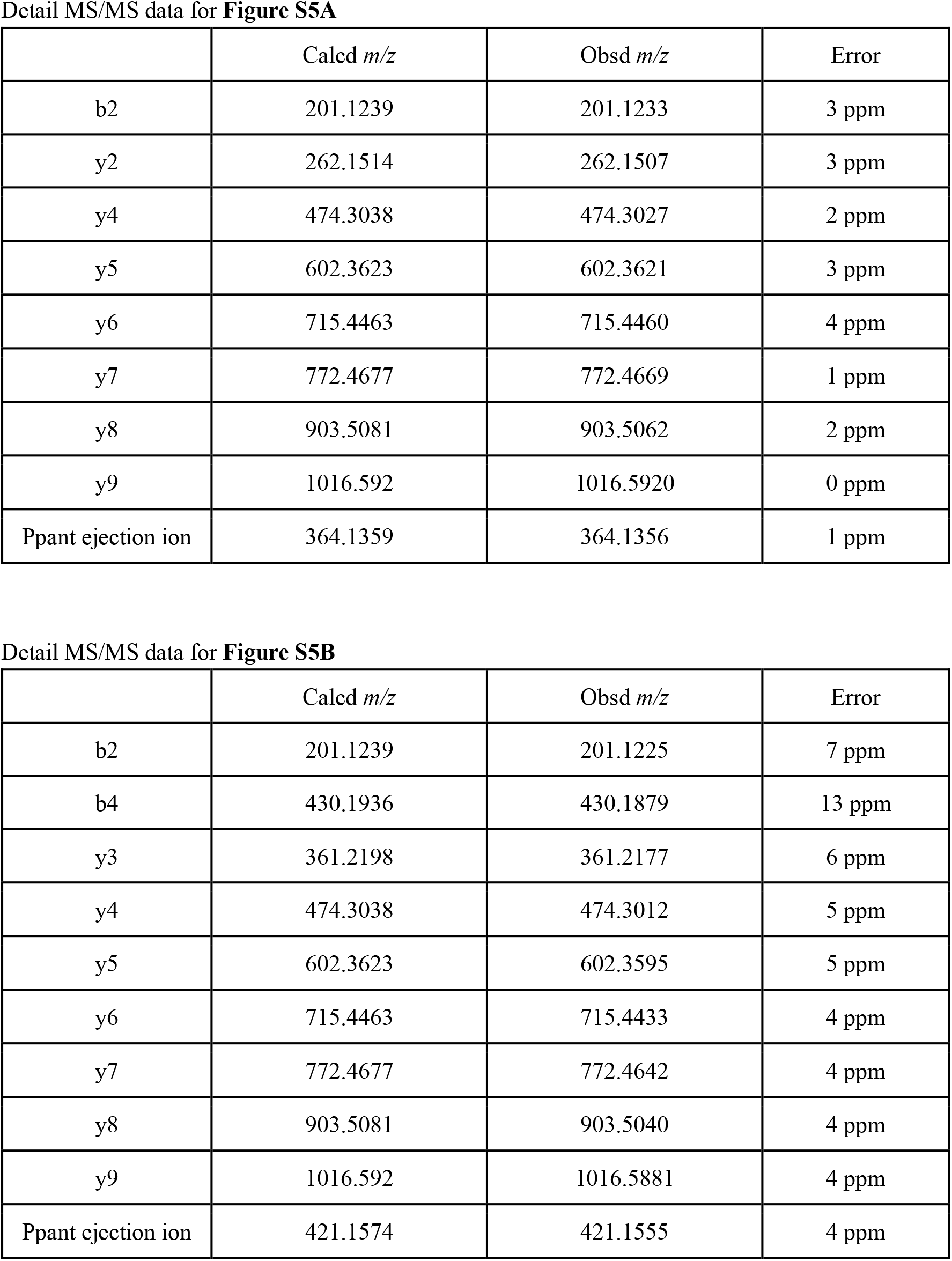
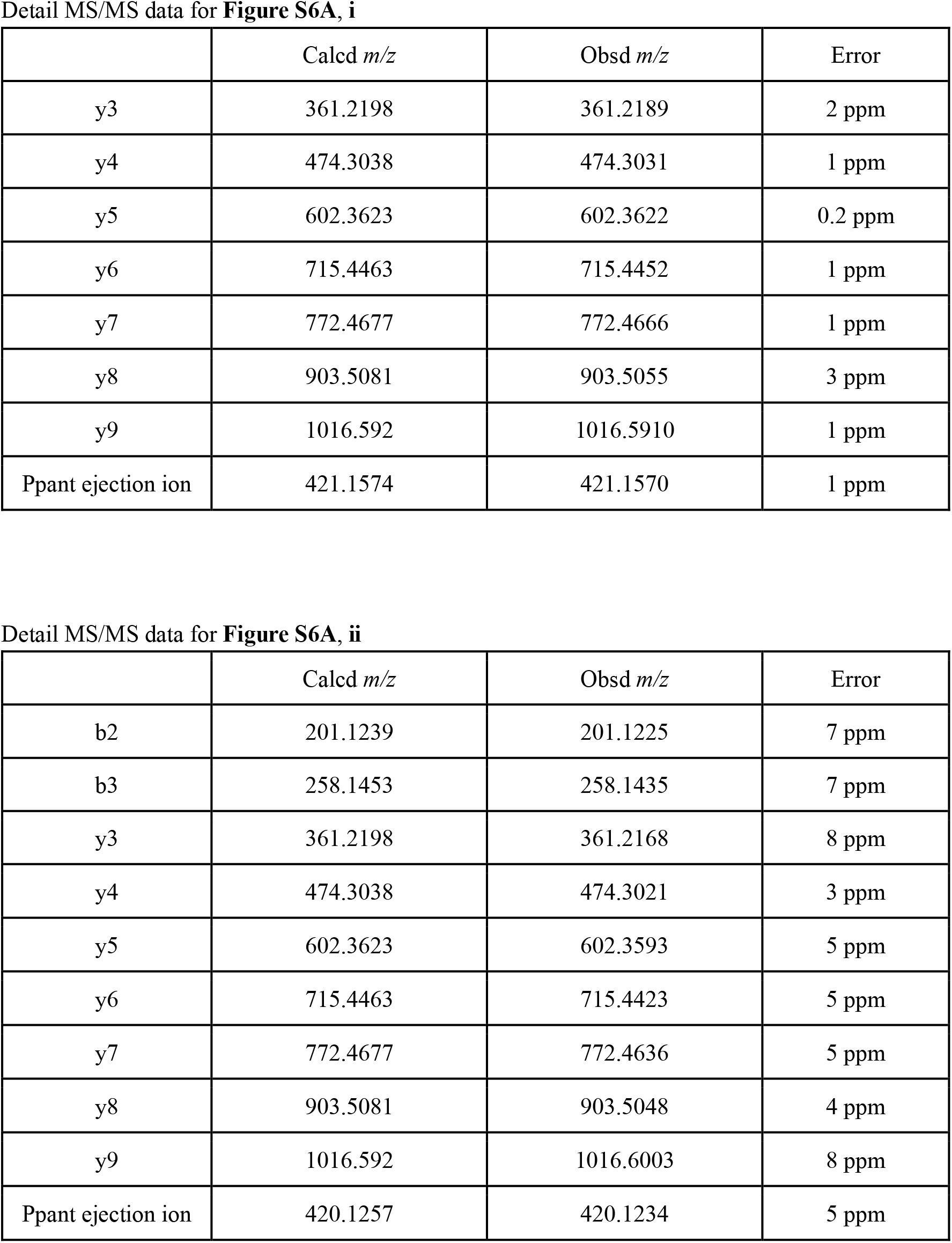
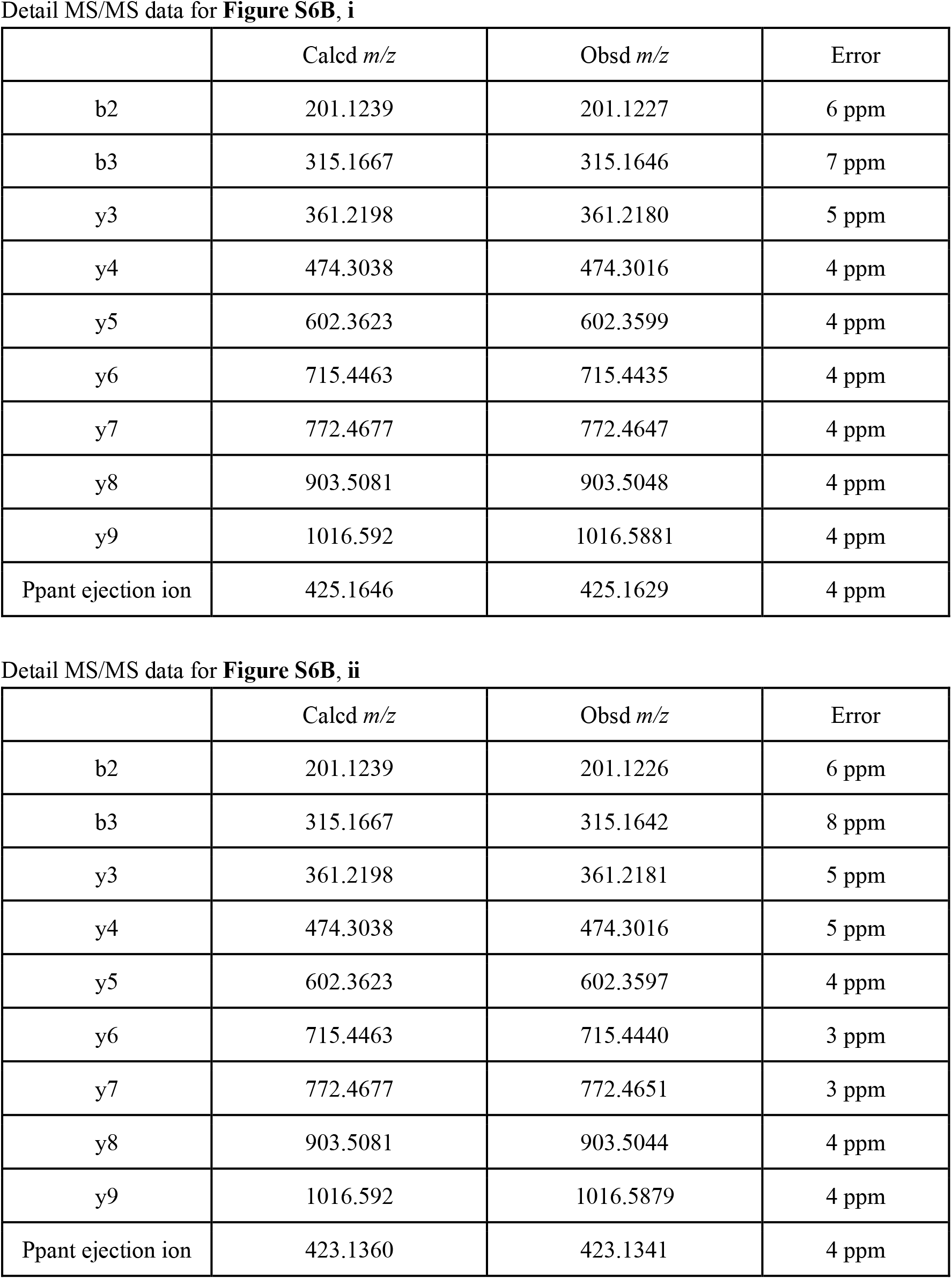
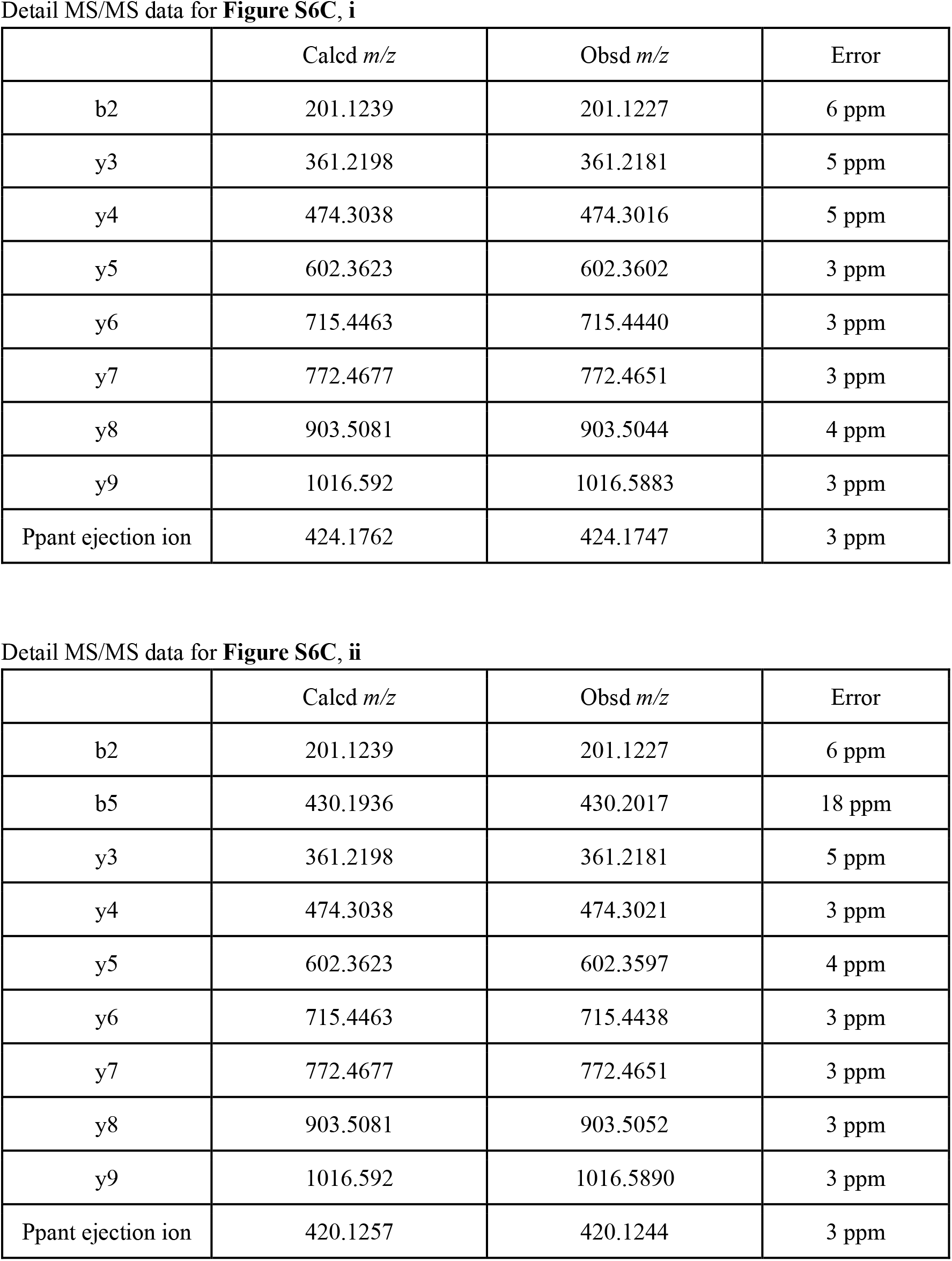
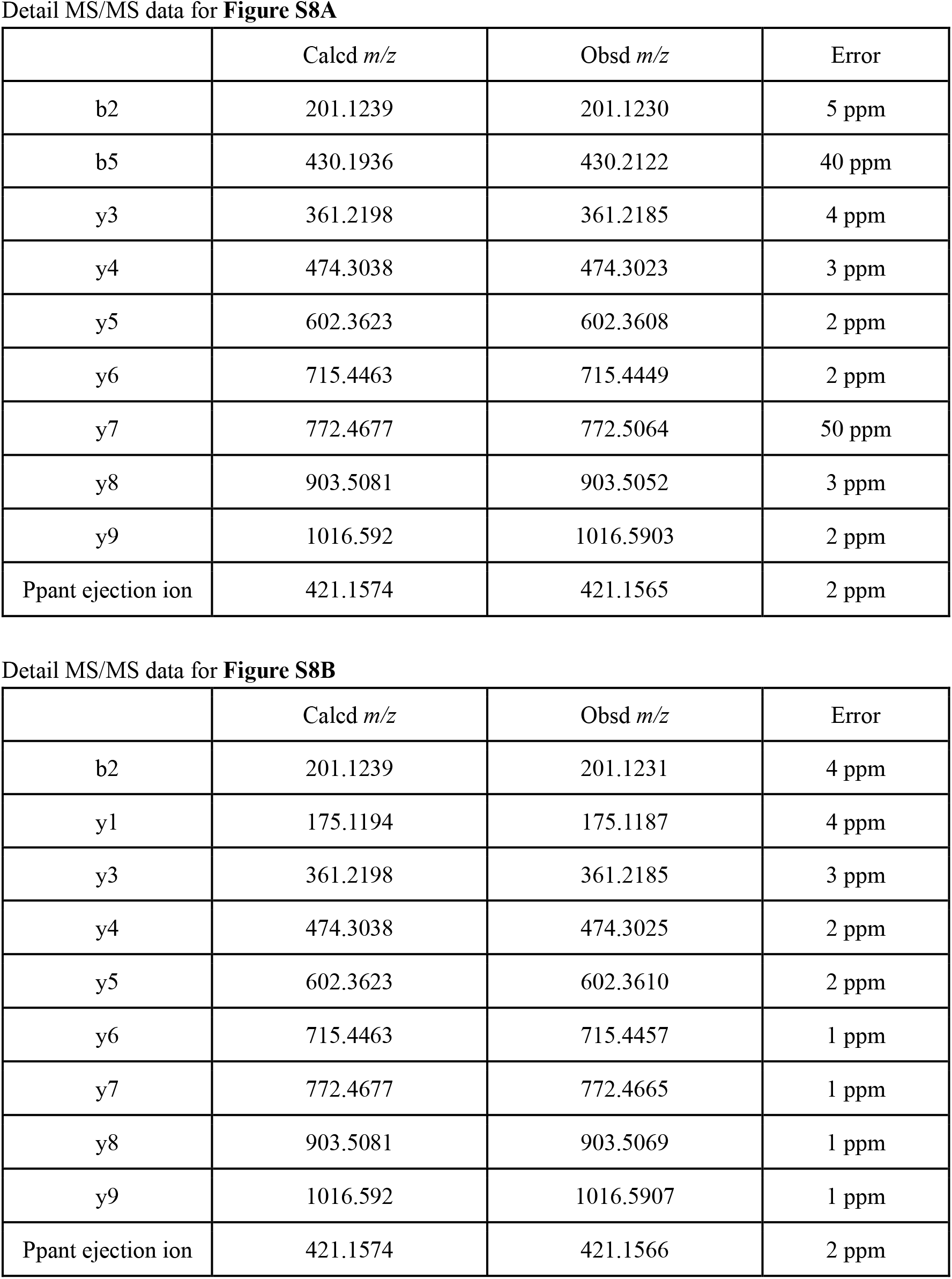
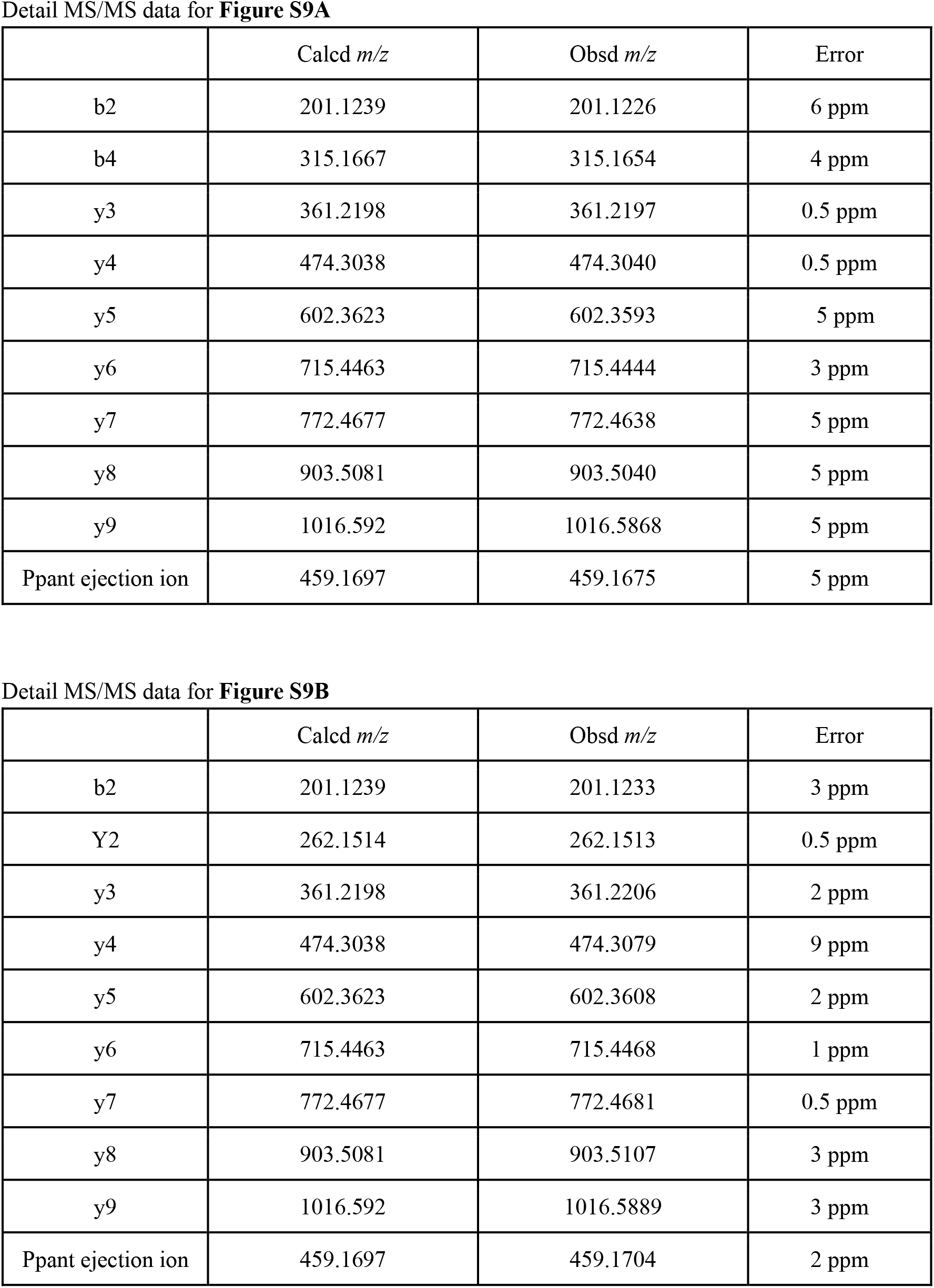
MS/MS data for PCP-aminoacylation on CaeA2.

